# Role of channels in the O_2_ permeability of murine red blood cells I. Stopped-flow and hematological studies

**DOI:** 10.1101/2025.03.05.639948

**Authors:** Pan Zhao, Fraser J. Moss, Rossana Occhipinti, R. Ryan Geyer, Dale E. Huffman, Howard J Meyerson, Walter F Boron

## Abstract

Many have believed that oxygen (O_2_) crosses red blood cell (RBC) membranes by dissolving in lipids that offer a finite resistance to diffusion or, alternatively, no resistance at all. In this first in a series of three interrelated papers, we examine these idea in murine RBCs. In this first paper, analyses of hemoglobin (Hb) absorbance spectra during O_2_ off-loading from mouse RBCs indicate that RBC membranes do indeed offer resistance to O_2_ diffusion, and that the resistance would be far higher if not for the presence of O_2_-permeable channels. Two agents— both excluded from the RBC interior—markedly reduce the rate constant for O_2_ off-loading (*k*_HbO_2__): p-chloromercuribenzenesulfonate (pCMBS) reduces membrane O_2_ permeability (*P*_M,O_2__ by ∼82% (computed from *k*_HbO_2__ in paper #3), and 4,4’-diisothiocyanatostilbene-2,2’-disulfonate (DIDS) by ∼56%. Because neither likely produces these effects via membrane lipids, we examined RBCs from mice genetically deficient in aquaporin-1 (AQP1), the Rh complex (Rh_Cx_ = Rhesus blood group-associated A glycoprotein, RhAG + Rhesus blood group D antigen, RhD), or both. The double knockout (dKO) reduces *P*_M,O_2__ by ∼55%, and pCMBS+dKO, by ∼91%. Proteomic analyses of RBC membranes, flow cytometry, hematology (see paper #2), and mathematical simulations (paper #3) rule out meaningful contributions from other membrane proteins, RBC geometry, or extracellular unconvected fluid (EUF). Our work represents a paradigm shift for O_2_ physiology by identifying the first two animal O_2_ channels, and points to the existence of at least a third, all of which could be subject to physiological regulation and pharmacological intervention.

**Key Points:**

- Some have held that red-blood-cell (RBC) membranes have a finite O_2_ permeability, governed by O_2_ solubility in membrane lipids; others, that membranes offer no resistance whatsoever to O_2_ diffusion.
- The present interdisciplinary study comprises three papers. This first paper describes stopped-flow absorbance spectroscopy in which we examine the rate constant of O_2_ offloading from intact RBCs (*k*_HbO_2__) from wild-type (WT) mice or those lacking AQP1 and/or RhAG, sometimes treated with p-chloromercuribenzenesulfonate (pCMBS) or 4,4’-diisothiocyanatostilbene-2,2’-disulfonate (DIDS).
- The second paper describes RBC morphometry and membrane proteomics. The third introduces a mathematical reaction-diffusion model that generates simulations comporting with physiological data.
- This first paper shows that pCMBS or DIDS treatments, or deletion of AQP1 and/or RhAG, or combinations thereof, substantially reduce *k*_HbO_2__.
- The three papers show that WT RBC membranes offer considerable resistance to O_2_ diffusion. AQP1 (∼22%), Rh (∼36%), and an unknown pCMBS-sensitive protein account for ∼91% of membrane O_2_ permeability.

## Introduction

Central to human life is the inhalation of oxygen (O_2_), its diffusion across the alveolar wall to pulmonary capillary blood plasma, O_2_ diffusion inward across the red blood cell (RBC) membrane and transfer to hemoglobin (Hb; Hb + O_2_ → HbO_2_),^1^ distribution of RBCs by blood flow throughout the body, O_2_ offloading from Hb, and finally O_2_ diffusion outward across the RBC membrane and other barriers to support oxidative metabolism. Critical to understanding this journey of O_2_ from air to mitochondria is the diffusion of O_2_ across various membranes, particularly that of the RBC.

Since the early 1800s, the mechanisms by which gases move across natural and artificial barriers have been the focus of numerous investigations (Graham, 1829, 1833; Mitchell, 1830, 1831a, 1831b, 1833; Fick, 1855; Overton, 1895; see review by Stannett, 1978). For example, Mitchell was the first to recognize that the “facility of transmission” (i.e., permeability) of gases through (primarily synthetic) membranes depends on their “power of infiltration” (i.e., solubility) in the organic molecular barrier (Mitchell, 1831a, 1831b). This was the first statement of “solubility theory” (see Stannett, 1978). Subsequently, Graham, who gave us Graham’s law, hypothesized that gas permeation across barriers depends not only on the ability of the gas to dissolve in the barrier but also to diffuse through it—the first statement of solubility-diffusion theory. In 1879, Wroblewski combined Mitchell’s and Graham’s observations with Henry’s law of gas solubility in liquids (Henry, 1803) and Fick’s law of diffusion (Fick, 1855b, 1855a) to produce our current, simplified mathematical description of the solubility-diffusion theory (Wroblewski, 1879; for reviews, see Stannett, 1978; Cooper *et al*., 2015; Michenkova *et al*., 2021). In the late 1890s, Overton—whose aim was to determine whether biological membranes have a lipoid component—showed that membrane permeability is proportional to the lipophilicity (i.e., the lipid-water partition coefficient) of the permeating molecule (Overton, 1895, 1899). Although Overton’s experiments were far more elegant than those of Mitchell, his conclusion was a restatement of Mitchell’s solubility theory. Over the years, biologists—like Overton, apparently unaware of the work of Mitchell—began to refer to Overton’s observations as “Overton’s rule”, even though Mitchell’s solubility theory predated Overton by more than 60 years, and the “rule” altogether ignores Graham’s more inclusive solubility-diffusion theory that predated Overton by three decades.

Thus, in the early 20^th^ century, many biologists believed that the resistance that membranes offer to the diffusion of gases depends only on the solubility of the gas in the lipid phase of the membrane. Later work on the diffusion of carbon dioxide (CO_2_) through artificial lipid membranes by (Blank & Roughton, 1960), (Gutknecht *et al*., 1977), and (Simon & Gutknecht, 1980) showed that the diffusion component of solubility-diffusion theory is quantitatively far more important than the solubility component (see review by Boron, 2010). Nevertheless, the concept that gases cross membranes merely by dissolving in the lipid phase of the membrane—Overton’s rule— became dogma, at least for most physiologists.

Interestingly, with a smaller group of 20^th^-century physiologists—influenced by the pioneering work of Krogh on O_2_ offloading from RBCs and delivery to the tissues (Krogh, 1919)— a very different dogma evolved. Krogh’s mathematical analysis of the “Krogh cylinder” required the implicit, simplifying assumption (see Kreuzer, 1982) that the various cell membranes, including the RBC membrane, offer no resistance to O_2_ diffusion. Although some notable physiologists argued otherwise (Hartridge & Roughton, 1927; Forster, 1964), Krogh’s implicit assumption became engrained in the literature, and many came to believe that O_2_ crosses RBC membranes simply by dissolving in lipids that offer no resistance whatsoever to diffusion.

Calling the above prevailing views into question were two discoveries, namely, the identification of: (1) membranes with no detectable CO_2_ permeability (Waisbren *et al*., 1994), specifically the apical (i.e., lumen-side membranes) of chief and parietal cells of gastric-glands; and (2) a membrane protein (i.e., aquaporin-1, AQP1; (Preston & Agre, 1991; Preston *et al*., 1992) with an additional role as a CO_2_ channel (Nakhoul *et al*., 1998; Cooper & Boron, 1998). The reagent p-chloromercuribenzenesulfonate (pCMBS), which reacts with the –SH of Cys-189 in the extracellular vestibule of the AQP1 monomeric pore, not only inhibits H_2_O movement through AQP1 (Preston *et al*., 1993), but also blocks a component of CO_2_ traffic through AQP1 (Cooper & Boron, 1998). Moreover, various AQPs exhibit CO_2_ vs. NH_3_ selectivity (Musa-Aziz *et al*., 2009; Geyer *et al*., 2013b). Other work suggested that rhesus (Rh) proteins also can conduct CO_2_ (Endeward *et al*., 2008). These observations revealed that gas transport across membranes must involve more than that predicted by solubility-diffusion theory: the solvation into the lipid bilayer, diffusion through the bilayer, and the desolvation at the opposite side of the bilayer. Work on human RBCs genetically deficient in AQP1 (Endeward *et al*., 2006) or the Rhesus blood group-associated A glycoprotein RhAG (Endeward *et al*., 2008) established that AQP1 and perhaps the Rh complex (Rh_Cx_= RhAG + RhD) as well—if human Rh-null RBCs are indeed biconcave disks— together account for >90% of CO_2_ permeability of RBCs. Moreover, that work established that 4,4’-diisothiocyanatostilbene-2,2’-disulfonate (DIDS), which binds to and can react with the – NH_3_^+/^NH_2_ moiety of Lys residues, blocks a component of CO_2_ traffic through AQP1 and perhaps also Rh_Cx_. Although neither pCMBS nor DIDS is specific, both are valuable tools because (as we show in the present paper) they do not enter RBCs, and they have minimal presence in the lipid phase of the plasma membrane (Vansteveninck *et al*., 1965; Rothstein *et al*., 1973; Cabantchik & Rothstein, 1974). Thus, we hypothesize that—like CO_2_—O_2_ crosses RBC membranes mainly via channels, and that pCMBS or DIDS would block these hypothetical O_2_ channels.

The present study comprises three tightly interwoven papers that aim to determine the extent to which AQP1, Rh_Cx_, and potentially other RBC membrane proteins contribute to the O_2_ permeability of the RBC membrane (*P*_M,O_2__).

In this first (the present) paper, we use stopped-flow (SF) absorbance spectroscopy to determine the rate constant for Hb deoxygenation (*k*_HbO_2__) as we mix oxygenated murine RBCs with a solution containing the O_2_ scavenger sodium dithionite (NDT). This first paper also examines the channel inhibitors pCMBS and DIDS in wild-type (WT) mice, and also RBCs from mice genetically deficient in AQP1, RhAG, or both (i.e., double knockouts, dKO). Recognizing that differences in *k*_HbO_2__ associated with drugs or knockouts (KOs) could reflect changes in parameters other than *P*_M,O_2__, we also examine the kinetics of the reaction HbO_2_ → Hb + O_2_ in free solution as well as hematological parameters, including mean corpuscular volume (MCV) and mean corpuscular hemoglobin (MCH). This first paper also serves as the overall summary of the three-paper project by briefly summarizing, in context, key elements of the other two papers, and providing references to specific locations in those papers.

The second paper (i.e., paper #2; (Moss *et al*., 2025) complements the first by examining RBC morphology at a microscopic scale, including measurements of RBC major diameter (Ø_Major_; i.e., maximum dimension) using imaging flow cytometry (IFC). These data, together with MCV data from the present paper, allow us (in paper #3) to estimate RBC thickness. These IFC data also show that, although nearly all RBCs under control conditions (Ctrl; i.e., no drug treatments) are biconcave disks (BCDs), some are non-biconcave disks (nBCDs); IFC determines nBCD prevalence. In addition, paper #2 includes a proteomic analysis that confirms that the RBC membranes of genetically deficient mice lack only the intended proteins among the 100 proteins with the greatest inferred abundance.

In the third paper (i.e., paper #3; (Occhipinti *et al*., 2025) we report the development of a novel mathematical reaction-diffusion model for O_2_ offloading from an RBC, based on first principles, in which we treat the RBC as a sphere with a diameter that reflects the computed RBC thickness. Together with physical constants, parameter values from the literature, and data from the first two papers, this model generates simulations that are in good agreement with the physiological data, allowing us to extract *P*_M,O_2__ values from *k*_HbO_2__ and other wet-lab data. Our graphical user interface (Huffman *et al*., 2025) expands the power of the model, especially for non-experts.

The overall conclusion of the three papers is that, contrary to the implicit assumption of Krogh, RBC membranes of WT mice under control conditions—that is, WT/Ctrl—offer considerable resistance to O_2_ diffusion, with a computed permeability ∼150 greater than an equivalent layer of water. AQP1 (∼22%) and Rh_Cx_ (∼36%) account for ∼55% of the usual *P*_M,O_2__. Moreover, at least one other pCMBS-sensitive protein is responsible for an additional ∼36%, so that that total protein-mediated component of *P*_M,O_2__ is at least ∼91%. Thus, the work presented in the present three papers changes the way we think of RBC O_2_ handling, and raises the prospect of treating human disease by modulating *P*_M,O_2__ by manipulating membrane proteins^2^.

## Methods

### Ethical approval and animal procedures

The Institutional Animal Care and Use Committee (IACUC) at Case Western Reserve University (CWRU) reviewed and approved all animal procedures.

### Mice

We originally obtained as *Aqp1*+/– mice—with a targeted disruption (replacement of the last 24 bp of exon 1 and 1.3 kb of intron 1 by the 1.8-kb PolII-neo selectable marker) of the *Aqp1* gene (Ma *et al*., 1998)—on a “C57BL/6” background (Kelmenson, 2016), as a generous gift of Alan Verkman. The *Aqp1*+/+ mice obtained from the initial crosses were backcrossed for >20 generations by our laboratory to define our lab-standard WT strain, C57BL/6_Case_.

We backcrossed *Aqp1*+/– mice into our C57BL/6_Case_ WT mice for >18 generations, and confirmed genotypes. Similarly, we backcrossed *Rhag*+/– mice (provided by Jean-Pierre Cartron as *Rhag*+/– mice on a “C57BL/6” background), with a targeted disruption of *Rhag* (Goossens *et al*., 2010), with C57BL/6_Case_ mice and backcrossed into C57BL/6_Case_ mice for at least 7 generations. From *Aqp1*–/– and *Rhag*–/– mice, we generated dKO (i.e., *Aqp1*–/–*Rhag*–/–) mice, and confirmed genotypes as described below. All three KO strains we routinely confirmed by genotyping (see <u>Genotyping</u>).

We contracted with TransnetYX, Inc. (Cordova, TN) to perform Genetic Background analysis on our four mouse strains, all on a C57BL/6_Case_ background. TransnetYX examined all ∼10,000 single-nucleotide polymorphisms (SNPs) in their catalog, SNPs from >200 strains of mice. These included 156 SNPs specific for C57BL/6J (from Jackson Laboratory), 31 SNPs for C57BL/6N (from the National Institutes of Health), and 21 SNPs for C57BL/6Group (characteristic of other C57BL/6 strains). Table 1 summarizes the results of this analysis, which shows that our C57BL/6_Case_ mice are on average ∼19.6% C57BL/6J, ∼20.6% C57BL/6N, ∼25.6% C57BL/6Group, for a total BL/6 character of ∼65.8%.

**Table 1.** Genetic background analysis of C57BL/6_Case_ mice, based on ∼10,000 single-nucleotide polymorphisms*.

| Character<br>Genotype | C57BL/6J | C57BL/6N | C57BL/6Group | Total |
| --- | --- | --- | --- | --- |
| *Aqp1*<sup>+/+</sup>*Rhag*<sup>+/+</sup> | 17.60 ± 0.89 % | 18.85 ± 1.35 % | 23.80 ± 0.00 % | 60.25 % |
| *Aqp1*<sup>-/-</sup> | 16.57 ± 1.49 % | 19.40 ± 0.00 % | 28.60 ± 0.00 % | 64.57 % |
| *Rhag*<sup>-/-</sup> | 22.77 ± 1.43 % | 19.40 ± 0.00 % | 28.60 ± 0.00 % | 70.77 % |
| *Aqp1*<sup>-/-</sup>*Rhag*<sup>-/-</sup> | 21.38 ± 3.20 % | 24.73 ± 2.61 % | 21.42 ± 5.85 % | 67.53 % |
| Average | 19.58 % | 20.60 % | 25.60 % | 65.78 % |

\*N is 6 for each genotype, values represent mean ± SD.

Mice were allowed access to food and water ad libitum. In experiments, we used mice of both sexes (8 to 19 weeks old). Not shown are data indicating that no changes in kinetic or hematological parameters occur up to 6 months of age for all genotypes.

### Genotyping

WT vs. *Aqp1*+/– vs. *Aqp1*–/– are genotyped by real-time PCR (performed by TransnetYX, Inc.). For detecting the *Aqp1*– allele, the forward primer is 5′-CAC CTC GGT AGC AAC TCG AA-3′, the reporter primer 1 is 5′-CTC CTC GAA GGA ATT C-3′, and the reverse primer is 5′- GGG CTC GAG ATC CAC TAG ACT-3′. For detecting WT, the forward primer is 5′-CTT AGG CTT AGG GTG TCT GTA GGA-3′, the reporter primer 1 is 5′-CTG GCT CCA TTT GCC-3′, and the reverse primer is 5′-CTT CAT CCT TCT CTC TGT TCT TCA GT-3′.

WT vs. *Rhag*+/– vs. *Rhag*–/– mice are genotyped by real-time PCR (performed by TransnetYX, Inc.) to determine the presence of a *Rhag*–allele. For detecting the KO allele, the forward primer is 5′-TGC TGC TAT GAC CAT TGG AA-3′, and the reverse primer is 5′-ATT GCA TCG CAT TGT CTG AG-3′. The KO PCR results 196-bp band.

If we detect a KO allele in a mouse, we must differentiate between a *Rhag*+/– and *Rhag*–/– mouse by attempting to detect the WT allele. We perform this assay in house because the PCR product is long (∼8 kb). We prepare highly purified genomic DNA from tail tissue using the DNeasy Blood and Tissue Kit (Qiagen, Germantown, MD), and perform *Rhag*-WT genotyping. Primer “*Rhag*-WT 3F” (5′-TGA AAT GTT TAA TGC TCA TCG ACC AC-3′) is designed to anneal to the end of Intron 1–2 of *Rhag*, and Primer “*Rhag*-WT 3R” (5′-GAG TGA CTT CA CCT TCT TAT CTT CTA AG-3′), to the middle of intron 6–7. The *Rhag*-WT PCR product is ∼8 kb, as noted above, using the Qiagen Long Range (LR) PCR Kit. Presence of the WT-PCR product confirms a *Rhag*+/– genotype for any mouse reported to be positive for *Rhag*–/– by TransnetYX. Absence of a WT-PCR product confirms the *Rhag*–/– genotype.

### Physiological solutions

Table 2 shows the compositions of all solutions used in the present study. We made pH measurements—either at room temperature (RT) or at 10°C—using a portable pH meter (model A121 Orion Star, Thermo Fischer Scientific (TFS), Waltham, MA) fitted with a pH electrode (Ross Sure-Flow combination pH Electrode, TFS). Beakers containing pH calibration buffers (pH at 6, 8 and 10; TFS), physiological solutions to be titrated, and the pH electrode in its storage solution (Beckman Coulter, Inc. Brea, CA) were equilibrated at either RT or 10°C, as appropriate. For work at 10°C, we used a refrigerated, constant-temperature, shaker water bath (model RWB 3220, TFS) and submersible Telesystem magnetic stirrers (TFS). We adjusted pH with NaOH (5 M) or HCl (5 M). Osmolality was measured using a vapor pressure osmometer (Vapro 5520; Wescor, Inc., Logan, UT) and, as necessary, adjusted upward by addition of NaCl.

**Table 2.**
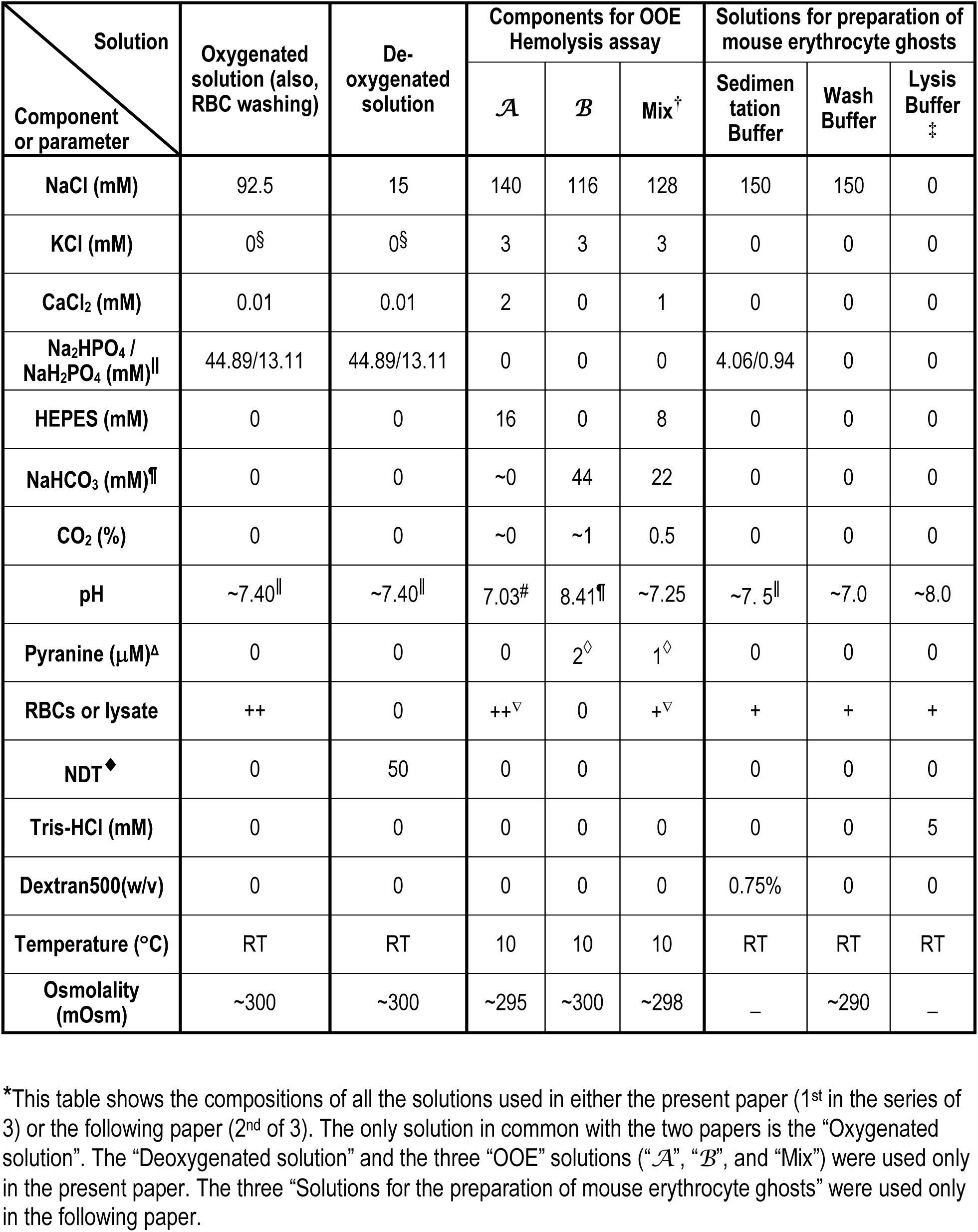
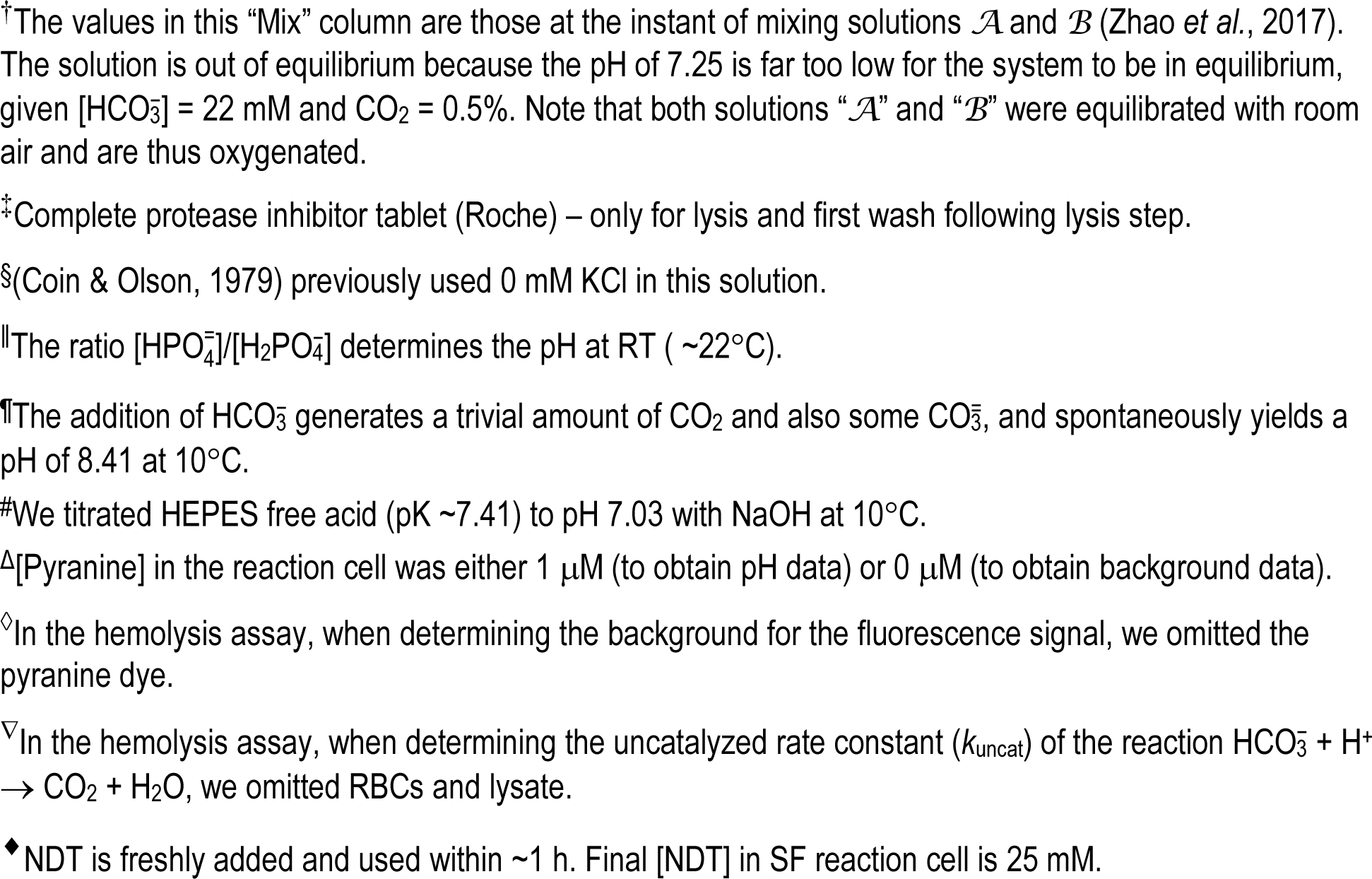
Physiological solutions*.

| Solution<br>Component<br>or parameter | Oxygenated<br>solution (also,<br>RBC washing) | De-<br>oxygenated<br>solution | Components for OOE<br>Hemolysis assay |  |  | Solutions for preparation of<br>mouse erythrocyte ghosts |  |  |
| --- | --- | --- | --- | --- | --- | --- | --- | --- |
| | | | $\mathcal{A}$ | $\mathcal{B}$ | Mix<sup>†</sup> | Sedimen-<br>tation<br>Buffer | Wash<br>Buffer | Lysis<br>Buffer<sup>‡</sup> |
| NaCl (mM) | 92.5 | 15 | 140 | 116 | 128 | 150 | 150 | 0 |
| KCl (mM) | 0<sup>§</sup> | 0<sup>§</sup> | 3 | 3 | 3 | 0 | 0 | 0 |
| CaCl<sub>2</sub> (mM) | 0.01 | 0.01 | 2 | 0 | 1 | 0 | 0 | 0 |
| Na<sub>2</sub> HPO<sub>4</sub> /<br>NaH<sub>2</sub> PO<sub>4</sub> (mM) | 44.89/13.11 | 44.89/13.11 | 0 | 0 | 0 | 4.06/0.94 | 0 | 0 |
| HEPES (mM) | 0 | 0 | 16 | 0 | 8 | 0 | 0 | 0 |
| NaHCO<sub>3</sub> (mM) | 0 | 0 | ~0 | 44 | 22 | 0 | 0 | 0 |
| CO<sub>2</sub> (%) | 0 | 0 | ~0 | ~1 | 0.5 | 0 | 0 | 0 |
| pH | ~7.40 | ~7.40 | 7.03<sup>#</sup> | 8.41 | ~7.25 | ~7.5 | ~7.0 | ~8.0 |
| Pyranine (μM)<sup>Δ</sup> | 0 | 0 | 0 | 2<sup>◇</sup> | 1<sup>◇</sup> | 0 | 0 | 0 |
| RBCs or lysate | ++ | 0 | ++<sup>▽</sup> | 0 | +<sup>▽</sup> | + | + | + |
| NDT<sup>◆</sup> | 0 | 50 | 0 | 0 |  | 0 | 0 | 0 |
| Tris-HCl (mM) | 0 | 0 | 0 | 0 | 0 | 0 | 0 | 5 |
| Dextran500(w/v) | 0 | 0 | 0 | 0 | 0 | 0.75% | 0 | 0 |
| Temperature (°C) | RT | RT | 10 | 10 | 10 | RT | RT | RT |
| Osmolality<br>(mOsm) | ~300 | ~300 | ~295 | ~300 | ~298 | — | ~290 | — |

\*This table shows the compositions of all the solutions used in either the present paper (1<sup>st</sup> in the series of 3) or the following paper (2<sup>nd</sup> of 3). The only solution in common with the two papers is the “Oxygenated solution”. The “Deoxygenated solution” and the three “OOE” solutions (“ $\mathcal{A}$ ”, “ $\mathcal{B}$ ”, and “Mix”) were used only in the present paper. The three “Solutions for the preparation of mouse erythrocyte ghosts” were used only in the following paper.
<sup>†</sup>The values in this “Mix” column are those at the instant of mixing solutions $\mathcal{A}$ and $\mathcal{B}$ (Zhao *et al.*, 2017). The solution is out of equilibrium because the pH of 7.25 is far too low for the system to be in equilibrium, given $[\text{HCO}_3^-] = 22 \text{ mM}$ and $\text{CO}_2 = 0.5\%$ . Note that both solutions “ $\mathcal{A}$ ” and “ $\mathcal{B}$ ” were equilibrated with room air and are thus oxygenated.
<sup>‡</sup>Complete protease inhibitor tablet (Roche) – only for lysis and first wash following lysis step.
<sup>§</sup>(Coin & Olson, 1979) previously used 0 mM KCl in this solution.
<sup>||</sup>The ratio $[\text{HPO}_4^{2-}]/[\text{H}_2\text{PO}_4^-]$ determines the pH at RT ( $\sim 22^\circ\text{C}$ ).
<sup>¶</sup>The addition of $\text{HCO}_3^-$ generates a trivial amount of $\text{CO}_2$ and also some $\text{CO}_3^{2-}$ , and spontaneously yields a pH of 8.41 at $10^\circ\text{C}$ .
<sup>#</sup>We titrated HEPES free acid ( $\text{pK} \sim 7.41$ ) to pH 7.03 with NaOH at $10^\circ\text{C}$ .
<sup>Δ</sup>[Pyranine] in the reaction cell was either $1 \mu\text{M}$ (to obtain pH data) or $0 \mu\text{M}$ (to obtain background data).
<sup>◇</sup>In the hemolysis assay, when determining the background for the fluorescence signal, we omitted the pyranine dye.
<sup>∇</sup>In the hemolysis assay, when determining the uncatalyzed rate constant ( $k_{\text{uncat}}$ ) of the reaction $\text{HCO}_3^- + \text{H}^+ \rightarrow \text{CO}_2 + \text{H}_2\text{O}$ , we omitted RBCs and lysate.
<sup>◆</sup>NDT is freshly added and used within $\sim 1 \text{ h}$ . Final [NDT] in SF reaction cell is 25 mM.

### Inhibitors

Stock solutions of 2 mM pCMBS (catalog no. C367750; Toronto Research Chemicals, Toronto, ON, Canada) and 2 mM DIDS (catalog no. 1226913; Invitrogen, Eugene, OR) were freshly prepared (generally 1–2 h before experiments) by dissolving the agents directly into either our “Oxygenated solution” or solution “*A*” (see Table 2), taking care to shield the solutions from light.

### Preparation of RBCs

**Collection of blood from mice.** We collected fresh blood from WT or KO mice for seven assays: (1) SF, (2) still microphotography, (3) microvideography, (4) flow cytometry, (5) automated hematology, (6) blood smears, and (7) proteomics/mass spectrometry. Early on (only for SF studies), we collected blood using the cardiac-puncture method (Parasuraman *et al*., 2010). Later (for the remaining SF and all other studies), so that we could return to the same mouse for more than one sample of blood, we used the submandibular-bleed method (Golde *et al*., 2005). For cardiac puncture, we used a 1-ml syringe (Becton, Dickinson and Co., Franklin Lakes, New Jersey) and attached 23-gauge PrecisionGlide needle (Becton, Dickinson and Co). For bleeds of the submandibular vein, we used a 3-, 4-, or 5-mm point-length sterile animal lancet (MEDIpoint, Inc., Mineola, NY), and took blood samples (∼200 μl) from the same mouse no sooner than 72 h from a previous blood sample, and on the contralateral side.

**Processing of RBCs for assays 1–4.** For the first four assays (SF, still microphotography, microvideography, flow cytometry), we collected the blood into 1.7-ml microcentrifuge tubes (Laboratory Products Sales Inc. Rochester, N.Y.) that we had previously rinsed with 0.1% sodium heparin (H4784, Sigma-Aldrich, St. Louis, MO). We then centrifuged in a Microfuge 16 Microcentrifuge (Beckman Coulter, Inc.) at 600 × g for 10 min, and aspirated and discarded the resulting supernatant and buffy coat. To remove residual extracellular Hb, we resuspended the pelleted RBCs in our oxygenated solution (see Table 2) to a hematocrit (Hct) of 5% to 10%, centrifuged at 600 × g for 5 min. After 4 such washes, we resuspended RBCs in our oxygenated solution to a final Hct of 25% to 30%, and maintained on ice (for that day) for experiments. At this point, we continued with the processing of RBCs for SF studies (the present paper; see <u>below</u> ^3^) or for the three studies in paper #2, namely, <u>still microphotography</u>, <u>microvideography</u>, or <u>flow-cytometry</u>^4^.

**Processing of RBCs for automated hematology.** For automated hematological studies (see <u>below</u>^5^), we collected whole blood into 20-μl plastic Boule MPA Micro pipettes (EDTA-K2, Boule Medical AB, Stockholm, Sweden).

**Processing of RBCs for blood smears.** For <u>blood smears</u>^6^, we collected whole blood into 2-ml K2E K2EDTA VACUETTE® tubes (Grelner Bio-One North America Inc, Monroe, NC).

**Processing of RBCs for proteomics.** For <u>Proteomic analysis</u>^7^, we collected whole blood into 1.7-ml heparinized microcentrifuge tubes (see <u>above</u>^8^).

**Processing of RBCs for stopped-flow studies.** For SF studies, after collection of the blood (see <u>above</u>^9^), we computed [Hb] in a suspension of intact RBCs from the Hb absorbance measured on a UV/Vis spectrophotometer (model 730 Life Science, Beckman Coulter, Inc.), using an equation described previously (Zhao *et al*., 2017).

As necessary, we produced an RBC lysate by osmotic lysis of 20 μl of freshly prepared, packed mouse RBCs in a 1:8∼1:10 dilution of Milli-Q H_2_O (Milli-Q® Integral Water Purification System, EMD Millipore Corporation, Billerica, MA), followed by centrifugation at 15000 × g for 5 min in a Microfuge 16 Microcentrifuge, at RT. We transferred the supernatant to a clean 1.7-ml microcentrifuge tube for a spectroscopic determination of free [Hb], as described previously (Zhao *et al*., 2017).

For all stopped-flow experiments—on intact RBCs or lysates or mixtures thereof—we dilute the material to a [Hb] of 5 μM before introduction into the SF device. After mixing in the SF reaction cell, [Hb] falls to a final values of 2.5 μM. In the case of deoxygenation assays (e.g., in Results, see Figure 3 and Figure 4), the diluent was our oxygenated solution (see Table 2). In the case of carbonic-anhydrase assays (e.g., in Results, see summary in Figure 6; see Zhao *et al*., 2017), the diluent was solution “*A*” (see Table 2), which is also oxygenated.

**Use of inhibitors in stopped-flow studies.** In all inhibitor studies, except those in which we studied short-term exposures to DIDS (introduced next), we pre-incubated samples (either ostensibly intact RBCs or a hemolysate) with the drug before introduction into the SF device:

1. pCMBS (1 mM): 15-min preincubation. For:

a. ostensibly intact RBCs, see Figure 3*C* and *D*, as well as filled bars in Figure 3*E* and *F*;
b. hemolysate, see open bars in Figure 3*F*.
2. DIDS (200 μM): 60-min preincubation. For:

a. ostensibly intact RBCs, see Figure 4*C* and *D* as well as filled bars in Figure 4*E* and *F*;
b. hemolysate, see open bars in Figure 4*F* as well as filled gray-blue bar in Figure 8*A*).

The next step, for both drugs, was to dilute the material in our oxygenated solution (see Table 2) to achieve a [Hb] of 5 μM plus either 1 mM pCMBS or 200 μM DIDS, added from a stock solution (see “Inhibitors”, <u>above</u>^10^). Finally, we performed stopped-flow studies by mixing the aforementioned oxygenated solution with the deoxygenated solution (see Table 2) to obtain *k*_HbO_2__.

In a few studies, we assessed the effect of millisecond-to-second DIDS exposures on Hb absorbance spectra of hemolysates (e.g., in Results, see Figure 8*B*, gray-blue open bar). Here, we did not preincubated the hemolysate in DIDS. Rather, we placed (1) the lysate in the oxygenated solution and (2) the DIDS in the deoxygenated solution (see Table 2). Thus, the DIDS did not contact the Hb in the lysate until reaching the SF reaction cell as we determined *k*_HbO_2__.

### Stopped-flow absorbance spectroscopy (workflow #1)

The flow chart in Figure 1 describes the series of steps—starting with the time course of RBC absorbance spectra (workflow: step #1) introduced in this section—to arrive at refined estimates for *k*_HbO_2__ and the associated *P*_M,O_2__.

**Figure 1.**
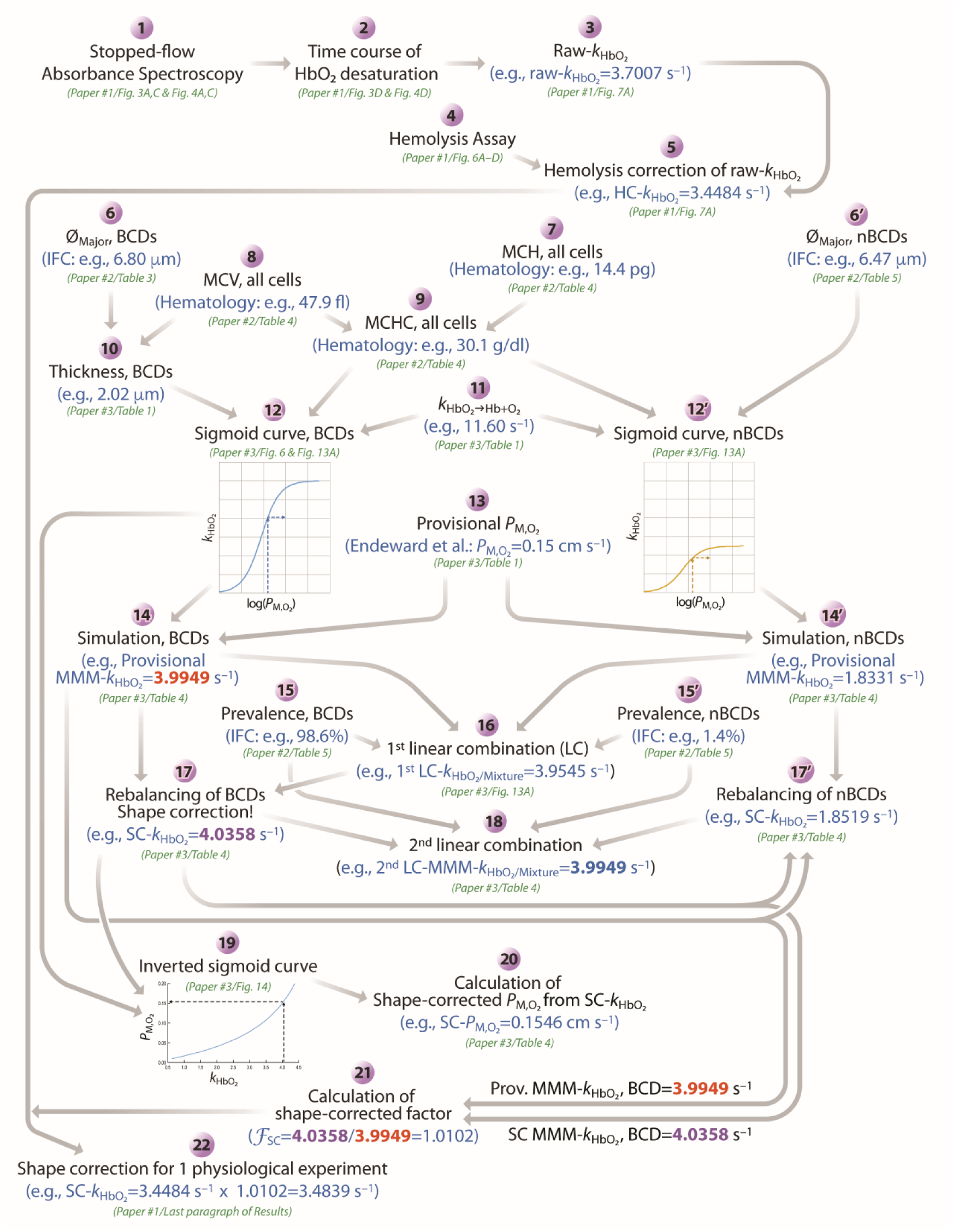
Workflow to obtain hemolysis-corrected and shape-corrected values of ***P*_M,O_2__**. The numerals 1 through 22 indicate the steps summarized in the present paper—and presented in detail, as appropriate, in the present paper (<u>paper #1</u>), <u>paper #2,</u> or <u>paper #3</u>—to arrive, first, at a hemolysis-corrected (HC) *k*_HbO_2__ and, ultimately, at a shape-corrected (SC) *k*_HbO_2__. At each step, we provide example values, if possible, and example figure/table numbers referenced to the present paper, <u>paper #2</u>, or <u>paper #3</u>. We repeat workflow steps 1–5 for each mouse, ultimately arriving at an HC-*k*_HbO_2__ value for each. Step 6 applies to biconcave disks (BCDs), whereas step 6′ applies to non-biconcave disks (nBCDs). The same is true for steps 12 vs. 12′, 14 vs. 14′, 15 vs. 15′, and for 17 vs. 17′. *F*_SC_, shape-correction factor; Hb, hemoglobin; IFC, imaging flow cytometry; MCH, mean corpuscular hemoglobin; *k*_HbO_2__, rate constant for deoxygenation of HbO_2_ within intact RBCs; *k*_HbO_2_→Hb+O_2__, rate constant for deoxygenation of HbO_2_ in free solution; LC, linear combination; MCH, mean corpuscular hemoglobin; MCHC, mean corpuscular hemoglobin concentration; MCV, mean corpuscular volume; MMM-k_HbO_2__, macroscopic mathematically modeled rate constant for deoxygenation of intracellular HbO_2_; *P*_M,O_2__, membrane permeability to O_2_; Prov., provisional; Ø_Major_, major diameter (of BCD or nBCD); SC-k_HbO_2__, shape-corrected rate constant for deoxygenation of intracellular HbO_2_.

We use an SX-20 stopped-flow apparatus (Applied Photophysics, Leatherhead, UK) in absorbance spectroscopy mode to determine the rate constant of hemoglobin deoxygenation. We load RBCs or RBC/ hemolysate mixtures or pure hemolysate ([Hb] = 5 μM, Hct ≅ 0.37%), suspended in our oxygenated solution (see Table 2), into one syringe of the SF apparatus. We load the deoxygenated solution containing the O_2_ scavenger NDT (50 mM; SIGMA-ALDRICH, Co, St, Louis, MO) into the other syringe. We record absorbance (*A*) in a 20-μl SF reaction cell (2-mm path length) at 10°C, over the portion of the Hb spectrum with highest Signal/Noise (wavelength [λ]: 410–450 nm, at 5 nm intervals), sampling at 10-ms intervals for 4000 ms. For each new set of loaded samples, we begin by performing ≥6 “shots” to ensure that the new solutions are loaded into the SF reaction cell, and then sequentially acquire *A*_λ_ vs. time during 9 separate shots at λ = 410, 415 … 450 nm—this constitutes one data set. We compile such data set into 3-D graphs (e.g., in Results, see *A*_λ_ vs. λ vs. time plots in Figure 3*A* and *C* as well as Figure 4*A* and *C*). We obtain at least 1 such data set (see criteria in next section)—but typically 2 to 4—for each set of experimental conditions, and then average these *k*_HbO_2__ values to obtain the raw-*k*_HbO_2__ value used in further analyses.

The above sequence is step #1 in Figure 1.

### Calculation of time course of HbO_2_ desaturation (workflow #2)

The time course of the decay HbO_2_ → Hb + O_2_ does not follow a perfect exponential function—nor should it according to our mathematical modeling (e.g., see figure 4 in paper #3). We therefore developed an unbiased approach^11^ for computing the quasi-rate constant *k*_HbO_2__, based on the time (*t*_37_) required for O_2_ saturation of Hb (HbSat) to fall to 1/e of its initial value. We proceed in the following sequence:

**Preprocess data.** Eliminate noisy absorbance data—which we standardize as the first 9 points (0.010 – 0.090 s)—and translate the time axis so that raw data at absolute times 0.100 – 4.000 s now covers the relative time range 0.000 – 3.900 s. We also eliminate data at 3 wavelengths (420, 445, 450 nm) that are near isosbestic points of the Hb/HbO_2_ absorbance spectra because, at these wavelengths, absorbance values do not change consistently during an experiment. The result is a data set describing *A*_λ_ vs. relative time (from 0.000 to 3.900 s) at each of 6 wavelengths (λ = 410, 415, 425, 430, 435, and 440 nm). We accept a data set only if each of the 6 records of *A*_λ_ vs. time approaches an asymptote.

**Obtain data parameters.** For each of 6 plots of *A*_λ_ vs. time, obtain the minimum (*A*_λ_,_min_) and maximum values (*A*_λ_,_max_), as well as the difference Δ*A*_λ_ = *A*_λ_,_max_ – *A*_λ_,_min_ (i.e,. magnitude of the absorbance change). The 6 values of Δ*A*_λ_ become weighting factors, applied two paragraphs below.

**Normalize.** Perform a vertical flip of plots of *A*_λ_ vs. time that were originally concave downwards (these are λ = 425, 430, 435, 440 nm), so that all plots are now concave upwards. Vertically translate each plot of *A*_λ_ vs. time so that the minimum value is zero. Normalize each plot of *A*_λ_ vs. time so that *A*_λ_,_max_ is now unity.

**Apply weighting factors.** Normalize the 6 weighting factors (i.e., Δ*A*_λ_) so that they sum to unity. Multiply each plot of *A*_λ_ vs. time by its weighting factor. At each time point, sum the 6 weighted *A*_λ_ values to obtain relative HbSat. The resulting plot of relative HbSat thus falls from unity to zero as relative time increases from 0.000 s to 3.900 s. Note that—because the RBCs or Hb are equilibrated in room air before being loaded into the SF device—the actual HbSat falls from a value of ∼98% when the RBCs first enter the SF reaction cell at absolute time 0.000 s, and has already fallen to some extent by the time of our first accepted sample at absolute time 0.100 s, and then continues to fall to zero during the remaining 3.900 s of the “shot.”

The above sequence is step #2 in Figure 1.

In Results, Figure 3*D*, Figure 4*D*, as well as Figure 5*A, C*, and *E* show examples of the final product of such analyses, that is, plots of HbSat vs. time.

**Table 3.** Summary of raw, hemolysis-corrected, and shape-corrected *k*_HbO_2__ values.

| Condition<br>Parameter | Ctrl | pCMBS | *P*-value | Ctrl | DIDS | *P*-value |
| --- | --- | --- | --- | --- | --- | --- |
| Raw-$k_{\text{HbO}_2}$ ($\text{s}^{-1}$) | 3.74 ± 0.31 | 1.81 ± 0.39 | 5.89$\times 10^{-8}$ | 3.56 ± 0.30 | 2.38 ± 0.12 | 1.08$\times 10^{-6}$ |
| HC-$k_{\text{HbO}_2}$ ($\text{s}^{-1}$) | 3.42 ± 0.25 | 1.26 ± 0.19 | 4.97$\times 10^{-7}$ | 3.21 ± 0.27 | 1.65 ± 0.19 | 3.84$\times 10^{-8}$ |
| SC-$k_{\text{HbO}_2}$ ($\text{s}^{-1}$) | 3.45 ± 0.25 | 1.34 ± 0.20 | 7.03$\times 10^{-7}$ | 3.24 ± 0.27 | 2.23 ± 0.26 | 1.07$\times 10^{-6}$ |
| N | 8 |  |  | 8 |  |  |

**B. $k_{\text{HbO}_2}$ of RBCs from WT and knockout mice**
| Condition<br>Parameter | WT | *Aqp1*<sup>-/-</sup> | *Rhag*<sup>-/-</sup> | *Aqp1*<sup>-/-</sup>*Rhag*<sup>-/-</sup> |
| --- | --- | --- | --- | --- |
| Raw-$k_{\text{HbO}_2}$ ($\text{s}^{-1}$) | 4.22 ± 0.51 | 3.81 ± 0.32 | 3.56 ± 0.21 | 2.99 ± 0.23 |
| HC-$k_{\text{HbO}_2}$ ($\text{s}^{-1}$) | 4.05 ± 0.48 | 3.68 ± 0.29 | 3.37 ± 0.15 | 2.80 ± 0.23 |
| SC-$k_{\text{HbO}_2}$ ($\text{s}^{-1}$) | 4.09 ± 0.49 | 3.73 ± 0.30 | 3.41 ± 0.15 | 2.84 ± 0.24 |
| N | 10 | 12 | 10 | 11 |

**C. $k_{\text{HbO}_2}$ of RBCs from double-knockout mice, ± inhibitors**
| Condition<br>Parameter | Ctrl | pCMBS | *P* value | Ctrl | DIDS | *P*-value |
| --- | --- | --- | --- | --- | --- | --- |
| Raw-$k_{\text{HbO}_2}$ ($\text{s}^{-1}$) | 2.90 ± 0.13 | 1.37 ± 0.06 | 3.47$\times 10^{-10}$ | 2.75 ± 0.30 | 2.20 ± 0.12 | 3.24$\times 10^{-4}$ |
| HC-$k_{\text{HbO}_2}$ ($\text{s}^{-1}$) | 2.59 ± 0.18 | 0.81 ± 0.23 | 3.99$\times 10^{-9}$ | 2.48 ± 0.33 | 1.47 ± 0.23 | 3.44$\times 10^{-5}$ |
| SC-$k_{\text{HbO}_2}$ ($\text{s}^{-1}$) | 2.63 ± 0.18 | 0.84 ± 0.24 | 4.96$\times 10^{-9}$ | 2.52 ± 0.34 | 1.70 ± 0.27 | 2.30$\times 10^{-4}$ |
| N | 8 |  |  | 9 |  |  |

Values represent mean ± SD. For each comparison we use paired two-tailed t-test.

**Table 4.**
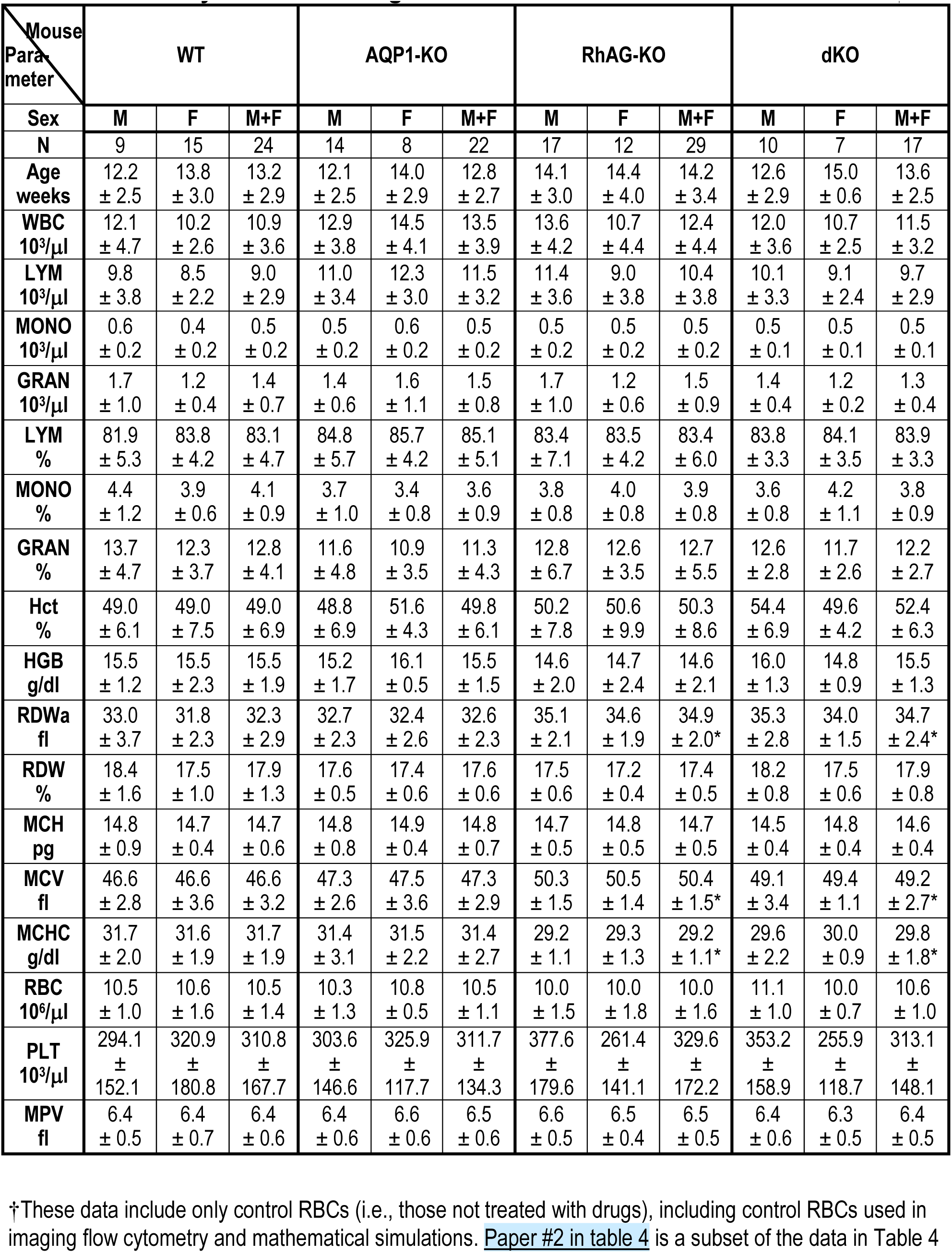
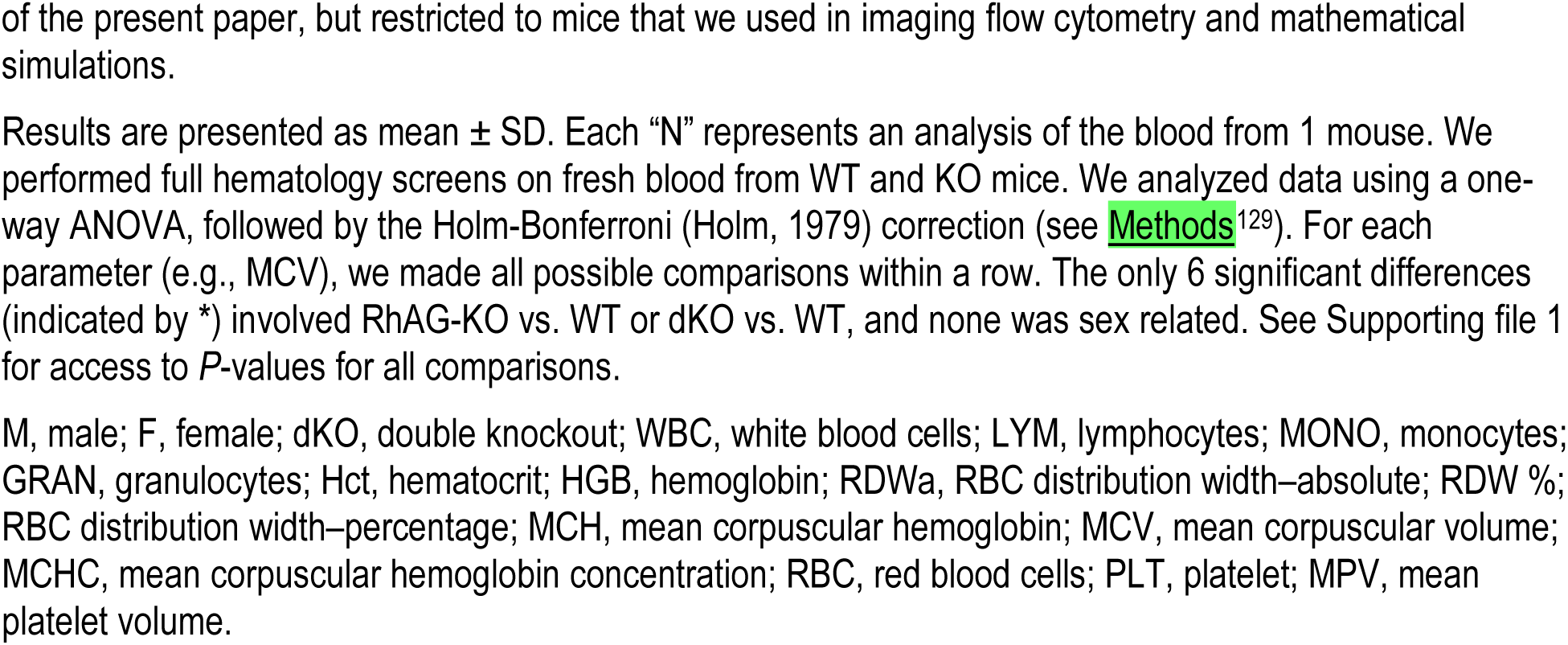
Summary of all hematological data obtained from four strains of mice†.

| Mouse<br>Parameter | WT |  |  | AQP1-KO |  |  | RhAG-KO |  |  | dKO |  |  |
| --- | --- | --- | --- | --- | --- | --- | --- | --- | --- | --- | --- | --- |
| Sex | M | F | M+F | M | F | M+F | M | F | M+F | M | F | M+F |
| N | 9 | 15 | 24 | 14 | 8 | 22 | 17 | 12 | 29 | 10 | 7 | 17 |
| Age<br>weeks | 12.2<br>± 2.5 | 13.8<br>± 3.0 | 13.2<br>± 2.9 | 12.1<br>± 2.5 | 14.0<br>± 2.9 | 12.8<br>± 2.7 | 14.1<br>± 3.0 | 14.4<br>± 4.0 | 14.2<br>± 3.4 | 12.6<br>± 2.9 | 15.0<br>± 0.6 | 13.6<br>± 2.5 |
| WBC<br>10<sup>3</sup>/μl | 12.1<br>± 4.7 | 10.2<br>± 2.6 | 10.9<br>± 3.6 | 12.9<br>± 3.8 | 14.5<br>± 4.1 | 13.5<br>± 3.9 | 13.6<br>± 4.2 | 10.7<br>± 4.4 | 12.4<br>± 4.4 | 12.0<br>± 3.6 | 10.7<br>± 2.5 | 11.5<br>± 3.2 |
| LYM<br>10<sup>3</sup>/μl | 9.8<br>± 3.8 | 8.5<br>± 2.2 | 9.0<br>± 2.9 | 11.0<br>± 3.4 | 12.3<br>± 3.0 | 11.5<br>± 3.2 | 11.4<br>± 3.6 | 9.0<br>± 3.8 | 10.4<br>± 3.8 | 10.1<br>± 3.3 | 9.1<br>± 2.4 | 9.7<br>± 2.9 |
| MONO<br>10<sup>3</sup>/μl | 0.6<br>± 0.2 | 0.4<br>± 0.2 | 0.5<br>± 0.2 | 0.5<br>± 0.2 | 0.6<br>± 0.2 | 0.5<br>± 0.2 | 0.5<br>± 0.2 | 0.5<br>± 0.2 | 0.5<br>± 0.2 | 0.5<br>± 0.1 | 0.5<br>± 0.1 | 0.5<br>± 0.1 |
| GRAN<br>10<sup>3</sup>/μl | 1.7<br>± 1.0 | 1.2<br>± 0.4 | 1.4<br>± 0.7 | 1.4<br>± 0.6 | 1.6<br>± 1.1 | 1.5<br>± 0.8 | 1.7<br>± 1.0 | 1.2<br>± 0.6 | 1.5<br>± 0.9 | 1.4<br>± 0.4 | 1.2<br>± 0.2 | 1.3<br>± 0.4 |
| LYM<br>% | 81.9<br>± 5.3 | 83.8<br>± 4.2 | 83.1<br>± 4.7 | 84.8<br>± 5.7 | 85.7<br>± 4.2 | 85.1<br>± 5.1 | 83.4<br>± 7.1 | 83.5<br>± 4.2 | 83.4<br>± 6.0 | 83.8<br>± 3.3 | 84.1<br>± 3.5 | 83.9<br>± 3.3 |
| MONO<br>% | 4.4<br>± 1.2 | 3.9<br>± 0.6 | 4.1<br>± 0.9 | 3.7<br>± 1.0 | 3.4<br>± 0.8 | 3.6<br>± 0.9 | 3.8<br>± 0.8 | 4.0<br>± 0.8 | 3.9<br>± 0.8 | 3.6<br>± 0.8 | 4.2<br>± 1.1 | 3.8<br>± 0.9 |
| GRAN<br>% | 13.7<br>± 4.7 | 12.3<br>± 3.7 | 12.8<br>± 4.1 | 11.6<br>± 4.8 | 10.9<br>± 3.5 | 11.3<br>± 4.3 | 12.8<br>± 6.7 | 12.6<br>± 3.5 | 12.7<br>± 5.5 | 12.6<br>± 2.8 | 11.7<br>± 2.6 | 12.2<br>± 2.7 |
| Hct<br>% | 49.0<br>± 6.1 | 49.0<br>± 7.5 | 49.0<br>± 6.9 | 48.8<br>± 6.9 | 51.6<br>± 4.3 | 49.8<br>± 6.1 | 50.2<br>± 7.8 | 50.6<br>± 9.9 | 50.3<br>± 8.6 | 54.4<br>± 6.9 | 49.6<br>± 4.2 | 52.4<br>± 6.3 |
| HGB<br>g/dl | 15.5<br>± 1.2 | 15.5<br>± 2.3 | 15.5<br>± 1.9 | 15.2<br>± 1.7 | 16.1<br>± 0.5 | 15.5<br>± 1.5 | 14.6<br>± 2.0 | 14.7<br>± 2.4 | 14.6<br>± 2.1 | 16.0<br>± 1.3 | 14.8<br>± 0.9 | 15.5<br>± 1.3 |
| RDW<sub>a</sub><br>fl | 33.0<br>± 3.7 | 31.8<br>± 2.3 | 32.3<br>± 2.9 | 32.7<br>± 2.3 | 32.4<br>± 2.6 | 32.6<br>± 2.3 | 35.1<br>± 2.1 | 34.6<br>± 1.9 | 34.9<br>± 2.0* | 35.3<br>± 2.8 | 34.0<br>± 1.5 | 34.7<br>± 2.4* |
| RDW<br>% | 18.4<br>± 1.6 | 17.5<br>± 1.0 | 17.9<br>± 1.3 | 17.6<br>± 0.5 | 17.4<br>± 0.6 | 17.6<br>± 0.6 | 17.5<br>± 0.6 | 17.2<br>± 0.4 | 17.4<br>± 0.5 | 18.2<br>± 0.8 | 17.5<br>± 0.6 | 17.9<br>± 0.8 |
| MCH<br>pg | 14.8<br>± 0.9 | 14.7<br>± 0.4 | 14.7<br>± 0.6 | 14.8<br>± 0.8 | 14.9<br>± 0.4 | 14.8<br>± 0.7 | 14.7<br>± 0.5 | 14.8<br>± 0.5 | 14.7<br>± 0.5 | 14.5<br>± 0.4 | 14.8<br>± 0.4 | 14.6<br>± 0.4 |
| MCV<br>fl | 46.6<br>± 2.8 | 46.6<br>± 3.6 | 46.6<br>± 3.2 | 47.3<br>± 2.6 | 47.5<br>± 3.6 | 47.3<br>± 2.9 | 50.3<br>± 1.5 | 50.5<br>± 1.4 | 50.4<br>± 1.5* | 49.1<br>± 3.4 | 49.4<br>± 1.1 | 49.2<br>± 2.7* |
| MCHC<br>g/dl | 31.7<br>± 2.0 | 31.6<br>± 1.9 | 31.7<br>± 1.9 | 31.4<br>± 3.1 | 31.5<br>± 2.2 | 31.4<br>± 2.7 | 29.2<br>± 1.1 | 29.3<br>± 1.3 | 29.2<br>± 1.1* | 29.6<br>± 2.2 | 30.0<br>± 0.9 | 29.8<br>± 1.8* |
| RBC<br>10<sup>6</sup>/μl | 10.5<br>± 1.0 | 10.6<br>± 1.6 | 10.5<br>± 1.4 | 10.3<br>± 1.3 | 10.8<br>± 0.5 | 10.5<br>± 1.1 | 10.0<br>± 1.5 | 10.0<br>± 1.8 | 10.0<br>± 1.6 | 11.1<br>± 1.0 | 10.0<br>± 0.7 | 10.6<br>± 1.0 |
| PLT<br>10<sup>3</sup>/μl | 294.1<br>± 152.1 | 320.9<br>± 180.8 | 310.8<br>± 167.7 | 303.6<br>± 146.6 | 325.9<br>± 117.7 | 311.7<br>± 134.3 | 377.6<br>± 179.6 | 261.4<br>± 141.1 | 329.6<br>± 172.2 | 353.2<br>± 158.9 | 255.9<br>± 118.7 | 313.1<br>± 148.1 |
| MPV<br>fl | 6.4<br>± 0.5 | 6.4<br>± 0.7 | 6.4<br>± 0.6 | 6.4<br>± 0.6 | 6.6<br>± 0.6 | 6.5<br>± 0.6 | 6.6<br>± 0.5 | 6.5<br>± 0.4 | 6.5<br>± 0.5 | 6.4<br>± 0.6 | 6.3<br>± 0.5 | 6.4<br>± 0.5 |

†These data include only control RBCs (i.e., those not treated with drugs), including control RBCs used in imaging flow cytometry and mathematical simulations. Paper #2 in table 4 is a subset of the data in Table 4
of the present paper, but restricted to mice that we used in imaging flow cytometry and mathematical simulations.
Results are presented as mean ± SD. Each “N” represents an analysis of the blood from 1 mouse. We performed full hematology screens on fresh blood from WT and KO mice. We analyzed data using a one-way ANOVA, followed by the Holm-Bonferroni (Holm, 1979) correction (see Methods<sup>129</sup>). For each parameter (e.g., MCV), we made all possible comparisons within a row. The only 6 significant differences (indicated by \*) involved RhAG-KO vs. WT or dKO vs. WT, and none was sex related. See Supporting file 1 for access to *P*-values for all comparisons.
M, male; F, female; dKO, double knockout; WBC, white blood cells; LYM, lymphocytes; MONO, monocytes; GRAN, granulocytes; Hct, hematocrit; HGB, hemoglobin; RDW<sub>a</sub>, RBC distribution width–absolute; RDW %; RBC distribution width–percentage; MCH, mean corpuscular hemoglobin; MCV, mean corpuscular volume; MCHC, mean corpuscular hemoglobin concentration; RBC, red blood cells; PLT, platelet; MPV, mean platelet volume.

### Calculation of raw-*k*HbO_2_ (workflow #3)

Examining the plot of relative HbSat vs. relative time data (see <u>above</u>^12^), we estimate *t*_37_. We then identify the 5 relative HbSat points to the left^13^ and the 5 points to the right of the estimated *t*_37_, and fit the 11 points with a polynomial of degree 2, and solve the best-fit polynomial for the actual relative *t*_37_, and compute the raw-*k*_HbO_2__ as 1/*t*_37_.

The above sequence is step #3 in Figure 1.

### Quantitation of hemolysis in SF reaction cell (workflow #4)

We have developed a novel assay (not depicted here; see Zhao *et al*., 2017), based on the activity of carbonic anhydrase (CA), for determining the degree of RBC hemolysis in the SF reaction cell in a time domain similar to that of our O_2_-efflux experiments (e.g., in Results, see Figure 3*A* and *C*). We use the SF apparatus to mix solutions *A* and *B* (see Table 2) in the SF reaction cell, and create an out-of-equilibrium (OOE) 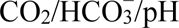 state, chosen such that the pH at the instant of mixing is too low for equilibrium (in Results, see Figure 6*A*). The OOE solution then re-equilibrates according to the reaction 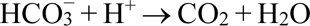 (i.e., raising pH), catalyzed by CA released from hemolyzed RBCs (in Results, see Figure 6*B–D*).

Solution *A* contains HEPES/pH 7.03 + RBCs or hemolysate. Solution *B* contains ∼1% CO_2_/44 mM 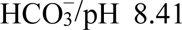 + the fluorescent pH-sensitive dye pyranine (catalog no. H348, Invitrogen), which reports extracellular pH (pH_o_). We excite the dye at 460 nm or 415 nm, while monitoring total fluorescence emission using a 488-nm long-pass filter. Immediately upon mixing of *A* and *B*, the solution in the SF reaction cell (10°C) has the composition ∼0.5% CO_2_/22 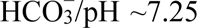, which is out of equilibrium. The pH rises exponentially (rate constant, *k*_ΔpH_, computed as described previously (Zhao *et al*., 2017)) to ∼7.50.

We calculate the actual percent hemolysis (*Act%H*) of ostensibly 100% intact RBCs (i.e., apparent percent hemolysis, *App%H*, is 0%) during a SF experiment, as described previously (Zhao *et al*., 2017), using a procedure that requires three rate constants: *k*_uncat_, *k*_RBC,Lysate_, and *k*_RBC,OstInt_. (1) *k*_uncat_ is the uncatalyzed *k*_ΔpH_ (without CA). (2) *k*_RBC,Lysate_ is *k*_ΔpH_ in the presence of fully lysed RBCs. Finally, (3) *k*_RBC,OstInt_ is *k*_ΔpH_ in the presence of ostensibly 100% intact RBCs. From these, we compute *k*_cat,min_ = *k*_RBC,OstInt_ – *k*_uncat_ and *k*_cat,max_ = *k*_RBC,Lysate_ – *k*_uncat_. Finally,

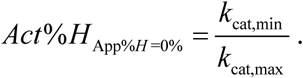

The above equation yields the actual percent hemolysis at a time when the apparent % hemolysis is 0%.

The above sequence is step #4 in Figure 1.

### Accommodation for hemolysis (workflow #5)

**Control and pCMBS-treated RBCs.** Because (1) the RBC membrane has a much higher resistance to O_2_ permeation than does an equivalent thickness of water and (2) NDT can approach HbO_2_ more closely in free solution, Hb released into free solution deoxygenates faster than Hb inside RBCs. We correct raw-*k*_HbO_2__ values (step #3 in Figure 1), based on actual % hemolysis values (step #4 in Figure 1). For each mouse, we generate simulated hemolysis samples for apparent hemolysis levels of 0% (i.e., ostensibly 100% intact RBCs), 2.5%, 5%, 10% and 25% by mixing ostensibly 100% intact RBCs (at relative levels of 100%, 97.5%, 95%, 90%, and 75%) with hemolysate (at relative levels of 0%, 2.5%, 5%, 10%, and 25%)—keeping total [Hb] constant at 5 μM. We determine raw-*k*_HbO_2__ (see <u>above</u>^14^) for each of these simulated hemolysis samples, and—using an algebraic scaling and translation of the x-axis—convert a plot of *k*_HbO_2__ vs. *App%H* (e.g., in Results, see blue axes in Figure 7*A* and *B*) to a plot of *k*_HbO_2__ vs. *Act%H* (see green axes). Extrapolating *k*_HbO_2__ back to 0% actual hemolysis yields an estimate of the actual *k*_HbO_2__ for 1 mouse. We used this approach for all experiments for RBCs treated with pCMBS or no drug.

The above sequence—for RBCs exposed to no drug or to pCMBS—is step #5 in Figure 1.

**DIDS-treated RBCs.** For RBCs treated with DIDS, we modified the <u>above</u>^15^ approach because DIDS decreases the *k*_HbO_2__ of free Hb, so that the actual slope of the *k*_HbO_2__ vs. *Act%H* regression line (*Slope* _DIDS,Act_) is less than the control value (*Slope* _Ctrl,Act_). Two factors are at work. (1) During the 1-h pretreatment of RBCs with DIDS, the drug produces a substantial reduction in *k*_HbO_2__ (in Results, see Figure 8*A*), although this affects only a minute fraction of RBCs that have hemolyzed before entry into the SF device (see Zhao *et al*., 2017). (2) During the actual SF experiment, DIDS produces a small reduction in *k*_HbO_2__ (in Results, see Figure 8*B*) for a larger fraction of Hb newly released in the SF reaction cell over a millisecond/second time frame. For Figure 8*A* and *B*, we obtained free Hb (from hemolysates) and exposed it to DIDS as described in the figure legends.

In a DIDS experiment, the actual slope of the regression line becomes

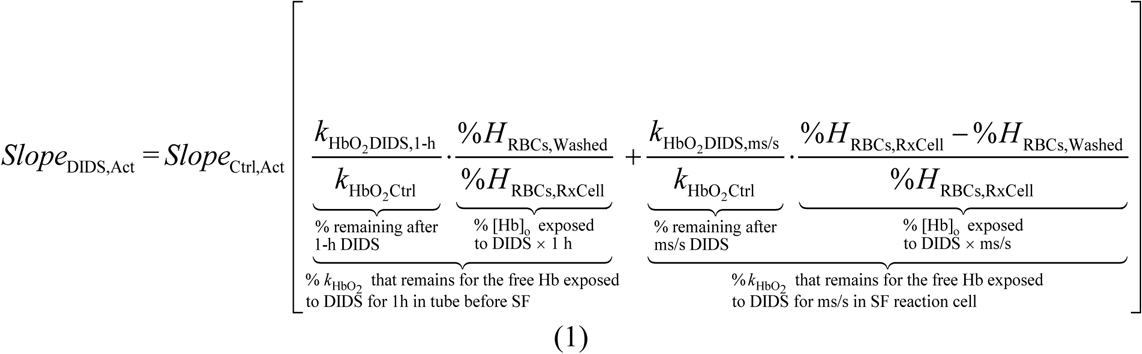

Thus, knowing the *k*_HbO_2__ measured in ostensibly 100% intact RBCs (*k*_HbO_2_ DIDS,OstInt_), we can estimate the actual *k*_HbO_2__ for DIDS-treated RBCs as

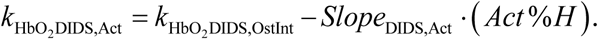

The above sequence—for DIDS-treated cells—is step #5 in Figure 1.

### Measurement of major cell diameter (workflow #6 & #6’)

As described in <u>paper #2</u>^16^, we use imaging flow cytometry to measure the major diameter (Ø_Major_) of cells shaped as biconcave disks or non-biconcave disks. The determination of Ø_Major_ for BCDs is step #6 in Figure 1, whereas the determination of the counterpart Ø_Major_ for nBCDs (assumed in our analysis to be spheres) is step #6′.

### Automated hematological analyses (workflow #7 – #9)

**Assays on normal whole blood.** We collect fresh whole blood into special pipettes (see <u>above</u>^17^), which we insert into a Heska HemaTrue® Veterinary Hematology Analyzer (Heska corporation, Loveland, CO) according to the manufacturer’s protocol. The HemaTrue® analysis yields a wide variety of parameters (introduced below in Table 4), including those related to white and red blood cells and platelets; Hct; MCH (step #7 in the flowchart, Figure 1); MCV (step #8); and MCHC (computed from MCH and MCV in step #9).

**Use of inhibitors in automated hematological studies.** After collecting fresh whole blood from WT or dKO mice, we processed the cells as in “first four assays” (see <u>above</u>^18^), except that—after centrifugation and removal of the buffy coat—we performed only 2 washes (rather than 4) to remove residual extracellular Hb. We then resuspended RBCs in our oxygenated solution (see Table 2), alternatively, (1) without any drug, (2) with pCMBS (1 mM × 15 min), or (3) with DIDS (200 µM × 1 h), to a final Hct of 25%. After centrifuging RBCs at 600 × g for 5 min, we aspirated and discarded the resulting supernatant to remove inhibitors. We then resuspended RBCs in our oxygenated solution to an Hct of 5% to 10%, centrifuged at 600 × g for 5 min, and performed 1 last wash. Finally, we either resuspended packed RBCs at ostensibly 50% Hct for automated hematological assays, or directed the RBCs to studies of <u>flow-cytometry</u>^19^.

The above sequence is step #7–#9 in Figure 1.

### Estimation of thickness of biconcave disks (workflow #10)

As described in <u>paper #3</u>^20^, we generate a torus that has a major outer diameter that matches the measured mean Ø_RBC,major_ of the RBC population (see <u>above</u>^21^), and has a volume that matches the measured mean MCV (see <u>above</u>^22^). We take the RBC thickness as the minor diameter of the torus.

The above sequence is step #10 in Figure 1.

### Determination of rate constant *k*HbO_2_→Hb+O_2_ (workflow #11)

We ascertain the rate constant of the reaction HbO_2_ → Hb + O_2_ in 100% hemolyzed RBCs—or mixtures of hemolysate and ostensibly intact RBCs—as described <u>above</u>^23^ for 100% ostensibly intact RBCs. <u>Above</u>^24^, we also describe the preparation of the lysate.

### Construction of “sigmoid curves” (workflow #12 & #12’)

Introduced below^25^ for BCDs (step #12) and nBCDs (step #12′).

### Estimation of provisional *P*M,O_2_ (workflow #13)

Introduced below^26^.

### Simulations (workflow #14 & #14’)

Introduced below^27^ for BCDs (step #14) and nBCDs (step #14′).

### Determination of prevalence of BCDs and nBCDs (workflow #15 & #15’)

As described in <u>paper #2</u>^28^, we use imaging flow cytometry to determine the fraction of a cell population in our samples represented by various cell types, including BCDs vs. nBCDs.

These estimations are steps #15 and #15′ in Figure 1.

### Macroscopic Mathematical Modeling (MMM) and simulation of HbO_2_-deoxygenation time courses, yielding the “sigmoidal curves” (workflow #12 & #12’)

The details on the macroscopic^29^ mathematical modeling and simulations are in <u>paper #3</u>^30^. In brief, we use a reaction-diffusion model—based on an approach originally developed for CO_2_ fluxes across the membranes of *Xenopus* oocytes by Somersalo *et al*. (2012)—to represent O_2_ efflux from an RBC that we model as a sphere with spherical-radial symmetry. Others have previously modeled RBCs as spheres (Vandegriff & Olson, 1984a; Williams & Kutchai, 1986; Deonikar & Kavdia, 2013; Deonikar *et al*., 2014). These earlier workers took the sphere diameter to be the RBC major diameter, Ø_Major_. In contrast, the value that we assign to the sphere diameter (Ø_Sphere_) is the computed RBC thickness (see <u>above</u>^31^).

As illustrated in Figure 2, the model system comprises (1) a spherical RBC, (2) a plasma membrane that encompasses (3) intracellular fluid (ICF), (4) Hb and HbO_2_ within the ICF, (5) a thin layer of extracellular unconvected fluid (EUF), and (6) an infinite reservoir of bulk extracellular fluid (bECF) that surrounds the EUF, and (7) O_2_ throughout the system. Reactions among O_2_, Hb and HbO_2_ occur only in the ICF. We model the reaction term as the simple, one-step reversible reaction

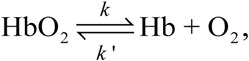

combined with an adaptation (see <u>paper #3</u>^32^) of the “variable rate coefficient” model proposed by Moll (1968), which guarantees that the hemoglobin-O_2_ saturation curve is sigmoidal. All three solutes (i.e., O_2_, Hb and HbO_2_) diffuse within the cytosol. O_2_ is the only solute that can move— via diffusion—through the plasma membrane, within the EUF and between EUF and bECF. Inputs to the simulations include (1) MCV and (2) MCH from automated hematological analyses (see <u>above</u>^33^), (3) the mean major diameter Ø_Major_ of the BCDs (or non-BCDs) from imaging flow cytometry (see <u>paper #2</u>^34^), and (4) the rate constant of HbO_2_ dissociation in an RBC lysate (*k*_HbO_2_→Hb+O_2__), which is equivalent to *k* in the above equation. Paper #3 contains details on the <u>computational model</u> ^35^, <u>parameter values</u> ^36^, and the <u>simulation</u> ^37^ of the time course of deoxygenation of HbO_2_.

**Figure 2.**
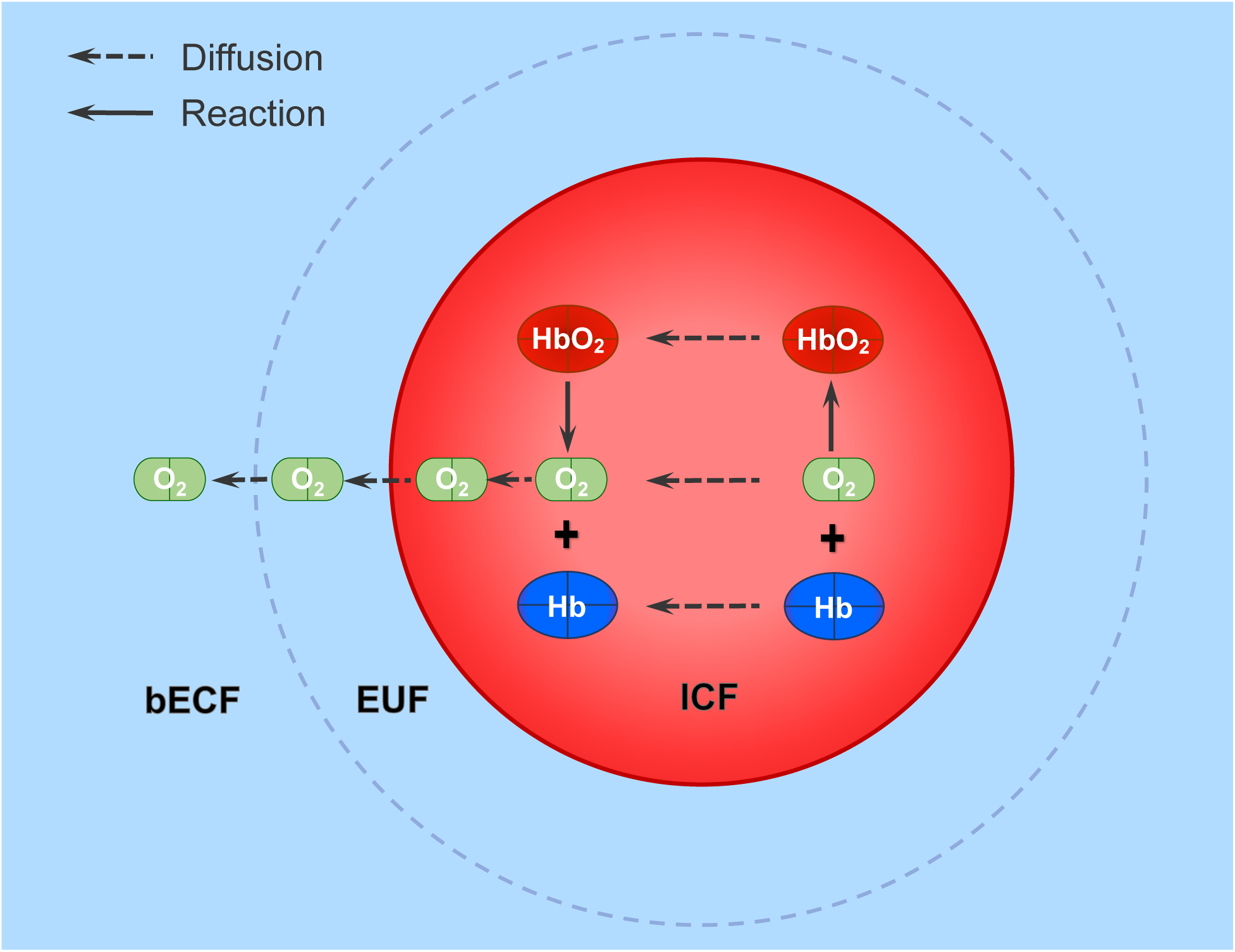
Model of O_2_ offloading from an RBC. The model system comprises (1) a spherical RBC, (2) a plasma membrane (red circle), (3) intracellular fluid (ICF), (4) Hb and HbO_2_ within the ICF, (5) a thin layer of extracellular unconvected fluid (EUF) just external to the PM, (6) an infinite reservoir of bulk extracellular fluid (bECF) that surrounds the EUF and is indicated by grey dashed circle, and (7) O_2_ throughout the system. Arrows: solid black line represents reaction; dashed black line represents diffusion.

Using the MMM together with a set of parameter values—including physical constants, literature values, experimentally determined values (see <u>paper #3</u>^38^), some of which are specific for a particular group of RBCs (e.g., WT controls; see <u>paper #3</u>^39^)—we can insert an assumed *P*_M,O_2__ value into the model and thereby predict the time course of HbSat (result of the reaction HbO_2_ → Hb + O_2_). For each input *P*_M,O_2__ value, this exercise yields a corresponding MMM-*k*_HbO_2__ value. Sweeping through a large number of *P*_M,O_2__ values, we construct the MMM-*k*_HbO_2__ vs. log(*P*_M,O_2__) curves—the “sigmoid curves”—illustrated as steps #12 for BCDs and #12′ for nBCDs in Figure 1.

### Accommodation for nBCDs (workflow #13, #14 & #14’, #16 – #22)

According to (1) our microvideography studies of RBCs on a coverslip, as visualized using a DIC microscopy (see <u>paper #2</u>^40^), and (2) imaging flow cytometry (see <u>paper #2</u>^41^), virtually all RBC samples comprise predominantly biconcave disks, but also some nBCDs. Although a nBCD ought to have the same intracellular total Hb content as a BCD, the intracellular diffusion distances are greater for nBCDs, resulting in a lower rate of O_2_ offloading (see Figure 2). Thus, a mixture of BCDs and nBCDs will have an aggregate *k*_HbO_2__ that is lower than the true *k*_HbO_2__ of BCDs alone.

To arrive at the true *k*_HbO_2__ for BCDs and extract the *P*_M,O_2__, we accommodate for the presence of nBCDs using the novel algorithm comprising steps #13, #14, and #16 through #22 in Figure 1. The principle is straightforward: The time course of HbO_2_ deoxygenation for a mixture of BCDs and nBCDs is the linear combination of the separate deoxygenation time courses of BCDs and nBCDs, each weighted for their relative numbers, as provided by imaging flow cytometry (see workflow #15 & #15′, above and in <u>paper #2</u>^42^). Our goal is to extract the contribution of BCDs from the mixture, as described in <u>paper #3</u>^43^. The following is a brief summary:

**Assignment of Provisional *P*_M,O_2__ (workflow #13).** For our initial estimate of *P*_M,O_2__, we assume that the membrane permeability for O_2_ in mouse RBCs has the same value as the CO_2_ permeability (*P*_M,CO_2__) of human RBCs, 0.15 cm s^−1^, measured by Endeward *et al*. (2006).

**Simulations (workflow #14 & #14′).** Using the *P*_M,O_2__ of 0.15 cm s^−1^ from step #13, we ascertain from the sigmoid curve for BCDs in step #12 that the corresponding MMM-*k*_HbO_2__ value is 3.99 s^−1^ for BCDs (upward and rightward arrows in step #12 of Figure 1). From this MMM-*k*_HbO_2__, we generate a plot of HbSat vs. time for BCDs. In parallel, we generate a second sigmoid curve, but for nBCDs^44^ in step #12′ to ascertain that, for the same *P*_M,O_2__ of 0.15 cm s^−1^, the MMM-*k*_HbO_2__ of nBCDs is only 1.83 s^−1^. From this second *k*_HbO_2__ values, we generate a comparable plot— but with a much slower time course—of HbSat vs. time for nBCDs.

**First linear combination (workflow #16).** A first linear combination of the simulated BCD and nBCD time courses yields an initial estimate of the time course—and *k*_HbO_2__—of the BCD-nBCD mixture.

**Various algebraic manipulations (workflow #17 – #22).** A series of steps involving rebalancing (#17 & #17′); a second linear combination (#18); generation of an inverted sigmoid plot (#19), which we use to compute a shape-corrected (SC) *P*_M,O_2__ of BCDs and nBCDs (#20); calculation of a shape-correction factor (#21) for each of eight individual physiological experimental protocols—WT/control, AQP1-KO/Ctrl, RhAG-KO/Ctrl, dKO/Ctrl, WT/pCMBS, WT/DIDS, dKO/pCMBS, and dKO/DIDS—to obtain an improved (i.e., shape-corrected) estimate of the true *k*_HbO_2__ of BCDs (#22) as well as an estimate for *P*_M,O_2__. We can apply the appropriate shape-correction factor to individual experiment of protocol (e.g., WT/Ctrl).

### Statistical analysis

We report results as mean ± SD. In each relevant figure legend, we report which statistical tests we performed, from among the following, to generate unadjusted *P*-values: (1) paired two-tailed t-test, or (2) one-way analysis of variance (ANOVA). For comparisons of 2 means, we performed #1. For comparisons among >2 means, we performed #2 and then, to control for type I errors across multiple means comparisons, we applied the Holm-Bonferroni correction (Holm, 1979), setting the familywise error rate (FWER) to α = 0.05. Briefly, we order the unadjusted *P*-values for all N comparisons in each dataset from lowest to highest. For the first test, we compare the lowest unadjusted *P*-value to the first adjusted α value, α/N. If the null hypothesis is rejected, then we compare the second-lowest *P*-value to the second adjusted α value, α/(N–1), and so on. If, at any point, the unadjusted *P*-value is ≥ the adjusted α, the null hypothesis is accepted and all subsequent hypotheses in the test group are considered null.

### Data availability

The data supporting the findings of this study are available within the paper. Any further relevant data are available from the corresponding author upon reasonable request.

## Results

In the present study, we use stopped-flow absorbance spectroscopy to monitor Hb deoxygenation as we mix oxygenated RBCs with a solution containing the O_2_ scavenger sodium dithionite (final concentration, 25 mM). This NDT protocol virtually eliminates the effect of the EUF on O_2_ efflux (Coin & Olson, 1979; Vandegriff & Olson, 1984b; Holland *et al*., 1985). We started by investigating the hypothesis that two small-molecule inhibitors (i.e., pCMBS, DIDS) block hypothetical O_2_ channels. After evaluating the effects of genetic deletions of two major membrane proteins (i.e, AQP1, RhAG) on O_2_ offloading from RBCs as part of this paper #1, we tested the hypotheses that observed effects on O_2_ offloading could reflect changes of RBC geometry or changes in the abundance of major RBC membrane proteins as part of paper #2. Finally, the mathematical simulations in paper #3 provide a quantitative basis for interpreting the data in papers #1 and #2, elucidate the dependence of Hb deoxygenation on membrane O_2_ permeability, and support the conclusion that two identified channel proteins as well as a third as-yet-unidentified protein account for nearly all of the O_2_ permeability of murine RBCs.

### Effect of pCMBS or DIDS on O_2_ offloading from RBCs

**Stopped-flow approach.** As detailed in Methods, our general approach is to mix, in the reaction cell of the SF device, (1) RBCs (Hct ≅ 0.37%) suspended in our oxygenated solution (see Table 2) with (2) a deoxygenated solution containing sodium dithionite at a nominal concentration of 50 mM. In the reaction cell, the NDT, now at a nominal concentration of 25 mM, annihilates the extracellular O_2_ in the reaction

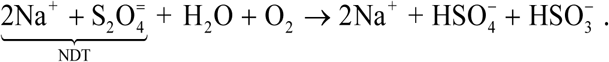

The effect of this level of NDT in eliminating extracellular O_2_ is so powerful that it is presumed to virtually eliminate the effect of the EUF (“unstirred layer”) on O_2_ efflux because [O_2_]_EUF_ ≅ 0 and thus extracellular O_2_ has a negligible effect in slowing the net efflux of O_2_ from the RBC (Coin & Olson, 1979; Vandegriff & Olson, 1984b; Holland *et al*., 1985). The transmembrane O_2_ gradient now strongly favors O_2_ efflux from the RBC (see Figure 2), which in turn leads to the diffusion of free O_2_ from the bulk intracellular fluid to the inner face of the membrane. Throughout the cell, the decrease in [O_2_]_i_ promotes the net reaction HbO_2_ → Hb + O_2_, which we assess by monitoring the absorbance spectrum of Hb. In parallel, HbO_2_ tends to diffuse toward the membrane, and free Hb tends to diffuse in the opposite direction. These processes proceed until no O_2_ remains in the cell and all hemoglobin is in the form of free Hb.

**Effect of pCMBS on O_2_ offloading from RBCs.** The 3D graph in Figure 3*A* shows an example of the time course of Hb absorbance spectra (see <u>Methods</u>^45^) of ostensibly intact, control RBCs harvested from WT mice. We refer to these cells as “ostensibly” intact because, although we take care to load into the SF device only what appear to be intact RBCs (*Act%H* ≅ 0.37%; Zhao *et al*., 2017), we will see that some fraction is lysed in the SF reaction cell; we introduce the accommodation for hemolysis in the next two paragraphs. These are “control” cells because we have not treated them with drugs. In parallel, we pretreat some cells with pCMBS (Figure 3*B*), a sulfhydryl reagent known to inhibit CO_2_ diffusion through AQP1, either with AQP1 expressed *Xenopus* oocytes (Cooper & Boron, 1998), or with native AQP1 in human RBCs (Endeward *et al*., 2006). Figure 3*C* shows a 3D graph for RBCs treated with pCMBS. From the data in Figure 3*A* and *C*, we compute the time courses of relative HbO_2_ saturation (see <u>Methods</u>^46^). The records in Figure 3*D* show that pCMBS markedly slows the deoxygenation of Hb within ostensibly intact RBCs. From each such HbSat-vs.-time record, we obtain a “raw” *k*_HbO_2__ value (see <u>Methods</u>^47^). In the first data row of the blue columns in Table 3*A*, we summarize data for a larger number of such experiments for paired control vs. pCMBS-treated RBCs. Based on raw values, pCMBS reduces *k*_HbO_2__ by 57.7%. The associated *P*-value indicates that the effect of pCMBS is statistically significant.

**Figure 3.**
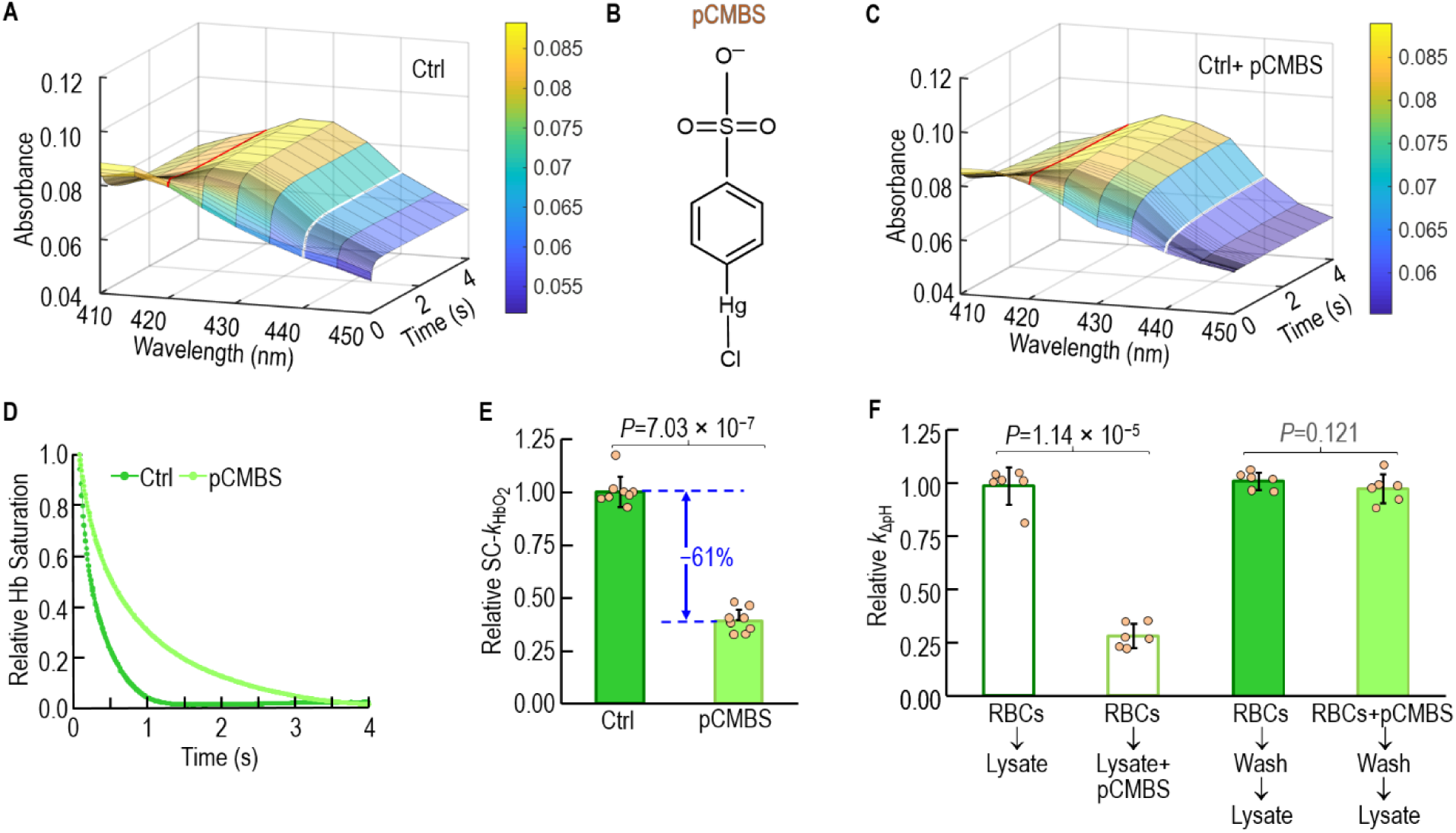
Effect of extracellular pCMBS on rate constant of deoxygenation of intracellular hemoglobin (*k*_HbO_2__) during O_2_ efflux from RBCs. *A,* Representative time course of hemoglobin absorbance spectra during deoxygenation of ostensibly intact wild-type (WT) RBCs under control conditions (Ctrl; absence of drugs). It is from a collection of spectra such as these that we compute raw-*k*_HbO_2__. *B,* Structural formula of p-chloromercuribenzenesulfonate (pCMBS). *C,* Representative collection of absorbance spectra, as in panel *A*, except from WT cells treated with p-chloromercuribenzenesulfonate (pCMBS). *D,* Time courses of relative hemoglobin desaturation (see <u>Methods</u>^132^) for the same two data sets in the previous panels. *E,* Summary of effect of pCMBS on relative *k*_HbO_2__ in intact RBCs, corrected for both hemolysis (see <u>Methods</u> ^133^) and shape changes (see <u>Methods</u> ^134^). The absolute SC-*k*_HbO_2__ underlying the Ctrl bar for this data set is 3.45 ± 0.25 s^−1^. Bars represent mean ± SD, N=8. *F,* Exclusion of pCMBS from RBC cytosol. Wash solution contained 0.2% bovine serum albumin. Bars represent mean ± SD, N=6. For *E & F,* each bar-associated point represents RBCs from 1 WT mouse. Bars represent mean ± SD. We performed paired t-tests (see <u>Methods</u>^135^).

As detailed <u>below</u>^48^, we accommodate each raw-*k*_HbO_2__ value for the small amount of HbO_2_ released into the bulk ECF by RBCs that lyse within the SF reaction cell. Because the released HbO_2_ deoxygenates more rapidly than HbO_2_ surrounded by the RBC membrane, it leads to an overestimate of *k*_HbO_2__. Thus, as summarized for ±pCMBS in the second data row of the blue columns in Table 3*A*, both hemolysis-corrected (HC) *k*_HbO_2__ values are lower than the corresponding raw values. Because the HC-*k*_HbO_2__ falls more for +pCMBS than for –pCMBS, the % inhibition by pCMBS rises somewhat to 63.2%.

Also as summarized <u>below</u> ^49^ —and as detailed in <u>paper #3</u> ^50^ —we additionally accommodate for the appearance of non-biconcave disks, which are rare among control cells but more common with pCMBS-treated and especially DIDS-treated cells. Because the mean diffusion distance is greater (slowing deoxygenation) in nBCDs than in BCDs, the shape-corrected values— summarized in the third data row of the blue columns in Table 3*A*—modestly exceed the corresponding HC-*k*_HbO_2__ values.

After accommodating for hemolysis and nBCDs, we conclude that pCMBS decreases *k*_HbO_2__ of intact BCDs by ∼61% (Figure 3*E*).

Although pCMBS, with its sulfonate group, was designed to be membrane impermeant, a trivial explanation for the drug effects in Figure 3*D* and *E* is that pCMBS somehow gains entry into the RBC and, directly or indirectly, slows the reaction HbO_2_ → Hb + O_2_. To determine whether pCMBS does indeed access the RBC cytosol, we use the RBC’s endogenous, cytosolic carbonic anhydrase as a sentinel. Our CA assay is the same one we use for assessing hemolysis (see <u>Methods</u>^51^), namely, mixing two dissimilar CO_2_/HCO_3_^−^ solutions (see Table 2) in the SF reaction cell to create an out-of-equilibrium state that favors the reaction: 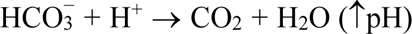, which is governed by the rate constant *k*_ΔpH_. The first bar in Figure 3*F* represents the *k*_ΔpH_ of the untreated RBC lysate. The second bar shows that incubating the hemolysate with pCMBS for 15 minutes markedly reduces lysate CA activity. The last two bars show that treating intact RBCs with pCMBS, and then washing, does not significantly affect the *k*_ΔpH_ of the subsequent lysate (Figure 3*F*). Thus, we conclude that pCMBS—under the conditions of our experiments—does not significantly cross RBC membranes. Therefore, the ability of pCMBS to decrease *k*_HbO_2__ must reflect the action of the chemical from the extracellular side of the membrane.

**Effect of DIDS on O_2_ offloading from RBCs.** Figure 4*A* shows a representative time courses of Hb absorbance spectra of untreated RBCs, panel *B* shows the structure of DIDS, and panel *C* shows a representative time course of Hb absorbance spectra of DIDS-treated RBCs from WT mice. Figure 4*D* shows time courses of relative HbSat in these experiments, and reveals that DIDS markedly slows O_2_ offloading (though not by as much with pCMBS). The first data row of the peach-colored columns of Table 3*A* summarizes paired raw-*k*_HbO_2__ values for a larger number of trials, and shows that DIDS lowers raw-*k*_HbO_2__ by ∼33%. The difference is statistically significant.

**Figure 4.**
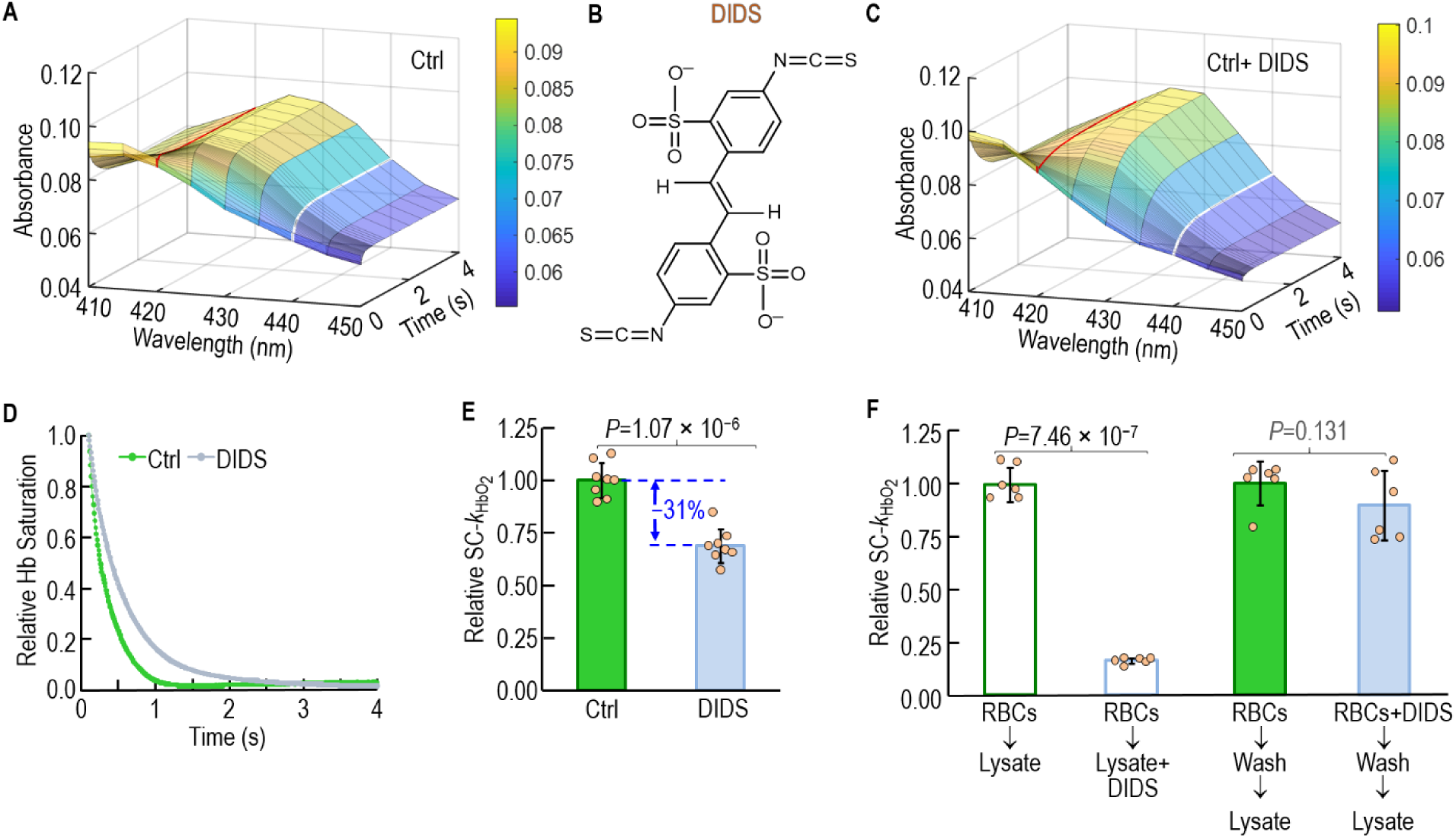
Effect of extracellular DIDS on rate constant of deoxygenation of intracellular hemoglobin (*k*_HbO_2__) during O_2_ efflux from RBCs. *A,* Representative time courses of hemoglobin absorbance spectra during deoxygenation of ostensibly intact control wild-type RBCs. We compute raw-*k*_HbO_2__ from a collection of spectra such as these. *B,* Structural formula of DIDS. *C,* Representative collection of absorbance spectra, as in panel *A*, except from WT cells treated with 4,4’-diisothiocyanatostilbene-2,2’-disulfonate (DIDS). *D,* Time courses of relative hemoglobin saturation (see <u>Methods</u>^132^) for the same two data sets in the previous panels. *E,* Summary of effect of DIDS on relative *k*_HbO_2__ in intact RBCs, corrected for both hemolysis (see <u>Methods</u>^133^) and shape changes (see <u>Methods</u>^134^). The absolute SC-*k*_HbO_2__ underlying the Ctrl bar for this data set is 3.24 ± 0.27 s^−1^. Bars represent mean ± SD, N=8. *F,* Exclusion of DIDS from RBC cytosol. For the data underlying the two open bars, after preparing an osmotic lysate (see Methods^136^), we pretreated the hemoglobin (Hb) ± 200 μM DIDS for 60 min (see Methods^137^), and determined the rate constant (*k*_HbO_2__) for hemoglobin deoxygenation as described in Methods^138^. For the experiments underlying the two filled bars, the wash solution contained 0.2% bovine serum albumin. Bars represent mean ± SD, N=6. For *E & F,* each bar-associated point represents RBCs from 1 WT mouse. Bars represent mean ± SD. We performed paired t-tests (see <u>Methods</u>^135^).

We accommodate these raw-*k*_HbO_2__ values for the Hb released into the bulk ECF by RBCs lysed within the SF reaction cell, using an approach (see <u>below</u>^52^) that is a modification of the one that we employ for control and pCMBS-treated cells. We summarize these HC-*k*_HbO_2__ values in the second data row of peach-colored columns in Table 3*A*. Although hemolysis correction lowers both *k*_HbO_2__ values, it lowers the DIDS HC-*k*_HbO_2__ to a greater extent, so that the DIDS inhibition rises to ∼49%.

Finally, after accommodation for nBCDs, we see in the third data row of peach-colored columns in Table 3*A* that SC-*k*_HbO_2__ rises slightly for the control cells but—because DIDS leads to more nBCDs formation—*k*_HbO_2__ rises more substantially for DIDS-treated cells. After accounting for hemolysis and nBCDs, we conclude that DIDS reduces the SC-*k*_HbO_2__ by ∼31% (Figure 4*E*).

To determine whether DIDS accesses RBC cytosol—and because DIDS (unlike pCMBS in Figure 3*F*) has little effect on CA activity (not shown)—we use cytosolic Hb as a sentinel. The left pair of bars in Figure 4*F* shows that incubating an RBC lysate with DIDS substantially reduces *k*_HbO_2__ of the lysate. However, the right pair of bars reveals no significant effect on *k*_HbO_2__ if we treat ostensibly intact RBCs with DIDS, then wash off the DIDS and, finally, form the lysate in the absence of DIDS. Thus, DIDS does not significantly cross RBC membranes. We conclude (as we did for pCMBS) that the ability of DIDS to decrease *k*_HbO_2__ is a consequence of an action of the chemical on the extracellular side of the membrane.

### Effect of genetic deletions on O_2_ offloading from RBCs

Because our data reveal that neither pCMBS nor DIDS crosses RBC membranes to enter the cytosol (Figure 3*F* and Figure 4*F*), and because we are unaware of evidence that either pCMBS (Vansteveninck *et al*., 1965; Rothstein *et al*., 1973) or DIDS interacts with the lipid component of RBC membranes (Cabantchik & Rothstein, 1974; Moriyama *et al*., 2015), we reason that pCMBS and DIDS must reduce *k*_HbO_2__ by inhibiting membrane proteins that conduct O_2_.

**Effects of single and double knockouts.** We begin our search for such proteins by examining the time course of HbSat of RBCs from mice genetically deficient in AQP1 or RhAG, proteins with established roles as major CO_2_ channels in human RBCs (Endeward *et al*., 2006, 2008). Data from representative experiments show that the speed of O_2_ offloading from Hb gradually decreases in the order WT > AQP1-KO > RhAG-KO > dKO (Figure 5*A*).

**Figure 5.**
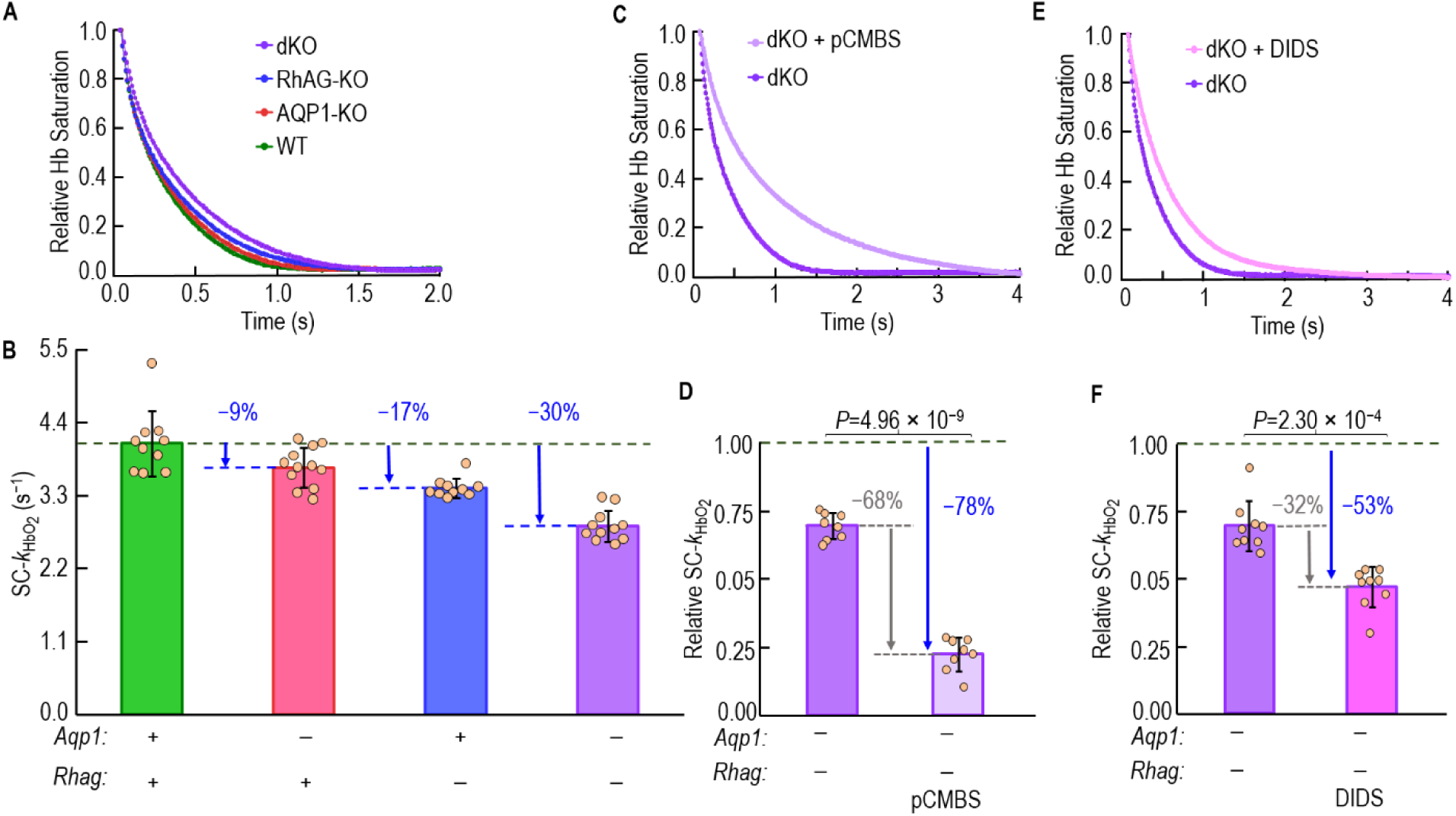
Effect of genetic deletions of *Aqp1*, *Rhag*, or both on rate constant of deoxygenation of intracellular hemoglobin (*k*_HbO_2__) during O_2_ efflux from RBCs. *A,* Representative time courses of hemoglobin desaturation of ostensibly intact RBCs from WT mice and those with genetic deletions of *Aqp1*, *Rhag* or both. It is from these time courses that we compute raw-*k*_HbO_2__. *B,* Summary of mean shape-corrected *k*_HbO_2__ (SC-*k*_HbO_2__) values for the full data set of which panel *A is* representative. The absolute SC-*k*_HbO_2__ underlying the +/+ green bar for the data set in panels *B* is 4.09 ± 0.49 s^−1^. Bars represent mean ± SD, N=10 for green bar, N=12 for red bar, N=10 for blue bar and N=11 for purple bar. We present *P*-values for all the comparisons of SC-*k*_HbO_2__ in Figure 5B (same data sets shown in Table 3*B* of SC-*k*_HbO_2__) in Statistics Table 1. *C & E,* Representative time courses of hemoglobin desaturation of ostensibly intact RBCs, untreated or pretreated with either pCMBS (panel *C*) or DIDS (panel *E*) from dKO mice. It is from these time courses that we compute raw-*k*_HbO_2__. *D & F,* Summary of mean SC-*k*_HbO_2__ for two full data sets of which panels *C & E* are representative. We normalize each of the two purple bars to the height of the purple bar in panel *B*, and appropriately scale the pCMBS and DIDS bars. The three purple bars in panels *B, D, and F* are from three different data sets. The absolute SC-*k*_HbO_2__ for the +/+ control RBCs (not shown here), to which we reference the two purple bars are 2.63 ± 0.18 s^−1^ for panel *D*, and 2.52 ± 0.34 s^−1^ for panel *F*. These reference data are in <u>paper #3></u>table 4. Bars represent mean ± SD, N=8 for panel *D*, N=9 for panel *F*. For panels *B*, *D*, and *F*, each bar-associated point represents RBCs from 1 mouse, with an independent hemolysis correction (see <u>Methods</u>^133^). For *B,* we performed a one-way ANOVA, followed by a Holm-Bonferroni correction (see <u>Methods</u>^135^). For *D & F,* we performed paired t-tests (see <u>Methods</u>^135^).

The four colored columns of Table 3*B* summarize raw-*k*_HbO_2__, HC-*k*_HbO_2__, and SC-*k*_HbO_2__— under control conditions (i.e., absence of drugs)—for a larger number of mice, for each of the four genotypes. The magnitudes of the differences from row to row—raw- to HC- to SC-*k*_HbO_2__—are rather small because both hemolysis and nBCDs prevalence are rather small. The bar-graph summary in Figure 5*B* shows that the genetic deletion of AQP1 reduces SC-*k*_HbO_2__ by ∼9%. The knockout of RhAG produces a decrease of ∼17%, and the double knockout of AQP1 and RhAG lowers SC-*k*_HbO_2__ by ∼30%. Our analysis shows that all six possible comparisons of bars in Figure 5*B* yield statistically significant differences. As summarized below <u>below</u>^53^ and detailed in paper #3, our mathematical simulations show that—because RBC dimensions, Hb content and concentrations, and Hb kinetics vary little among the four genotypes—the percent decreases in *k*_HbO_2__ correspond to far larger decreases in *P*_M,O_2__, ∼55% in the case of comparing RBCs from WT vs. dKO mice.

As described in <u>paper #2</u>^54^, our proteomic analysis of RBC ghosts, which covers >1100 proteins, shows that the two single knockouts and the double knockout appropriately deplete the membranes of AQP1 and/or RhAG and Rhesus blood group D antigen (RhD). However, the two KOs or the dKO do not significantly affect the inferred abundance of any other RBC proteins among the top-ranked 100. Indeed, only 27 proteins (out of the >1100) exhibit a significant change in one of the KOs or the dKO, and these are proteins with relatively low inferred abundance, often originating from cells other than RBCs per se. Of plasma-membrane–associated (PMA) proteins, the one with the highest inferred abundance to exhibit a significant change is the integrin ITGA2A, which <u>ranked</u>^55^ #136 overall and #48 among PMA proteins. Of non-PMA proteins, the highest <u>ranking</u> ^56^ one with a significant change was PSMA4 (#101 overall). These two proteins had baseline inferred abundances <1% of the top-ranked protein, AE1 (i.e., SLC4A1). We conclude that it is highly unlikely that a change in the expression level of an off-target protein could account for our SC-*k*_HbO_2__ data.

**Combined effects of dKO and pCMBS.** We also examined the effect of pCMBS or DIDS on *k*_HbO_2__ in RBCs isolated from dKO mice. Figure 5*C* contains representative time courses of HbO_2_ desaturation for RBCs from a dKO mouse, both under control conditions and after pre-treatment with pCMBS. It shows that O_2_ offloading—already lower in RBCs from dKO vs. WT mice—slows even further after pCMBS treatment.

The blue columns of Table 3*C* summarize paired raw-*k*_HbO_2__, HC-*k*_HbO_2__, and SC-*k*_HbO_2__ for RBCs from dKO mice, under control vs. pCMBS conditions. The differences are all statistically significant. Figure 5*D* shows that, on a dKO background, pCMBS reduces SC-*k*_HbO_2__ by a further ∼68%. Because the dKO by itself produced a ∼30% decrease, this additional decrease caused by pCMBS corresponds to a ∼78% overall decrease in SC-*k*_HbO_2__. Our mathematical simulations <u>below</u>^57^ indicate that this ∼78% decrease in SC-*k*_HbO_2__ corresponds to a ∼91% decrease in *P*_M,O_2__.

**Combined effects of dKO and DIDS.** As illustrated in Figure 5*E* for representative paired experiments on a dKO background, DIDS slows the time course of HbO_2_ desaturation, though not by as much as pCMBS.

The peach-colored columns of Table 3*C* summarize paired raw-*k*_HbO_2__, HC-*k*_HbO_2__, and SC-*k*_HbO_2__ for RBCs from dKO mice, under control vs. DIDS conditions. Figure 5*F* shows that, on a dKO background—which by itself reduces SC-*k*_HbO_2__ by ∼30% decrease—DIDS reduces SC-*k*_HbO_2__ by a further ∼32%. This additional decrease caused by DIDS corresponds to a ∼53% overall decrease in SC-*k*_HbO_2__. Our mathematical simulations <u>below</u>^57^ indicate that this ∼53% decrease in SC-*k*_HbO_2__ corresponds to a ∼76% decrease in *P*_M,O_2__.

### Accommodation for Hemolysis

In a previous study in which we used the same methods as in the present one, the fraction of RBCs hemolyzed was ∼0.37% before they enter the SF device, but ∼5% while the cells were in the SF reaction cell (Zhao *et al*., 2017). This increase presumably reflects the susceptibility of a subfraction of RBCs to the stress of the SF procedure, resulting in cell lysis and the release of Hb and HbO_2_ into the bECF. We hypothesize that this released HbO_2_ (because the RBC membrane does not separate it from the NDT): (1) deoxygenates faster than the HbO_2_ within intact RBCs, leading to (2) an overestimate of *k*_HbO_2__. If RBCs treated with inhibitors or isolated from KO mice have an increased degree of hemolysis, we would underestimate the degree of inhibition. Therefore, for each RBC preparation from each mouse, we assay hemolysis as outlined in <u>Methods</u>^58^.

#### Hemolysis measurements

**Effect of drug treatments on hemolysis.** Figure 6*A* schematically illustrates our approach for assessing hemolysis within the SF reaction cell (Zhao *et al*., 2017). Figure 6*B* schematizes this approach to control cells (i.e., no drugs); Figure 6*C*, to pCMBS-treated cells (i.e., pCMBS); and Figure 6*D*, to DIDS-treated cells (i.e., DIDS). Our analysis of hemolysis shows that, compared to WT/Ctrl RBCs (∼5.2% hemolysis), those pretreated with pCMBS (∼8.2%) or DIDS (∼13.4%) exhibited a significant increase (Figure 6*E*). Note that we accommodate for hemolysis, as described <u>below</u>^59^.

**Figure 6.**
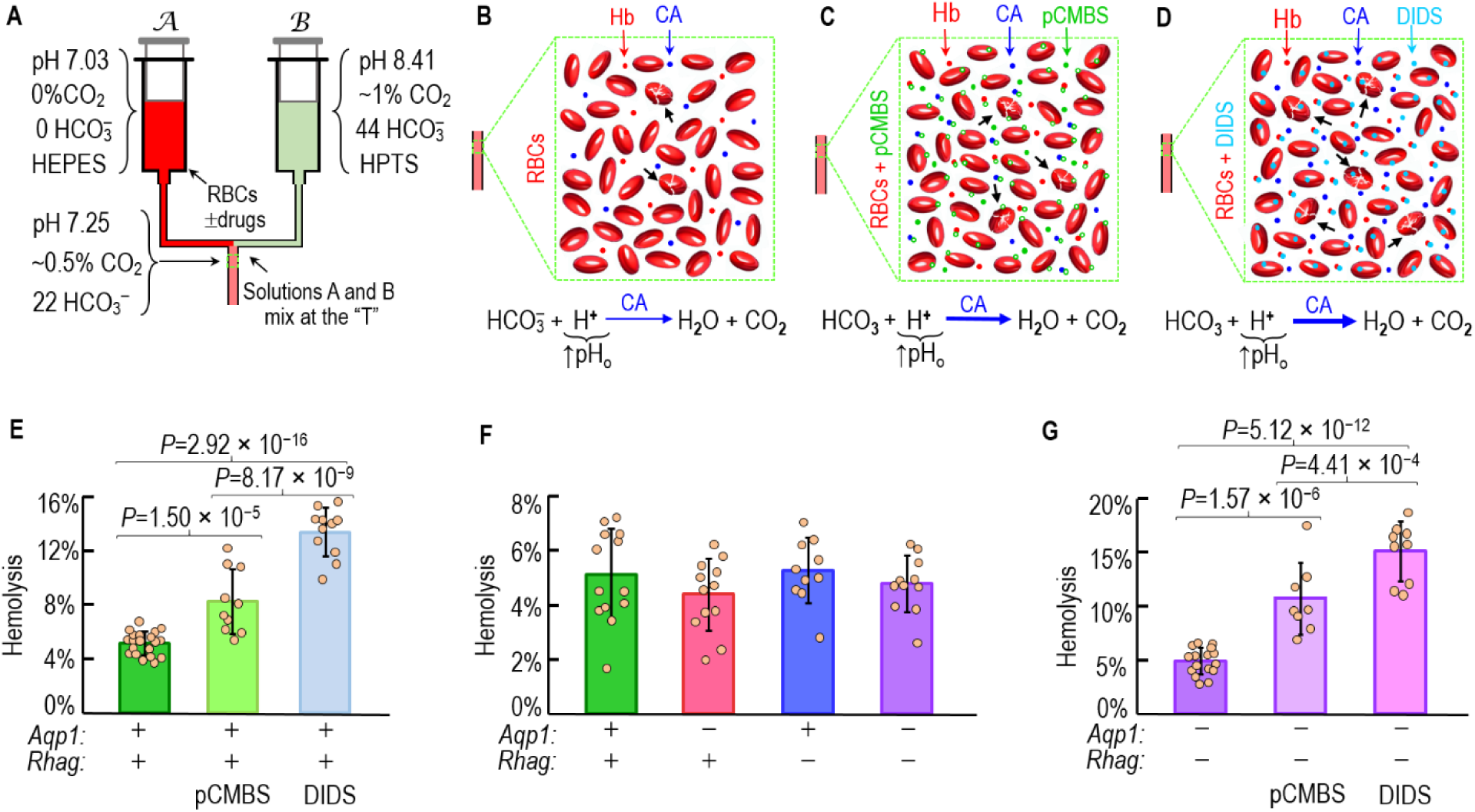
Degree of hemolysis in experiments on ostensibly intact RBCs. *A,* Schematic diagram of CO_2_/HCO^−^ out-of-equilibrium (OOE) studies in stopped-flow (SF) apparatus. Solutions *A* and *B* mix at the “T” and thereby create, in the SF cell, a new solution that is out of equilibrium with ∼0% CO_2_/22 mM HCO^−^/pH 7.25 at 10°C. *B,* Control cells (without drugs). Upper panel, schematic diagram of Ctrl RBCs in an SF cell. Hemolysis is typically 0.4% as we load RBCs into the SF machine. However, in the SF reaction cell, actual hemolysis rises to ∼5%. The lysed cells (black Arrows) release both hemoglobin (Hb, red dots) and carbonic anhydrases (CAs, blue dots). Lower panel, CO_2_/HCO^−^ reaction. Starting from the OOE state, HCO^−^ and H^+^ combine to form CO_2_ and H_2_O. Overall reaction is greatly accelerated by the CAs released along with Hb from the lysed RBCs. Our hemolysis assay (Zhao *et al*., 2017) is based on the rate constant of the pH increase (*k*_ΔpH_) caused by this reaction. *C,* pCMBS-treated cells. Upper panel, schematic diagram of pCMBS-treated RBCs in an SF cell. Actual hemolysis in the SF reaction cell is ∼8% for pCMBS-treated RBCs from WT mice. The small amount of lysis that occurs during the 15-min pCMBS preincubation period, before the cells enter the SF reaction cell (∼0.4%), releases CA that reacts extensively with pCMBS, markedly reducing that enzymatic activity (see Figure 3F, open pair of bars). Later, free pCMBS enters the SF reaction cell and is exposed to a much higher levels of CA from RBCs that lyse in the SF reaction cell (∼7.5%), but only for a very brief period (100’s of ms). Thus, pCMBS reacts with only a tiny fraction of the newly released CA, so that extracellular CA activity is high in the SF reaction cell. The filled green color dots indicate free pCMBS, whereas the open green dots represent Hg that has reacted with the cell surface, released CA (blue dots), or released Hb (red dots). Lower panel, CO_2_/HCO^−^ reaction. Elevated extracellular CA activity from the increased hemolysis speeds the reaction and increases *k*_ΔpH_, and reports the elevated hemolysis. *D,* DIDS-treated cells. Upper panel, schematic diagram of DIDS-treated RBCs in an SF cell. Actual hemolysis in the SF reaction cell is ∼13.4% for DIDS-treated RBCs from WT mice. Unlike pCMBS (panel *C*), DIDS has little effect on CA activity, so that the chemistry of the hemolysis assay is basically the same as in panel *B*. (Note that DIDS does react with Hb as shown in Figure 4F but HbO_2_ deoxygenation is not part of our hemolysis assay). The filled teal-colored dots indicate DIDS, some of which remains free, and some of which has reacted with the cell surface and released Hb, but not released CA. Lower panel, CO_2_/HCO^−^ reaction. Because hemolysis is greater than with pCMBS in panel *C*, *k*_ΔpH_ is greater. *E,* Summary of hemolysis in experiments on WT-mouse RBCs, either controls or cells treated with pCMBS or DIDS. RBCs were treated with 1 mM pCMBS for 15 min or 200 μM DIDS for 60 min. N=20 for green bar, N=10 for light green bar and N=11 for gray-blue bar. *F,* Summary of hemolysis in experiments on control mouse RBCs of all four genotypes used in the present study. Note the compressed scale of the y-axis, compared to panel *A*, which has bars for pCMBS- or DIDS-treated cells. The RBCs from WT-mice here in panel *F* represent a different data set than in panel *E*. N=13 for green bar, N=12 for red bar, N=10 for blue bar and N=11 for purple bar. We present *P*-values for all statistical comparisons in Statistics Table 2. *G,* Summary of hemolysis in experiments on RBCs from double-knockout mice, either controls or cells treated with pCMBS or DIDS. N=17 for purple bar, N=8 for light purple bar and N=9 for pink bar. In panels *E*, *F*, and *G*, each bar-associated point represents 1 mouse. Bars represent mean ± SD. We performed a one-way ANOVA, followed by the Holm-Bonferroni correction (see <u>Methods</u>^135^).

**Effect of channel knockouts on hemolysis.** Our analysis of the two KO sub-strains and the dKO reveal no significant difference in hemolysis compared to the WT (Figure 6*F*). These results are consistent with the report that deletion of *Rhd* and *Rhag* do not increase RBC fragility (Goossens *et al*., 2010).

Among RBCs from dKO mice, we find that pretreatment with pCMBS or DIDS—as was the case for RBCs isolated from WT mice—produce significant increases in hemolysis (Figure 6*G*). Moreover, pCMBS and DIDS produce somewhat greater degrees of hemolysis on the dKO background (Figure 6*G*) than on the WT background (Figure 6*E*). Note that we accommodate for hemolysis, as described immediately below.

#### Accommodation of raw-*k*_HbO_2__ for hemolysis

We have developed two variants of a novel approach to accommodate for hemolysis in the SF reaction cell. The first variant applies to cells that are either not treated with a drug (i.e., controls) or those pretreated with an agent(s) that does not appreciably affect the rate of the extracellular reaction HbO_2_ → Hb + O_2_. The second variant applies to experiments in which the agent(s) (e.g., DIDS) does significantly affect this reaction.

**Control or pCMBS-treated RBCs.** Figure 7*A* shows the result of applying this approach (as illustrated in Figure 6*A* and *B*) to a representative control experiment (i.e., no drugs) on RBCs from a WT mouse. In this figure, the blue x-axis is the apparent hemolysis, whereas the green x-axis is the actual hemolysis, measured as described in <u>Methods</u>^60^. One might unwittingly assume that the *Act%H* in SF reaction cell is 0% (i.e., 100% of RBCs are ostensibly intact), and thus believe that the measured raw-*k*_HbO_2__ of 3.70 s^−1^ (the value in step #3 of Figure 1) in this experiment is in fact the actual *k*_HbO_2__ (although not yet shape corrected). However, the hemolysis assay in this example shows that *Act%H* is in fact 4.4% (shown on the green x-axis). To determine the actual *k*_HbO_2__ when *Act%H* truly is 0%, we mix 100% ostensibly intact RBCs (*Act%H* ≅ 4.4%) with 100% hemolyzed RBCs (*Act%H* = 100%) in increasing ratios, as shown by the blue points in Figure 7*A*. When we reference these blue points to the blue x-axis, we obtain a regression line that describes apparent raw-*k*_HbO_2__ vs. *App%H*, with a slope of ∼5.5256 s^−1^/%*H* and a y-intercept (i.e., the raw-*k*_HbO_2__) of ∼3.70 s^−1^ on the blue axes. To reference these same blue points to the green x-axis, we scale the x-axis and the slope of the regression line by the factor

**Figure 7.**
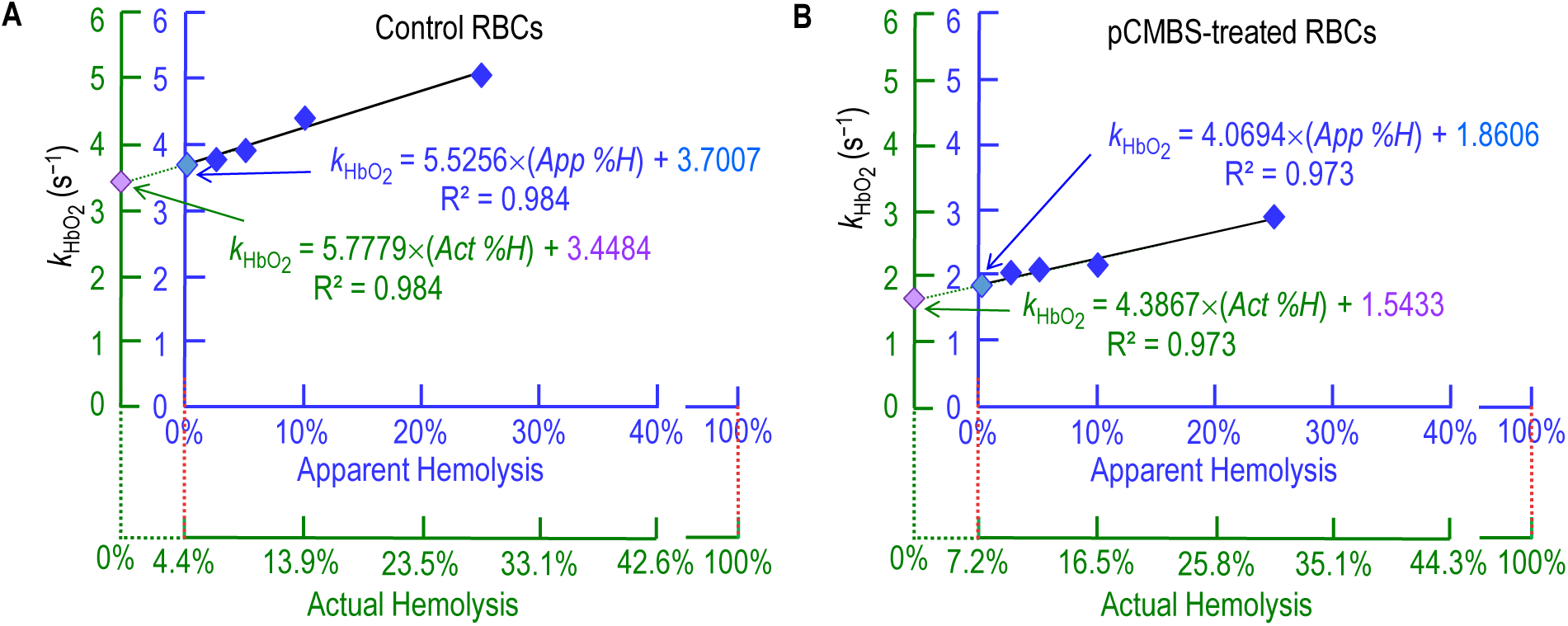
Accommodation of raw-*k*_HbO_2__ for hemolysis in experiments on ostensibly intact RBCs, either without inhibitors (*A*), or pretreated with pCMBS (*B*) on blood from the same WT mouse. The raw rate constant (*k*_HbO_2__) is a mixture of contributions from (1) the relatively slow deoxygenation of cytosolic HbO_2_ in truly intact RBCs and (2) the relatively rapid deoxygenation of extracellular HbO_2_ released from hemolyzed RBCs. Here, we use a novel approach to correct raw-*k*_HbO_2__ for hemolysis in a series of five steps: (1) Use the assays in Figure 6 to determine actual percent hemolysis (*Act%H*) of ostensibly 100%-intact RBCs as described in Methods^139^ for control RBCs (4.4% on green x-axis for mouse in panel *A*) and pCMBS-treated RBCs (7.2% on green x-axis for mouse in panel *B*). (2) Determine *k*_HbO_2__ for ostensibly 100%-intact RBCs—that is, apparent percent hemolysis [*App%H*] = 0%— as described in Methods^140^. These are the light-blue diamonds with the black perimeters in panels *A* and *B* (on the blue y-axes). (3) Determine *k*_HbO_2__ for mixtures of ostensibly intact RBCs (97.5%, 95%, 90%, and 75%) and fully hemolyzed RBCs (2.5%, 5%, 10%, and 25%) to keep total [Hb] = 2.5 μM in the SF reaction cell. The *App%H* values are on the blue x-axes. (4) Rescale and translate the blue *App%H* axes to obtain the green *Act%H* axes for mixtures of ostensibly 100%-intact RBCs and 100% hemolysate: (*Act%H*) = (*App%H*) + (*Act%H*_App%H=0%_)*(100% – *App%H*). (5) Back-extrapolate *k*_HbO_2__ to *Act%H* = 0% to obtain hemolysis-corrected *k*_HbO_2__ values (lavender diamonds), which for the sample in panel *A* was 3.45 s^−1^ for control RBCs and, for the sample in panel *B*, 1.54 s^−1^ for pCMBS-treated RBCs. We repeat this analysis for each mouse.

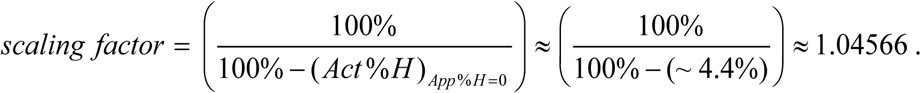

We also translate the green x-axis so that the measured *Act%H* (i.e., ∼4.4%) lines up with the *App%H* of 0% on the blue x-axis. We calculate actual *k*_HbO_2__ by back-extrapolating the regression line—now referenced to the scaled/translated green axes—to *Act%H* = 0. As illustrated in Figure 7*A*, here the y-intercept is ∼3.45 s^−1^, which represents the hemolysis-corrected *k*_HbO_2__ (the value in step #5 of Figure 1). As we will see at the end of Results, the further correction of these RBCs for shape yields a SC-*k*_HbO_2__ of 3.48 s^−1^ (the value in step #22 of Figure 1).

We apply the above approach to the pCMBS-treated cells as schematized in Figure 6*C*, and exemplified in Figure 7*B*. Indeed, we performed the hemolysis accommodation, as in Figure 7, for all experiments, except those for cells pretreated with DIDS—as discussed next.

**DIDS-treated RBCs.** As noted in <u>Methods</u>^61^, for two reasons we must approach the accommodation for hemolysis somewhat differently in DIDS-pretreated RBCs than in control or pCMBS-treated cells. First, in 100%-hemolyzed RBCs, a 1-h pretreatment with DIDS markedly (∼85%) reduces *k*_HbO_2__ (compare green and gray-blue bars in Figure 8*A*. The large difference in bar heights is relevant for RBCs as we load them into the SF device, when the fraction of hemolyzed cells is minute (i.e., *Act%H* ≅ 0.37%; see Zhao *et al*., 2017), but the duration of drug exposure is sizeable. Second, during the actual SF experiment—and on a ms-to-s time scale—DIDS produces only a small (∼4%) reduction in *k*_HbO_2__ (compare green and gray-blue open bars in Figure 8*B*). In this case, although the fractional inhibition is small, the fraction of Hb newly released in the SF reaction cell is much larger.

**Figure 8.**
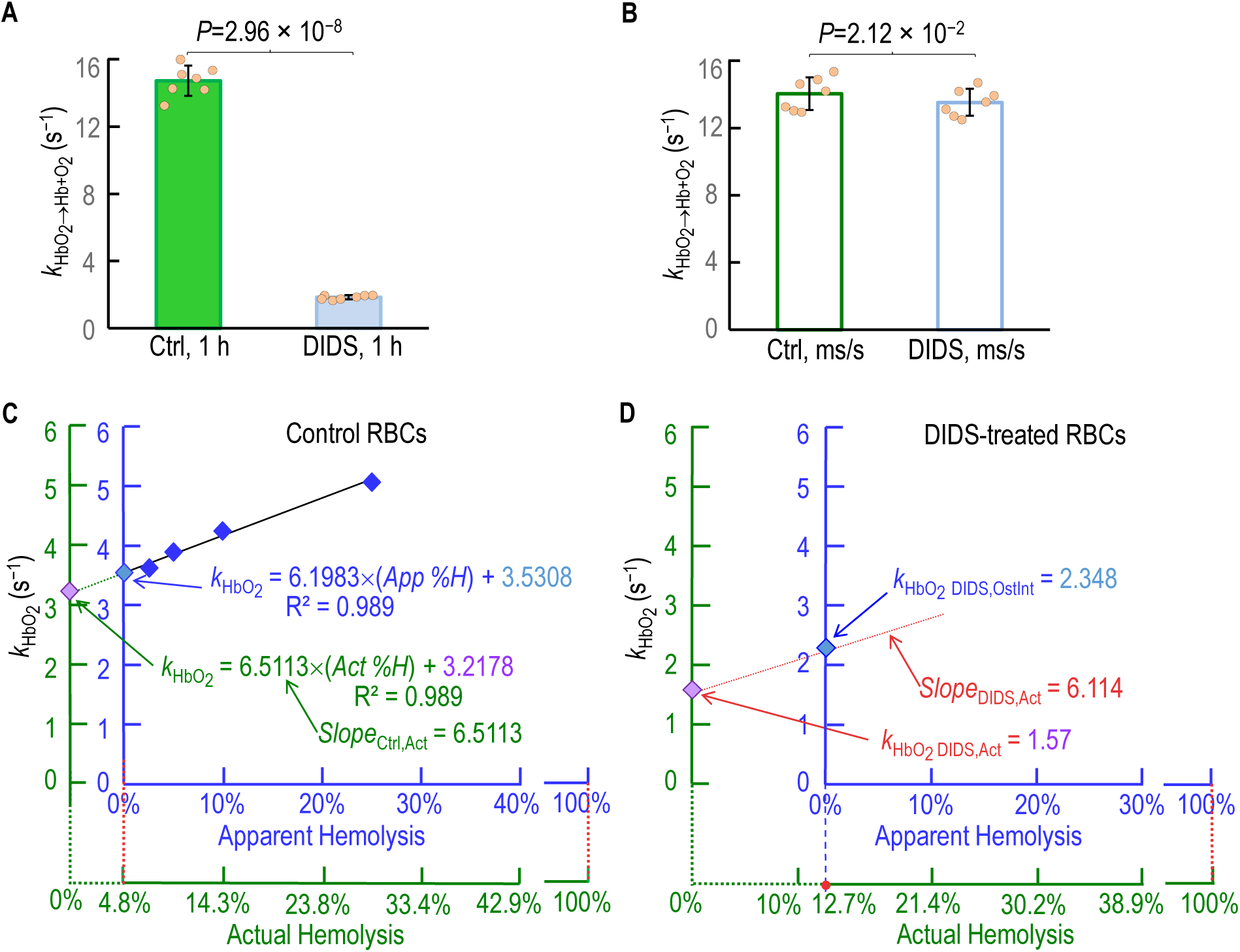
Correction of *k*_HbO_2__ for hemolysis in experiments on RBCs treated with DIDS. *A,* Effect of DIDS on *k*_HbO_2__ in a 100% hemolysate over a 1-hour time frame. This experiment is basically the same as the one summarized by the two open bars in Figure 4F, but on a different data set. After preparing an osmotic lysate (see Methods^136^), we pretreated the hemoglobin (Hb) ±200 μM DIDS for 60 min (see Methods^137^, and determined the rate constant (*k*_HbO_2__) for hemoglobin deoxygenation as described in Methods^138^. *B,* Effect of DIDS on *k*_HbO_2__ in a 100% hemolysate over millisecond-to-second timeframe. After (1) preparing an osmotic lysate as in panel *A*, we (2) diluted the hemolysate in our oxygenated solution and the DIDS (200 μM final concentration) in our deoxygenated solution (see Table 2) as described in Methods^141^, and (3) after rapid mixing of hemolysate and DIDS in the SF reaction cell, (4) determined *k*_HbO_2__ as in panel *A*. For *A* & *B,* each bar-associated point represents RBCs from 1 WT mouse. Bars represent mean ± SD. Ctrl, control (i.e., no drugs). N=7 for each group. We performed paired t-tests. *C,* Determination of control slope (*Slope*_Ctrl,Act_) in plot of *k*_HbO_2__ vs. actual percent hemolysis (*Act%H*, green) for RBCs without inhibitors. This is the same protocol as in Figure 7A, though on a different data set. The Extrapolation of the *k*_HbO_2__ data back to *Act%H* = 0% yields an actual *k*_HbO_2__ (lavender diamond on green y-axis), which in this example was 3.22 s^−1^ for control RBCs. *D,* Correction of *k*_HbO_2__ for hemolysis in experiments on RBCs pretreated with DIDS. We determined raw-*k*_HbO_2__ values for ostensibly “100%-intact” RBCs as described for panel *C*. For the sample here in panel *D* (from same mouse as in panel *C*), this uncorrected value (blue diamond on blue y-axis) was ∼2.35 s^−1^ for DIDS-treated RBCs. We calculate the slope (*Slope*_DIDS,Act_) of the regression line (red dashed line) using the data in panel *C*, as described in Methods^142^. Extrapolation of this line yields a hemolysis-corrected *k*_HbO_2__ (indicated as lavender diamond (on green y-axis) for *k*_HbO_2_ DIDS,Act_) of ∼1.57 s^−1^.

In the control RBCs in Figure 8*C*, the approach for obtaining the regression line is the same as in Figure 7*A*. Figure 8*C* shows the *k*_HbO_2__ hemolysis correction for RBCs—from 1 mouse— studied under control conditions (i.e., no DIDS). Here, *Slope* _Ctrl,Act_ is 6.51 s^−1^/*Act%H*.

For the DIDS-treated cells in Figure 8*D*, we use a novel approach to obtain the regression line. In Methods, Eqn (1) describes (without a numerical example) the actual slope of this regression line. For RBCs from the same mouse as in Figure 8*C*, but treated ×1 h with 200 μM DIDS before entry into the SF device, the *Act%H* is 12.7% in the SF reaction cell. Inserting these %*H* values for this mouse—and the average *k*_HbO_2__ for 7 mice in Figure 8*A–B*—into Eqn (1) yields

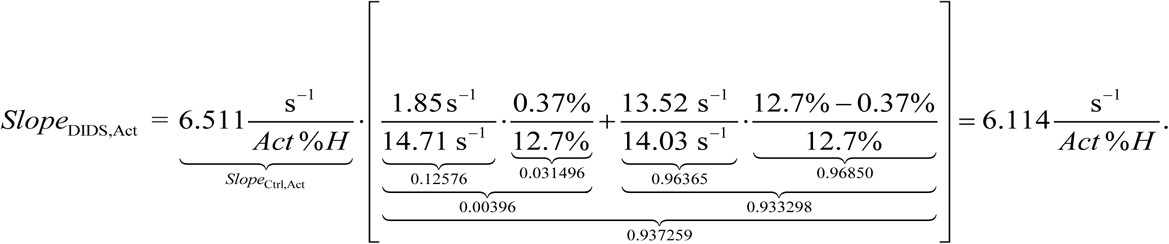

Note that although this slope correction^62^ amounts to only 100% – (∼93.7%) = ∼6%, it affords improved accuracy and shows that potential errors are not likely to be large. As summarized in Figure 8*D*, for this particular mouse^63^

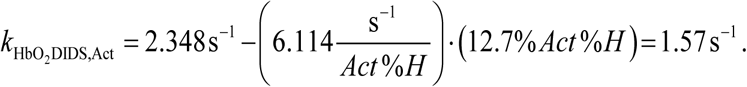

### Hematological and related parameters

We present the details of the macroscopic mathematical model as well as simulations of O_2_ offloading in paper #3. In the present paper, <u>below</u>^64^, we summarize elements of this MMM that are of particular relevance to the present paper. It is perhaps intuitively obvious that *k*_HbO_2__ should decrease with (1) increases in MCV (due to increased diffusion distances) and (2) increases in MCHC (due to increased interactions between O_2_ and Hb that slow effective O_2_ diffusion), and (3) with decreased kinetics of HbO_2_ dissociation. To evaluate potentially trivial explanations for the decreases in *k*_HbO_2__ observed in Figure 3, Figure 4, and Figure 5, we performed a series of hematological, morphometric, and HbO_2_ kinetics studies.

#### Automated hematology

**Effect of KOs.** We performed a battery of assays (see <u>Methods</u>^65^) on the fresh blood of a large subset of mice used in the present study. In the largest group of such studies, we examined all four genotypes (Table 4). Because we observe no significant sex-related differences, we grouped the data from male and female mice (M+F). The only genotype-related differences that reach statistical significance are with RBCs from RhAG-KO and dKO mice, which for M+F exhibit slightly higher mean values for RDWa and MCV, but slightly lower values for MCHC.

The slightly higher RDWa values may correlate with the trend for KO mice, particularly RhAG-KO and dKO, to have higher percentages of reticulocytes (see paper #2^66^).

Regarding the increased MCV but decreased MCHC values, our mathematical simulations indicate that these effects tend to cancel (<u>paper #3</u>^67^). The bar graphs in Figure 9 summarize the most pertinent automated hematology data, including those with statistically significant differences (see Figure 9*C*, *E*, *F*).

**Figure 9.**
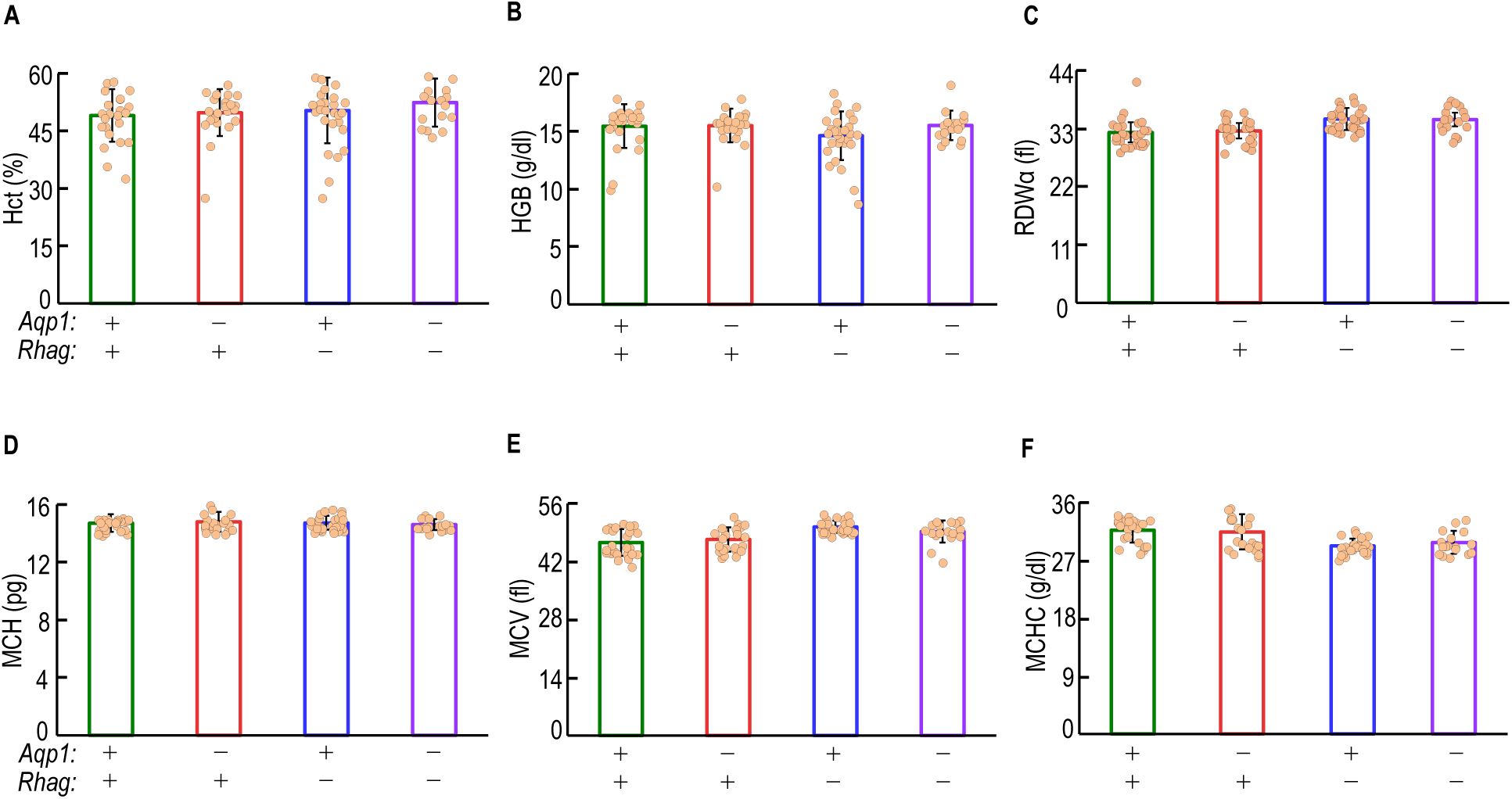
Effect of genetic deletions of *Aqp1*, *Rhag*, or both on key hematological parameters. The bar graphs summarize key parameter values—with data from both sexes combined—from automated hematological analyses. See Table 4 for the detailed data, separated by sexes, on WT, *Aqp1*–/–, *Rhag*–/–, and dKO mice). *A,* Hematocrit (Hct). *B,* Hemoglobin (HGB). *C,* RBC distribution width—absolute (RDWa). *D,* Mean corpuscular hemoglobin content (MCH). *E,* Mean corpuscular volume (MCV). *F,* Mean corpuscular hemoglobin concentration (MCHC). Each bar-associated point represents RBCs from 1 mouse. Bars represent mean ± SD. N=24 for green open bar, N=22 for red open bar, N=29 for blue open bar and N=17 for purple open bar. We performed a one-way ANOVA, followed by the Holm-Bonferroni correction (see Methods^135^). We present *P*-values for all statistical comparisons in for panels *A*-*F* in Statistics Table 3.

**Effects of inhibitors.** For hematological studies with inhibitors (see <u>Methods</u>^68^), we generated preparations of RBCs from WT and dKO mice, and then pretreated these with no drug, pCMBS, or DIDS—a total of six conditions ([WT vs. dKO] × [control vs. pCMBS vs. DIDS]). Most of the statistically significant effects (Table 5) involve white blood cells or platelets, which the drugs generally tend to reduce in number. The effects on RBCs mainly involve MCV and MCHC, and these effects—as noted in the previous paragraph—are almost always in the directions that would tend to cancel vis-à-vis *k*_HbO_2__.

**Table 5.**
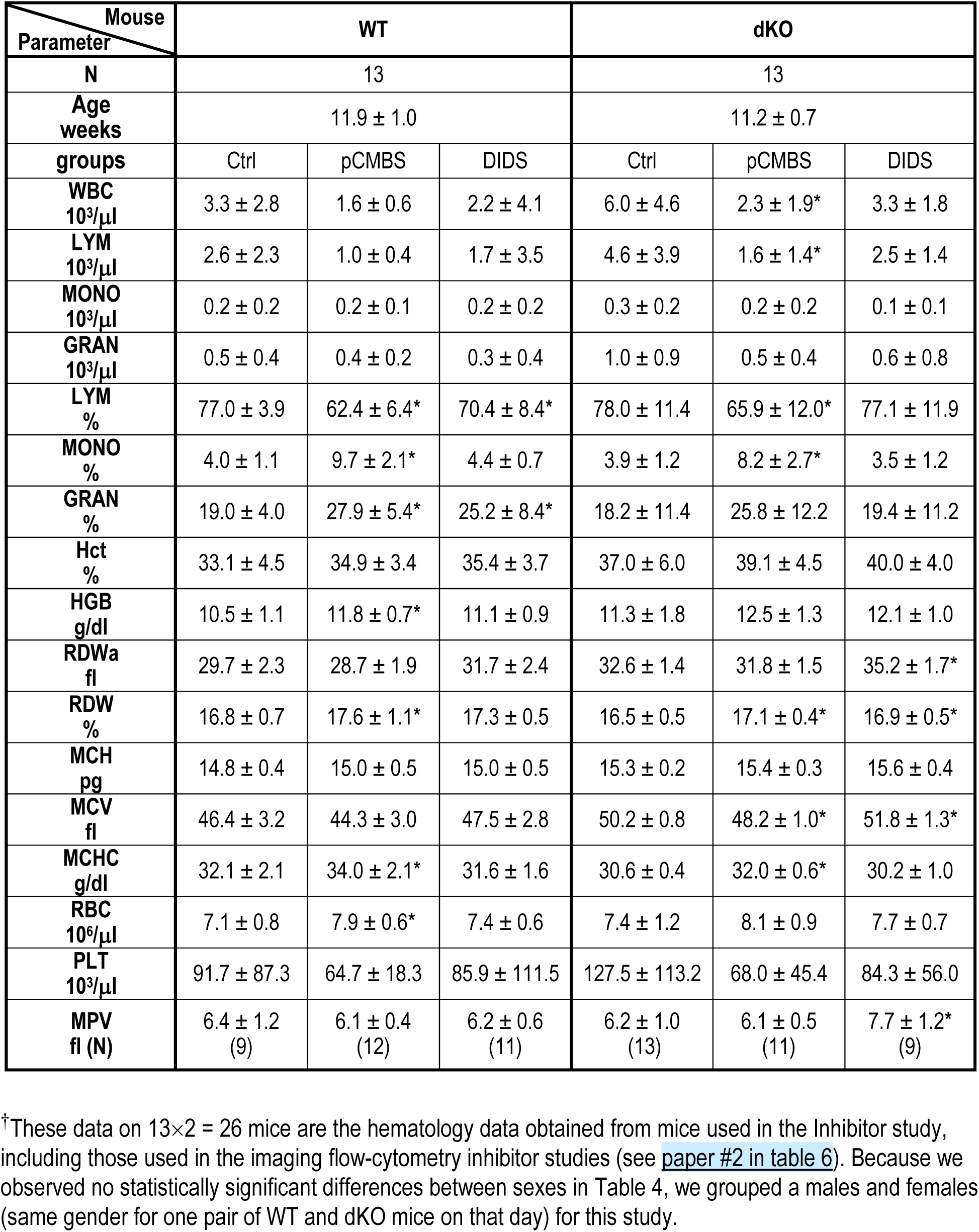
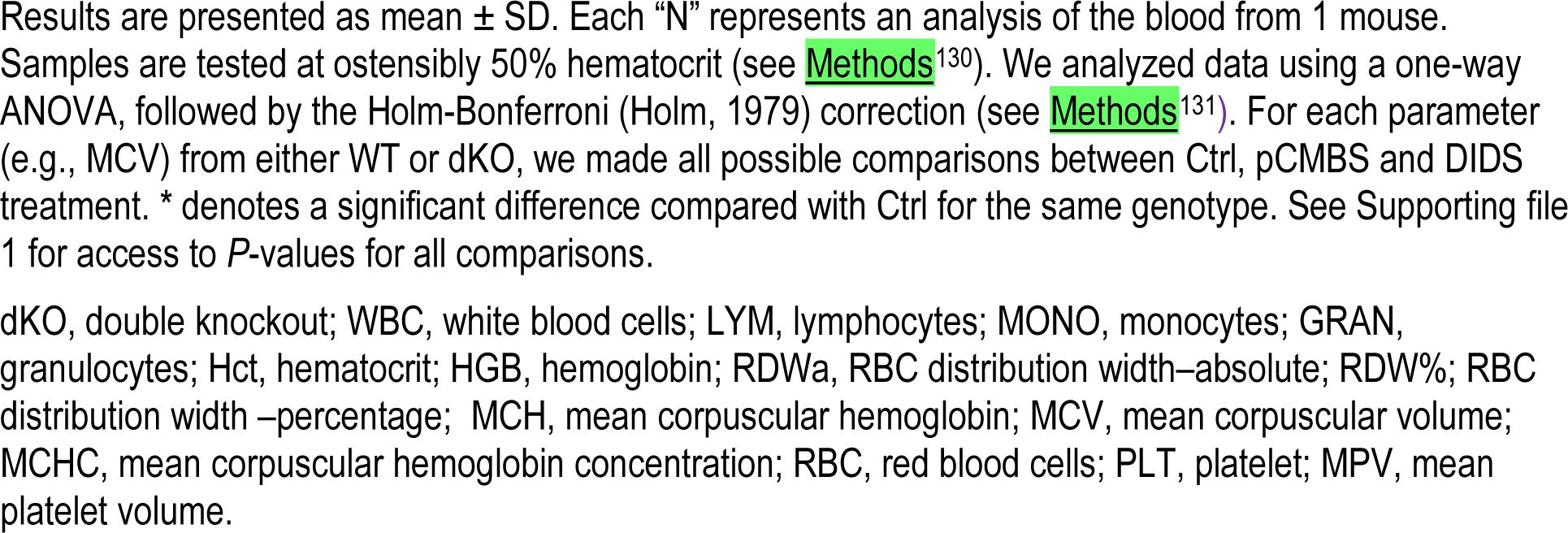
Summary of automated hematology data obtained from mice used in Inhibitor study†.

| Mouse<br>Parameter | WT |  |  | dKO |  |  |
| --- | --- | --- | --- | --- | --- | --- |
| N | 13 |  |  | 13 |  |  |
| Age weeks | 11.9 ± 1.0 |  |  | 11.2 ± 0.7 |  |  |
| groups | Ctrl | pCMBS | DIDS | Ctrl | pCMBS | DIDS |
| WBC<br>10<sup>3</sup>/μl | 3.3 ± 2.8 | 1.6 ± 0.6 | 2.2 ± 4.1 | 6.0 ± 4.6 | 2.3 ± 1.9* | 3.3 ± 1.8 |
| LYM<br>10<sup>3</sup>/μl | 2.6 ± 2.3 | 1.0 ± 0.4 | 1.7 ± 3.5 | 4.6 ± 3.9 | 1.6 ± 1.4* | 2.5 ± 1.4 |
| MONO<br>10<sup>3</sup>/μl | 0.2 ± 0.2 | 0.2 ± 0.1 | 0.2 ± 0.2 | 0.3 ± 0.2 | 0.2 ± 0.2 | 0.1 ± 0.1 |
| GRAN<br>10<sup>3</sup>/μl | 0.5 ± 0.4 | 0.4 ± 0.2 | 0.3 ± 0.4 | 1.0 ± 0.9 | 0.5 ± 0.4 | 0.6 ± 0.8 |
| LYM<br>% | 77.0 ± 3.9 | 62.4 ± 6.4* | 70.4 ± 8.4* | 78.0 ± 11.4 | 65.9 ± 12.0* | 77.1 ± 11.9 |
| MONO<br>% | 4.0 ± 1.1 | 9.7 ± 2.1* | 4.4 ± 0.7 | 3.9 ± 1.2 | 8.2 ± 2.7* | 3.5 ± 1.2 |
| GRAN<br>% | 19.0 ± 4.0 | 27.9 ± 5.4* | 25.2 ± 8.4* | 18.2 ± 11.4 | 25.8 ± 12.2 | 19.4 ± 11.2 |
| Hct<br>% | 33.1 ± 4.5 | 34.9 ± 3.4 | 35.4 ± 3.7 | 37.0 ± 6.0 | 39.1 ± 4.5 | 40.0 ± 4.0 |
| HGB<br>g/dl | 10.5 ± 1.1 | 11.8 ± 0.7* | 11.1 ± 0.9 | 11.3 ± 1.8 | 12.5 ± 1.3 | 12.1 ± 1.0 |
| RDW<sub>a</sub><br>fl | 29.7 ± 2.3 | 28.7 ± 1.9 | 31.7 ± 2.4 | 32.6 ± 1.4 | 31.8 ± 1.5 | 35.2 ± 1.7* |
| RDW<br>% | 16.8 ± 0.7 | 17.6 ± 1.1* | 17.3 ± 0.5 | 16.5 ± 0.5 | 17.1 ± 0.4* | 16.9 ± 0.5* |
| MCH<br>pg | 14.8 ± 0.4 | 15.0 ± 0.5 | 15.0 ± 0.5 | 15.3 ± 0.2 | 15.4 ± 0.3 | 15.6 ± 0.4 |
| MCV<br>fl | 46.4 ± 3.2 | 44.3 ± 3.0 | 47.5 ± 2.8 | 50.2 ± 0.8 | 48.2 ± 1.0* | 51.8 ± 1.3* |
| MCHC<br>g/dl | 32.1 ± 2.1 | 34.0 ± 2.1* | 31.6 ± 1.6 | 30.6 ± 0.4 | 32.0 ± 0.6* | 30.2 ± 1.0 |
| RBC<br>10<sup>6</sup>/μl | 7.1 ± 0.8 | 7.9 ± 0.6* | 7.4 ± 0.6 | 7.4 ± 1.2 | 8.1 ± 0.9 | 7.7 ± 0.7 |
| PLT<br>10<sup>3</sup>/μl | 91.7 ± 87.3 | 64.7 ± 18.3 | 85.9 ± 111.5 | 127.5 ± 113.2 | 68.0 ± 45.4 | 84.3 ± 56.0 |
| MPV<br>fl (N) | 6.4 ± 1.2<br>(9) | 6.1 ± 0.4<br>(12) | 6.2 ± 0.6<br>(11) | 6.2 ± 1.0<br>(13) | 6.1 ± 0.5<br>(11) | 7.7 ± 1.2*<br>(9) |

<sup>†</sup>These data on 13×2 = 26 mice are the hematology data obtained from mice used in the Inhibitor study, including those used in the imaging flow-cytometry inhibitor studies (see paper #2 in table 6). Because we observed no statistically significant differences between sexes in Table 4, we grouped a males and females (same gender for one pair of WT and dKO mice on that day) for this study.
Results are presented as mean ± SD. Each “N” represents an analysis of the blood from 1 mouse. Samples are tested at ostensibly 50% hematocrit (see Methods<sup>130</sup>). We analyzed data using a one-way ANOVA, followed by the Holm-Bonferroni (Holm, 1979) correction (see Methods<sup>131</sup>). For each parameter (e.g., MCV) from either WT or dKO, we made all possible comparisons between Ctrl, pCMBS and DIDS treatment. \* denotes a significant difference compared with Ctrl for the same genotype. See Supporting file 1 for access to *P*-values for all comparisons.
dKO, double knockout; WBC, white blood cells; LYM, lymphocytes; MONO, monocytes; GRAN, granulocytes; Hct, hematocrit; HGB, hemoglobin; RDW<sub>a</sub>, RBC distribution width–absolute; RDW%, RBC distribution width –percentage; MCH, mean corpuscular hemoglobin; MCV, mean corpuscular volume; MCHC, mean corpuscular hemoglobin concentration; RBC, red blood cells; PLT, platelet; MPV, mean platelet volume.

The RBCs summarized in Table 5 include those blood samples that we used for the IFC studies in paper #2^69^.

#### Morphometry

**Blood smears.** In <u>paper #2</u>^70^, we present two sets of blood-smear data. In the first, we examined control cells (i.e., those not exposed to pCMBS or DIDS) of all four genotypes. The pathologist read the RBC morphology (see <u>paper #2</u>^71^) as “unremarkable, demonstrating only occasional target cells and stomatocytes, with no differences noted among knockout strains and WT mice.”

In the second set of blood-smear data, we examined blood from (1) WT or (2) dKO mice pretreated with (a) neither pCMBS (controls), (b) with pCMBS, or (c) with DIDS (i.e., a total of 2 × 3 = 6). The pathologist read the RBC morphology (see <u>paper #2</u>^72^) as showing “no significant differences in red cell morphology … under any of the [six] conditions.” The absence of a comment about increased nBCDs prevalence in the drug-treated cells in the blood smears (where we plate, stain, and dry the cells after drug pretreatment) is interesting, inasmuch as nBCDs were apparent in images of living, tumbling RBCs or RBCs visualized by IFC. On the other hand, we made the blood smears immediately after drug treatment (e.g., <5 min), whereas the delays between drug treatment and live-cell imaging (see <u>below</u>^73^) were as long as 90 min.

**Living RBCs tumbling through the focal plane of an inverted microscope.** In paper #2, we also examine RBCs in suspension as we drop them onto a coverslip on the stage of an inverted microscope, and then image the cells as they fall through the focal plane (<u>paper #2</u>^74^). For control RBCs (no inhibitors present) of all four genotypes, the RBCs appear as normal biconcave disks, with occasional (∼1.35%) poikilocytes, both in still images (see <u>paper #2</u>^75^) and video recordings (see <u>paper #2</u>^76^).

In a second set of studies, we similarly examined RBCs—untreated, pretreated with pCMBS, or pretreated with DIDS—from WT or dKO mice. Whereas the appearance of the control RBCs (no inhibitor) was again unremarkable, we observed spherical cells (sometimes with spicules)—“nBCDs”—in the pCMBS experiments, and more so in the DIDS experiments (see <u>paper #2</u>^77^).

**Light-scattering flow cytometry.** As presented in <u>paper #2</u>^78^, these studies—which included the development of gating schemes (based on staining with fluorescent markers) to distinguish various cell types—showed that RhAG-KO and dKO mice tend to have slightly larger and more varied RBC sizes compared to WT and AQP1-KO mice. However, the overall differences in cell size and shape among the genotypes were not significant.

**Living RBCs visualized using imaging flow cytometry (IFC).** In a fourth and final approach for assessing RBC morphometry—and as presented in <u>paper #2</u>^79^—we establish gating schemes and then obtain photographs of single cells as they fly before a microscope objective.

*Cell shape.* We found that even control (not treated with inhibitors) RBCs from WT and dKO mice have ∼2% nBCDs (see <u>paper #2</u> ^80^). After pretreatment with pCMBS, the fraction increased significantly for the WT (∼8.7%) but less so in the dKO (∼5.7%). The pattern was the same following DIDS pretreatments, except that the fraction of nBCDs was far higher (41% for WT vs. 21% for dKO). The presence of nBCDs, particularly after drug pretreatment, led us to develop a novel approach to accommodate for these effects, as summarized <u>below</u>^81^.

As mentioned above, the time delays between completion of drug treatment and live-cell imaging (both DIC microscopy and IFC) were as long as 90 min, vs. the <5 min delay before either SF or preparation of blood smears.

The decrease in nBCD prevalence—both for samples pretreated with pCMBS or with DIDS (see previous paragraph)—in dKOs vs. WTs is consistent with the hypothesis that the drug-induced shape changes depend, at least in part, on interactions of the drugs with AQP1 and/or Rh_Cx_.

*Major RBC diameter (*Ø_Major_). The IFC experiments not only identify cell shape, but also provide—for cells with the proper orientation—the major diameter of BCDs and of nBCDs (see <u>paper #2</u>^82^). These values are an important input to the mathematical simulations (see <u>below</u>^83^).

##### Hb kinetics (*k*_HbO_2_→Hb+O_2__)

To examine the possibility that the knockout of one or both channels could affect the kinetics of HbO_2_ dissociation per se, we determined the rate constant of the reaction HbO_2_ → Hb + O_2_ in 100% hemolyzed RBCs. The approach was the same as for 100% ostensibly intact RBCs (see Figure 3 and Figure 4), or for various mixtures of 100% ostensibly intact and 100% hemolyzed RBCs (see Figure 7 and Figure 8). On the one hand, we find no significant differences among the mean *k*_HbO_2_→Hb+O_2__ values for WT, AQP1-KO, RhAG-KO mice (Figure 10). However, we are somewhat surprised—inasmuch as the individual AQP1 and RhAG deletions were without effect—that the dKO produced a ∼9% decrease of *k*_HbO_2_→Hb+O_2__ (i.e., 100% lysate).

**Figure 10.**
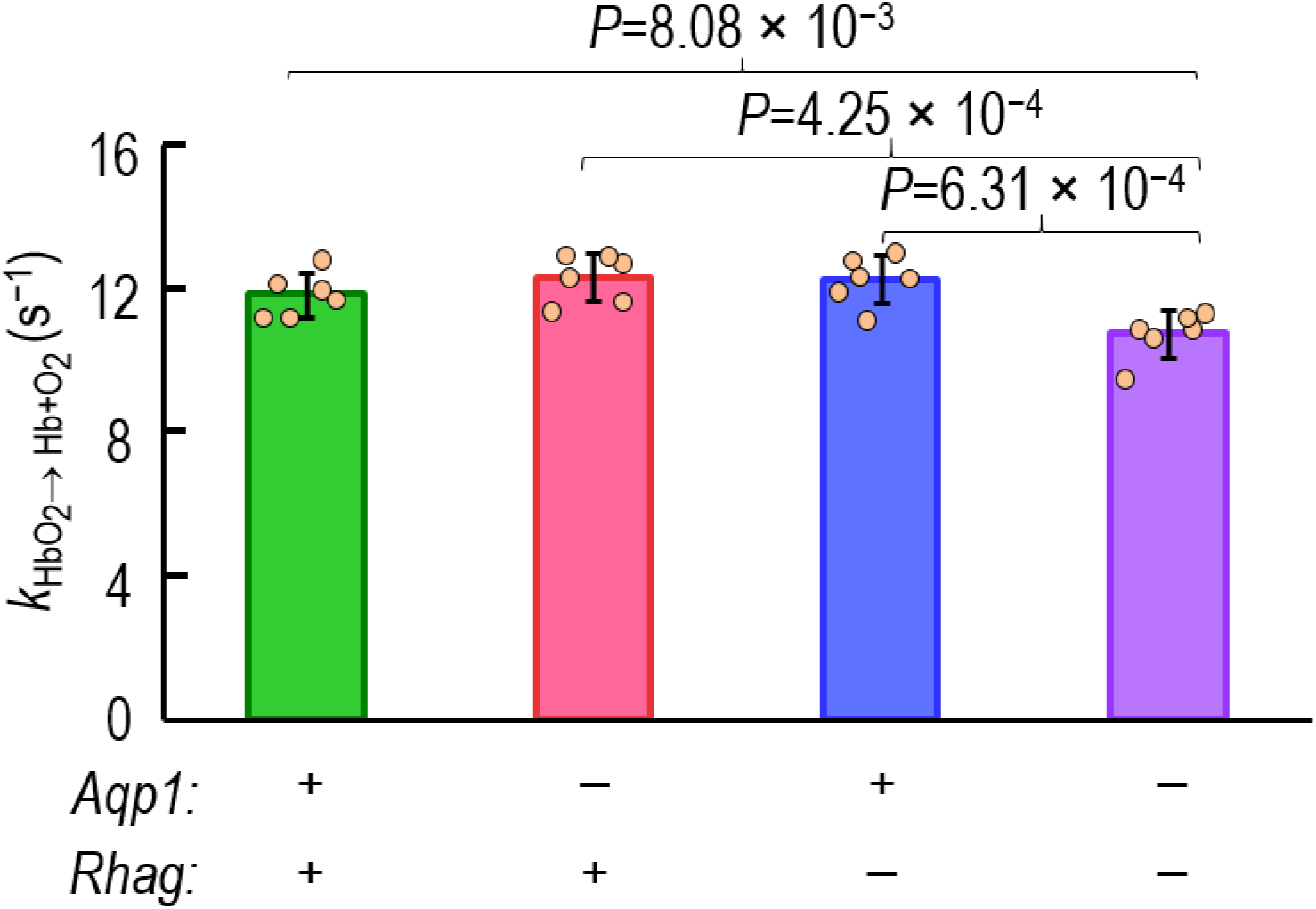
Effect of genetic deletions of *Aqp1*, *Rhag*, or both on rate constant of the reaction HbO_2_ → Hb + O_2_ (*k_HbO_2__→_Hb+O_2__*) in RBC lysates. See <u>Methods</u>^143^ for a description of how we measure *k*_HbO_2__. The approach here was similar, except that we worked with 100% hemolysates (see Methods^136^). Each bar-associated point represents RBCs from 1 mouse. Bars represent mean ± SD. N=6 for each bar. We performed a one-way ANOVA, followed by a Holm-Bonferroni correction (see <u>Methods</u>^135^). We present *P*-values for all statistical comparisons in Statistics Table 4.

Because the reaction HbO_2_ → Hb + O_2_ is only one of several processes that contribute to *k*_HbO_2__, intuition tells us that the ∼9% decrease in *k*_HbO_2_→Hb+O_2__ should account for even a smaller fraction of the ∼30% decrease that we observed in dKO mice. Indeed, the sensitivity analysis in <u>paper #3</u>^84^ indicates that a 9% decrease in *k*_HbO_2_→Hb+O_2__ would, by itself, lower *k*_HbO_2__ by only 1.5%. Viewed differently, in order to account, by itself, for the 30% decrease in *k*_HbO_2__ observed in physiological experiments, *k*_HbO_2_→Hb+O_2__ would have to fall by ∼63%.

### Mathematical simulations

**Overview.** Our goal in the macroscopic mathematical modeling and simulations is not to adjust simulation parameter values to produce a MMM-*k*_HbO_2__ (i.e., the simulated value) that matches the observed raw-*k*_HbO_2__ (e.g., the control curve for WT RBCs in Figure 3*C*), but rather to predict—from first principles—the time course of hemoglobin desaturation (see <u>paper #3</u>^85^). The numerical inputs for our simulations (see <u>paper #3</u>^86^) are physical constants and other values taken from the <u>literature</u>^87^ as well as <u>genotype-specific values</u>^88^ generated in the present paper and paper #2. Among the literature values, we took *P*_M,O_2__ to be 0.15 cm s^−1^ (Figure 1, workflow #13), the measured value of *P*_M,CO_2__ of Endeward *et al*. in human RBCs (Endeward *et al*., 2006). For the thickness of the unconvected layer, we chose 1 μm, which (given the high level of NTD in our experiments) may be an overestimate (Holland *et al*., 1985). Among genotype-specific values, the Ø_Major_ (workflow #6 and #6′, from <u>paper #2</u>^89^) and MCV (workflow #8 from <u>paper #2</u>^90^) are inputs for computing BCD thickness (workflow #10). MCH (workflow #7) as well as MCV (workflow #8) together yield MCHC (workflow #9). In the next major section, we summarize accommodations for nBCDs that yield shape-corrected values for *k*_HbO_2__.

**Summary.** The result of our simulation for control WT RBCs (i.e, without inhibitors)— all cells assumed to be 100% intact BCDs—is an MMM-*k*_HbO_2__ of ∼3.99 s^−1^ (workflow #14; see <u>paper #3</u>^91^). For the data set summarized in Figure 3*E*, the mean SC-*k*_HbO_2__ value underlying the control/WT value (darker green bar) is 3.45 s^−1^. For the (different) data set summarized in Figure 4*E* (green bar), the comparable value is 3.24 s^−1^, and for Figure 5*B* (green bar), the value is 4.09 s^−1^. Thus, our MMM-*k*_HbO_2__ value determined from first principles—3.99 s^−1^—ranges from ∼2.5% lower than the highest SC-*k*_HbO_2__ value to ∼23% higher than the lowest value. We conclude that the MMM simulation provides a reasonable estimate of HbO_2_ desaturation under the conditions of our experiments.

### Accommodation for nBCDs

Because it appears that all RBC preparations contain at least a small fraction of nBCDs, we developed a novel linear-combination method to accommodate for the presence of nBCDs (for summary, see <u>Methods</u>^92^; for detailed account, see <u>paper #3</u>^93^). This procedure, which corrects for the presence of “slower” nBCDs (lower *k*_HbO_2__) amidst the “faster” BCDs (higher *k*_HbO_2__), raises a hemolysis-corrected *k*_HbO_2__ value (e.g., our standard value of ∼3.99 s^−1^) to the corresponding shape-corrected value (i.e., ∼4.04 s^−1^ in this example). We can then compute a shape-correction factor *F*_SC_ (i.e., ∼ 4.04/3.99 ≅ 1.01), and apply it to any experiment on WT/Ctrl RBCs. Thus, if RBCs of the particular mouse exemplified in Figure 1 and Figure 7*A* have a raw-*k*_HbO_2__ of 3.70 s^−1^ (step #3 in Figure 1) and an HC-*k*_HbO_2__ of 3.45 s^−1^ (step #5), the linear-combination accommodation will yield an SC-*k*_HbO_2__ of ∼3.48 s^−1^ (step #22). This shape-correction factors is small with control RBCs (WT or various KOs) but becomes larger with pCMBS and especially DIDS.

## Discussion

### Overview

The combined results of the present paper and its two companion papers show that two channel proteins—AQP1 and the Rh complex—contribute ∼55% of the O_2_ permeability to murine RBCs. On the background of the AQP1/Rh_Cx_-dKO, we estimate that other protein(s) blocked by DIDS contribute an additional ∼21% of *P*_M,O_2__, whereas proteins blocked by pCMBS contribute ∼36%. If the DIDS-sensitive proteins (∼21%) are fully a subset of the pCMBS-sensitive ones (∼36%), then at least ∼91% of murine RBC O_2_ traffic occurs via membrane channels under the conditions of our experiments. To the extent that a subset of DIDS-sensitive proteins in the dKO are not also pCMBS sensitive, the contribution of unidentified channels would be greater than ∼36%, and the full collection of O_2_ channels would account for even more than ∼91% of O_2_ traffic in mice under the conditions of our experiments.

**Implications of O_2_ and CO_2_ channels.** Together with earlier CO_2_ studies, the present work shows that, of the O_2_ and CO_2_ traffic across mammalian RBC membranes, at least 90% occurs through protein channels. Our work raises questions that should trigger new lines of research: What is the molecular mechanism of O_2_ diffusion through the channels? We suggest that the most likely routes are the hydrophobic central pores at the middle of the AQP1 tetramers and Rh trimers, and/or other hydrophobic pathways between monomers. Which other membrane proteins contribute to *P*_M,O_2__ in RBCs? And to what extent do AQPs, Rh proteins, and other membrane proteins contribute to O_2_ permeability in other cell types? An important implication of our O_2_-channel work is that it provides a mechanism for cellular regulation of O_2_ permeability by controlling the synthesis/trafficking of these channels, or by using post-translational modifications or protein-protein interactions to modulate single-channel permeability. Our work raises the possibility of O_2_-channelopathies in human disease. Finally, the O_2_-channel paradigm offers the future physician the possibility of raising O_2_ permeability (e.g., to enhance performance or wound healing) or lowering O_2_ permeability (e.g., to treat oxygen toxicity) by altering the (1) number or (2) intrinsic permeability of channels.

**The Krogh cylinder.** A century ago, Krogh (1919) performed a mathematical analysis of O_2_ off-loading from RBCs, which included not only dissociation of O_2_ from HbO_2_, but also the diffusion of O_2_—lumped as a single process—through RBC intracellular fluid, across the RBC membrane to the bulk extracellular fluid of systemic capillary blood plasma, and then across the capillary endothelium, ultimately to the ICF of O_2_-metabolizing cells (see <u>paper #3</u>^94^).

At the time, Krogh’s analysis was a *tour de force* that blended state-of-the-art experimental physiology with mathematics. This mathematical approach, in the days before digital computational methods, required numerous simplifying assumptions to yield equations amenable to analytical solutions. Without these simplifying assumptions, Krogh’s quantitative model—with its demonstrated great value to physiology—would have been impossible. Sixty years after Krogh, Kreuzer (1982) enumerated 15 implicit assumptions in the Krogh analysis, one of which implies that the RBC membrane offers no resistance to O_2_ diffusion (i.e., *R*_M,O_2__ = 0). Whether or not Krogh himself would have agreed that *R*_M,O_2__ is, in fact, greater than 0, we suggest that his implicit assumption (i.e., *R*_M,O_2__ = 0) was a reasonable first step inasmuch as it enabled a mathematical analysis that provided the first quantitative insights into the Krogh cylinder. Nevertheless, this implicit assumption became dogma for at least a subset of physiologists. The present work, as discussed <u>below</u>^95^, proves that the implicit assumption is incorrect.

**Overton’s rule.** A larger subset of physiologists have adhered to a very different idea: Overton’s rule (a subset of the broader, previously proposed solubility-diffusion theory), which holds that the diffusion of substances across membranes depends exclusively on their solubility in the lipid phase of the membrane. At the time of Overton, the concept of a membrane protein’s mediating transport across a membrane would have been entirely foreign. For most of the panoply of substances that cross cell membranes—water, inorganic ions, small molecules, and even large substances like fatty acids and prostaglandins—Overton’s rule has long-since been abandoned in favor of channels and various transport proteins (for review, see Aronson *et al*., 2017). The present work with the knockout and blockade of channels (see <u>below</u>^95^)—together with previous work from this and other laboratories (Waisbren *et al*., 1994; Nakhoul *et al*., 1998; Cooper & Boron, 1998; Uehlein *et al*., 2003, 2008, 2012a, 2012b; Endeward *et al*., 2006, 2008; Musa-Aziz *et al*., 2007b, 2025; Geyer *et al*., 2013b, 2013a, 2013c; Nakhoul & Hamm, 2014; Wang *et al*., 2016, 2025; Zwiazek *et al*., 2017)—provides further evidence that the narrow view, attributed by others to Overton, does not hold universally for dissolved gases. Rather, *P*_M,O_2__ in murine RBCs depends largely (∼90%) on the presence and activity of at least three membrane proteins: AQP1, Rh_Cx_, and at least one unidentified protein.

### Automated hematology experiments

Our hematology assays (see Table 4 and Figure 9) did not reveal substantial differences among RBCs from mice of different genotypes, or of RBCs from WT or dKO mice treated with drugs. The critical values from the present paper for the mathematical simulations are MCV, MCH, and the derived MCHC (MCHC = MCH/MCV), all of which we passed along to paper #3 for simulations. As mentioned above and discussed in detail <u>below</u>^96^, the KO-specific changes in MCV and MCHC tend to cancel out, leading to the conclusion that automated-hematology data cannot account for the genotypic effects on *k*_HbO_2__.

In parallel with the two critical hematological parameters (MCV and MCHC), major diameter and nBCD prevalence pass in parallel from paper #2 to paper #3, providing the final values required for the mathematical simulations and accommodations.

### Stopped-flow experiments

**Hemolysis.** We developed our SF hemolysis assay (Zhao *et al*., 2017), which is based on the release of CA from RBCs into the extracellular fluid, because we anticipated that some RBCs lyse within the SF reaction cell and thereby lead to an overestimate of *k*_HbO_2__. In the present paper, we develop what we believe are the first approaches (Figure 7 and Figure 8) for accommodating for the hemolysis that must always be occurring during SF studies. In brief, we artificially create precise degrees of incremental hemolysis, measuring *k*_HbO_2__ at each. As predicted, *k*_HbO_2__ rises as apparent hemolysis rises. After a rescaling and translation to create an x-axis that represents actual hemolysis, a back-extrapolation yields the true *k*_HbO_2__ when actual hemolysis should be zero. Because this hemolysis-corrected *k*_HbO_2__ is always less than the raw-*k*_HbO_2__, the fractional inhibition based on raw-*k*_HbO_2__ values will underestimate those based on HC-*k*_HbO_2__ values.

In the case of control RBCs (not treated with a drug), these hemolysis corrections are modest, in the range of 4.0% to 7.0% for WTs, ∼3.4% for AQP1-KOs, ∼5.3% for RhAG-KOs, and ∼9.8% to ∼10.7% for dKOs (see Table 3). For drug-treated cells, the hemolysis corrections are larger, ∼17% (pCMBS) and ∼25% (DIDS) for WTs and 41% (pCMBS) and 33% (DIDS) for dKOs. The drug values are larger than their control counterparts because (1) the actual % hemolysis is larger than for control cells (see Figure 6) so that the extrapolation is somewhat greater and (2) drug-treated cells have lower raw-*k*_HbO_2__ values so that even the same absolute hemolysis correction produces a greater fractional change.

**Non-biconcave disks (nBCDs).** We developed our accommodation for nBCDs because we observed them in DIC images of living drug-treated RBCs (see <u>paper #2</u>^97^). Imaging flow-cytometry confirms that nBCDs—a mixture of poikilocyte types (mainly echinocytes, acanthocytes, and spherocytes)—appear in all of our preparations (see <u>paper #2</u>^98^)—at least after the prolonged times needed to execute these assays. The prevalence (see <u>paper #2</u>^99^) was very low in control RBCs from WT and dKO mice (∼2%), larger with pCMBS pretreatment (∼9% in WT, falling to ∼6% in dKOs), and high in DIDS-treated cells (∼41% in WT, falling to ∼21% in dKOs). Because the delay between drug treatment and IFC experiments is greater (∼30 to 90 min) than between drug treatment and SF experiments or preparation of blood smears (∼5 min), the nBCD prevalence from IFC studies probably overestimates the actual prevalence in SF studies, and thus probably leads to underestimates of the true drug effects on *P*_M,O_2__.

To our knowledge, our IFC study is the first quantitative search in physiological experiments for nBCDs, which nevertheless were presumably present in previous studies. In future studies, it would be valuable to determine the interrelationships among (1) drug concentration and exposure time, (2) decrease in *P*_M,O_2__, and (3) nBCDs prevalence, with the goal of determining protocols that maximize inhibition while minimizing nBCDs formation.

In the present study, we accommodated for nBCDs by (1) using IFC data to obtain diameter and prevalence, (2) assuming that the nBCDs are spheres, (3) assuming that membrane properties (e.g., *P*_M,O_2__) and MCH (i.e., total amount of Hb/cell) are the same in BCDs and nBCDs, (4) using our mathematical simulations to make a provisional estimate of the *k*_HbO_2__ of nBCDs (always lower than for BCDs), and (5) applying our linear-combination approach (see <u>paper #3</u>^100^) to extract the *k*_HbO_2__ for BCDs (always higher than HC-*k*_HbO_2__).

Note that although we assume that nBCDs are perfect spheres (minimal surface/volume ratio, slowest HbO_2_ offloading), in fact nBCD surface geometry is often more complex (higher surface/volume, faster offloading). Thus, our *k*_HbO_2__ of nBCDs probably tends to be an underestimate, so that our linear-combination approach tends to over-correct and yield a deduced *k*_HbO_2__ of BCDs—the SC-*k*_HbO_2__—that overestimates the true *k*_HbO_2__ of BCDs. As a result, when computing fractional inhibition, we tend to underestimate very slightly the effect of channel KOs under control (i.e., drug-free) conditions, and underestimate more substantially the inhibition produced by pCMBS and particularly DIDS.

In the case of control RBCs (not treated with a drug), both WT and KO, the shape-change correction factors—probably slight overestimates both because of the likelihood of our overestimating nBCD prevalence and underestimating nBCD surface-to-volume ratio of nBCDs— are small, *F*_SC_ ≅ 1%. In the case of drug-treated cells, the *F*_SC_ values (probably greater overestimates) are larger, ∼6% for pCMBS and ∼35% for DIDS. To the extent that we overestimated *F*_SC_, we underestimated the inhibitory effects of the drugs, and underestimated the total fraction of transmembrane O_2_ traffic attributable to membrane proteins under the conditions of our work. Thus, this fraction could well exceed our 91% estimate.

**Overall assessment of accommodations.** For control RBCs of all genotypes, the rate of hemolysis is small (*Act%H* ≅ 5%; see Figure 6*F*), the *k*_HbO_2__ correction for hemolysis is similarly small (∼4% to ∼6%; see Table 3*B*), nBCDs prevalence is low (∼2%; see <u>paper #2</u>^101^), and the additional correction for nBCDs is similarly small (*F*_SC_ ≅ 1%; see Table 3*B*). We regard both our hemolysis assay and hemolysis correction as robust. Even so, the correction is likely to be most reliable with low *Act%H*. The shape-change correction is likely most reliable with low nBCD prevalence. Thus, we have our highest degree of confidence in our control data on RBCs from WT, AQP1-KO, RhAG-KO, and dKO mice—that is, the dKO likely reduces *P*_M,O_2__ by a substantial amount (best estimate ≅ 55%; see <u>paper #3</u>^102^).

*pCMBS.* For WT and dKO cells pretreated with pCMBS, hemolysis is still modest (*Act%H* ≅ 8% and 11%, respectively; see Figure 6*E* and *G*), the *k*_HbO_2__ hemolysis correction is modest to large (about –30% and –41%, respectively; see Table 3*A* and *C*), nBCDs prevalence is modest to small (∼9% and ∼6%, respectively; see <u>paper #2</u>^101^ and the correction for nBCDs is small (about +6% for WT, +4% for dKO; see Table 3*A* and *C*). In our favor, the hemolysis and shape corrections are in opposite directions. Thus, we are confident that, in RBCs from WT mice, pCMBS substantially reduces both *k*_HbO_2__ (best estimate ≅ 61%; see Figure 3*E* and <u>paper #3</u>^102^) and *P*_M,O_2__ (best estimate ≅ 82%; see <u>paper #3</u>^102^). On a dKO background, we have greater confidence (because of a smaller *F*_SC_) that pCMBS produces a further decrease in *k*_HbO_2__ beyond the dKO effect (best estimate ≅ 68%; see Figure 5*D*). Thus, we are confident that the combination of dKO and pCMBS produces an even-larger total reduction in *k*_HbO_2__ (best estimate ≅ 78%; see Figure 5*D* and <u>paper #3</u>^102^). This last value translates to a maximal reduction in *P*_M,O_2__ of ∼91% (see <u>paper #3</u>^102^).

*DIDS.* WT and (to a lesser extent) dKO cells pretreated with DIDS are the most problematic in the present study. Hemolysis is rather high (*Act%H* ≅ 13% and 15%, respectively; see Figure 6*E* and *G*), the *k*_HbO_2__ hemolysis corrections are high (about –31% and –33%, respectively; see Table 3*A* and *C*), nBCD prevalences are high (∼41% and ∼21%, respectively; see <u>paper #2</u>^101^), and the correction for nBCDs is high (about +35% for WT, +16% for dKO; see Table 3*A* and *C*). The corrections are large, but they are similar in magnitude but opposite in direction. As a result, we are confident that, in RBCs from WT mice, DIDS reduces both *k*_HbO_2__ (best estimate ≅ 31%; see Figure 4*E* and <u>paper #3</u>^102^) and *P*_M,O_2__ (best estimate ≅ 56%; see <u>paper #3</u>^102^). On a dKO background, we have somewhat greater confidence (again, because of a smaller *F*_SC_) that DIDS produces a further decrease in *k*_HbO_2__ beyond the dKO effect (best estimate ≅ 32%; see Figure 5*F*). Thus, we are confident that the combination of dKO and DIDS produces an even-larger total reduction in *k*_HbO_2__ (best estimate ≅ 53%; see Figure 5*F* and <u>paper #3</u>^102^). This last value translates to a maximal reduction in *P*_M,O_2__ of ∼76% (see <u>paper #3</u>^102^).

Although we believe that our data show that DIDS blocks a major component of *P*_M,O_2__ in murine RBCs under the conditions of our experiments, we appreciate that one might be concerned that DIDS-treated RBCs have a rather high values for *Act%H* and a nBCD prevalence—even if we have attempted to accommodate for these effects. Nevertheless, even if we were to ignore the DIDS data in the present paper, we would still come to the conclusion that the dKO reduces *P*_M,O_2__ by ∼55%, and that pCMBS blocks an additional ∼36%, to take the total protein-mediated component of *P*_M,O_2__ to at least ∼91%.

**Mechanism of drug action.** Our work shows that neither pCMBS (see Figure 3*E*) nor DIDS (Figure 4*E*) gains significant entry to the RBC cytosol. Thus, the effects of pCMBS (a sulfhydryl reagent) and DIDS (an amino-reactive agent) must be at the level of the exterior surface of the plasma membrane. Other authors have presumed that both drugs—neither of which is specific for one target—act on membrane proteins (Rothstein *et al*., 1973; Cabantchik & Rothstein, 1974), and we know of no evidence that either interacts with membrane lipids (Vansteveninck *et al*., 1965; Rothstein *et al*., 1973; Cabantchik & Rothstein, 1974). Moreover, if either did interact with membrane lipids, we expect that the result—if any—would be to decrease the barrier to O_2_ diffusion. Thus, the inhibitor data led us to explore channel KOs.

**Effects of KOs on drug susceptibility.** Although neither pCMBS nor DIDS is specific for a single target, AQP1 and Rh_Cx_ appear to be targets of each. Macey and Farmer (1970) were the first to show that mercurials reduce the water permeability of RBCs (for early review, see Macey, 1984). After the identification of AQP1 as the first H_2_O channel, Preston *et al*. (1993) created the C189S mutation and demonstrated that the inhibitory effect of mercury on the H_2_O permeability of AQP1 depends specifically on the interaction with Cys-189 (Preston *et al*., 1993). Moreover, the mutation AQP1-C189S eliminates the ability of pCMBS to reduce *P*_M,CO_2__ (Cooper & Boron, 1998). Preliminary work suggests that DIDS also reduces the CO_2_ permeability AQP1, as expressed in oocytes (Musa-Aziz *et al*., 2007b, 2007a, 2025).

In the present study, our mathematical simulations suggest that, on a WT background, pCMBS reduces *P*_M,O_2__ by ∼82% (see <u>paper #3</u>^103^). On the other hand, on a dKO background, the drug decreases *P*_M,O_2__ only by ∼91% – ∼55% = ∼36% (i.e., inhibition is reduced by 56% on dKO background). For DIDS, the two degrees of inhibition for SC-*k*_HbO_2__ are ∼56% on a WT background, and ∼76% – ∼55% = ∼21% on a dKO background (i.e., inhibition is reduced by 63% on dKO background). Thus, the combined absence of AQP1 and Rh_Cx_ reduces the susceptibility of *P*_M,O_2__ to each drug by a substantial amount. We have already seen that (compared to a WT background) the dKO background reduces the susceptibility of RBCs to nBCD formation (see <u>paper #2</u>^104^). Thus, both from the perspective of *P*_M,O_2__ reduction and nBCD formation, both AQP1 and Rh_Cx_ appear to be major—although not the exclusive—targets of both pCMBS and DIDS.

### Implications of morphometry and proteomics experiments

**Morphometry.** We were relieved to see that, for all three KO genotypes, the control cells—as for the WTs—were overwhelmingly shaped as biconcave disks (see <u>paper #2</u> ^105^). Therefore, on the basis of RBC morphology alone, we conclude that we have no reason to be suspicious of the genotype-dependent decreases in SC-*k*_HbO_2__ and inferred *P*_M,O_2__ and the deduction—independent of any inhibitor data—that AQP1 and Rh_Cx_ can be conduits for O_2_.

In the absence of inhibitors, nBCD prevalence detected by IFC in WTs and dKOs was <2% (see <u>paper #2</u>^101^); our linear-combination accommodations ought to be most reliable for these control cells (see <u>paper #3</u>^106^). For cells treated with pCMBS, nBCD prevalence is still modest and we believe that our accommodation (i.e., SC-*k*_HbO_2__) is likely to be nearly as reliable as for control cells. For cells treated with DIDS, which produces smaller *k*_HbO_2__ inhibitory effects than does pCMBS but a much larger nBCD prevalence, we have a lesser degree of confidence in the accuracy of SC-*k*_HbO_2__ and deduced the *P*_M,O_2__. Nevertheless, even if we were to ignore the DIDS data, the overall conclusions of our three papers would be unchanged. That is, besides AQP1 and Rh_Cx_ (the support for which comes from KO data on control cells), we conclude that murine RBCs have at least one additional pCMBS-sensitive conduit accounts for ∼36% of *P*_M,O_2__. If we accept the DIDS SC-*k*_HbO_2__ values at face value, murine RBCs have—in addition to AQP1 and Rh_Cx_—at least one additional DIDS-sensitive conduit accounts for ∼21%. As noted at the outset of the Discussion, the 21% could overlap completely or partially with the 36%.

Measurements of major diameter and nBCD prevalence flow from paper #2 paper #3 for mathematical simulations and accommodations. Because of the large number of cells on which we made measurements by IFC (<u>paper #2</u>^107^), we have high confidence in both measurements.

**Proteomics.** Our proteomics analysis shows that the genetic deletions of AQP1 and RhAG appropriately eliminated those proteins (plus RhD in the case of the RhAG-KO), but had no significant effect on the apparent abundance of any other proteins among those ranked in the top 100 of greatest inferred abundance (see <u>paper #2</u> ^108^). Thus, the most straightforward explanation for our genotype-base data on RBCs is that it was the absence of AQP1 or RhCx per se that was responsible for the KO-induced decreases in SC-*k*_HbO_2__ and deduced *P*_M,O_2__.

### Implications of the macroscopic mathematical model

#### Basic features of the model

**The sphere.** We model the RBC as a sphere, the diameter of which is the same as the calculated thickness of the RBCs used in physiological experiments. Thus, our approach emphasizes the free edge of the BCD, which contains most of the HbO_2_. Others also have regarded BCD thickness as the dimension most relevant for diffusion (Roughton, 1932; Ponder, 1948; Forster, 1964; Endeward, 2012).

**Simulated *k*_HbO_2__ for WT/Ctrl RBCs.** Our simulations—based on physical constants and values provided either by the literature or our own measurements on control RBCs from WT mice—produce an SC-*k*_HbO_2__ value of 4.04 s^−1^, which is within the range of our physiologically observed values obtained in different data sets (i.e., 3.24 s^−1^ in Figure 4*E*, 3.45 s^−1^ in Figure 3*E*, 4.09 s^−1^ in Figure 5*D*. We believe that this range of physiologically obtained SC-*k*_HbO_2__ values reflects variations in the potency of the manufacturer-supplied NDT supply, with low potency (i.e., lower effective [NDT]) generating a greater effective unconvected layer around the RBCs (Vandegriff & Olson, 1984b), and thus a lower apparent *k*_HbO_2__ value (see <u>paper #3</u>^109^). We presume that our highest-observed SC-*k*_HbO_2__ value reflects our most potent batch of NDT^110^ and thus our most reliable estimate of a reference SC-*k*_HbO_2__ for WT/Ctrl RBCs of our C57BL/6_Case_ mouse strain. This value of 4.09 s^−1^ also is closest to our simulated value of 4.04 s^−1^.

In our studies with pCMBS (e.g., see Figure 3) and with DIDS (e.g., see Figure 4), we always examined RBCs, ±inhibitor, from the same mouse on the same day, thereby minimizing effects of NDT potency.

**Predictions for effects on *P*_M,O_2__.** Starting with the simulation in which we assumed a provisional *P*_M,O_2__ of 0.15 cm s^−1^—this yielded the provisional MMM-*k*_HbO_2__ value of 3.99 s^−1^ noted above (see also <u>paper #3</u>^111^)—we systematically varied *P*_M,O_2__ in a series of nearly 75 simulations. The result is a sigmoidal dependence of *k*_HbO_2__ on log(*P*_M,O_2__), as shown in <u>paper #3</u>^112^. The highest simulated *P*_M,O_2__ is ∼22.6 cm s^−1^, which is near the asymptote of SC-*k*_HbO_2__ ≅ 6 s^−1^, corresponds to a situation in which we replace the membrane with an equivalent thickness of water. Thus, simulating a physiological SC-*k*_HbO_2__ of 3.99 s^−1^ requires that we reduce the *P*_M,O_2__ of a “water membrane” by a factor of (22.6/0.15) ≅ 150. These data are inconsistent with the implicit assumption of Krogh that membranes, including RBC membranes, offer no resistance to O_2_ diffusion. In fact, RBC membrane, even with a full complement of O_2_ channels, offer substantial resistance to O_2_ diffusion, and this resistance increases even further with the genetic deletion of channels and/or treatment with inhibitors (<u>paper #3</u>^113^).

**Ability of inhibition of *P*_M,O_2__ to decrease *k*_HbO_2__.** Figure 11 is a transformation of the data in the aforementioned sigmoidal curve in <u>paper #3</u>^112^. Here in Figure 11 we show how the computed “% decrease of SC-*k*_HbO_2__” depends on the imposed “% decrease in *P*_M,O_2__”. At the lower left, the WT/Ctrl point confirms that 0% decrease in permeability (*P*_M,O_2__ = 0.1546 cm s^−1^) yields 0% decrease of SC-*k*_HbO_2__ (*k*_HbO_2__ = 4.04 s^−1^). The point at the upper right confirms that a 100% decrease in permeability (*P*_M,O_2__ = 0) fully blocks O_2_ offloading (*k*_HbO_2__ = 0). Between these two extremes are points that represent key results of the present paper. For example, simulating the ∼30% decrease in SC-*k*_HbO_2__ that we observe with dKO/Ctrl RBCs requires that we reduce the *P*_M,O_2__ from its reference value of 0.1546 cm s^−1^ by 55%. Replicating the ∼78% decrease in SC-*k*_HbO_2__ that we observed for dKO/pCMBS requires that we reduce *P*_M,O_2__ by 91%. If O_2_ permeability depended only on O_2_ solubility in the lipid phase of the membrane, as required by Overton’s rule, then the seven experimental maneuvers (from AQP1-KO/Ctrl near the lower left to dKO/pCMBS near the upper right) would have been without effect on either *P*_M,O_2__ or *k*_HbO_2__, and all the points in Figure 11 would overlie one another at the origin. This is clearly not the case. Thus, Overton’s rule, at least as applied to the O_2_ permeability of our murine RBCs under our experimental conditions, must be false.

**Figure 11.**
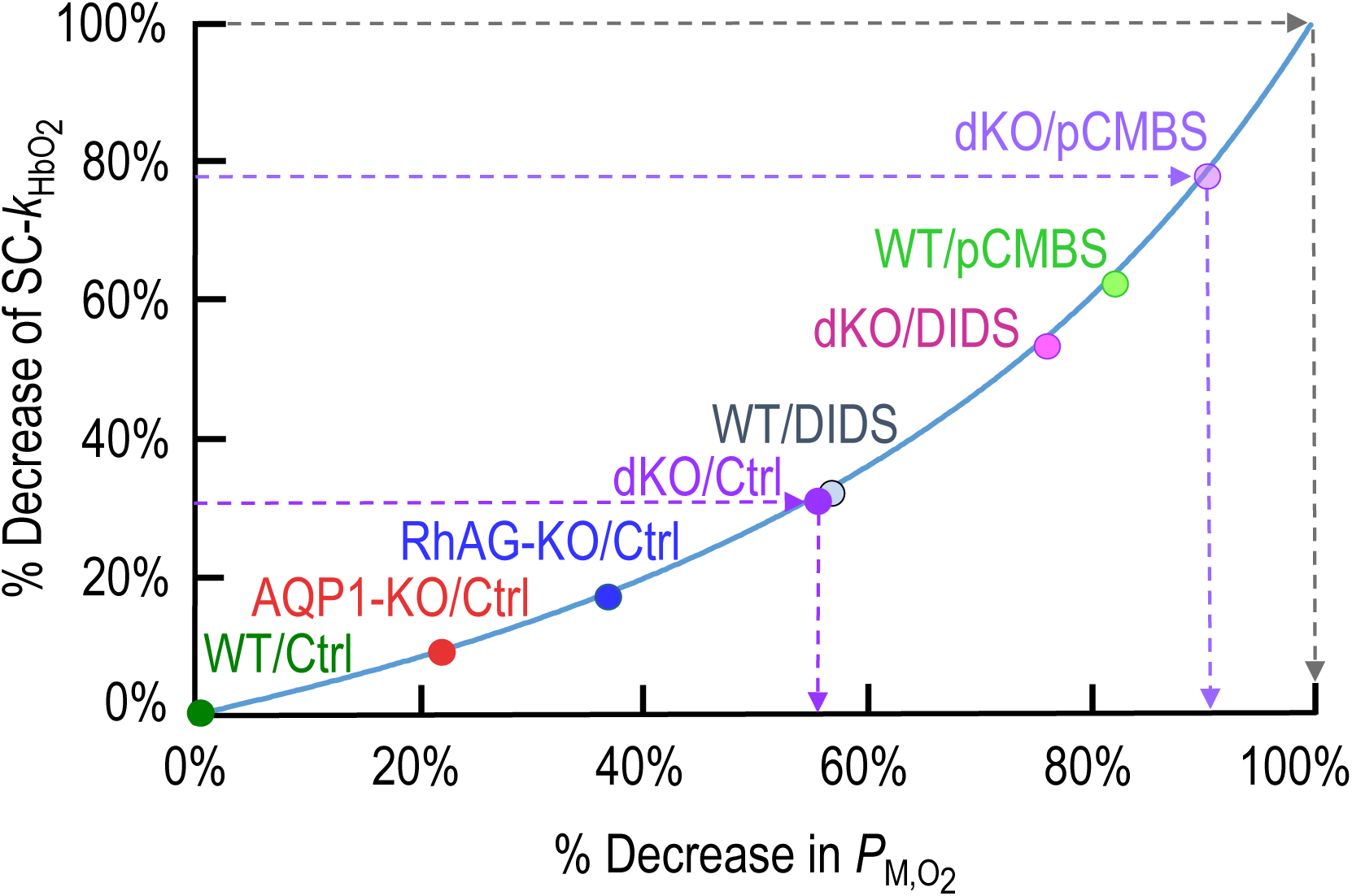
Dependence of percent decrease of *k*_HbO_2__ on percent decrease of *P*_M,O_2_ from_ macroscopic mathematical modeling and simulations. Effect of decreasing the O_2_ permeability of the RBC membrane (*P*_M,O_2__) on the predicted shape-corrected (SC) *k*_HbO_2__. We reference both the % decrease in *k*_HbO_2__ (equivalent to the shape-corrected value of a physiological experiment) and the % decrease in *P*_M,O_2__ (equivalent to the inhibition produced by a KO and/or drug) to their respective values for control (Ctrl; i.e., without inhibitors) RBCs from a wild-type (WT) mouse. Note: this plot is a transformation of the sigmoidal MMM-*k*_HbO_2__ vs. log(*P*_M,O_2__) modeling/simulation data in figure 6 <u>in paper</u> <u>#3</u>. Using our reaction-diffusion model (see <u>paper #3</u>^144^), we simulate the time course of HbO_2_ desaturation, employing hematological and morphological data gathered for RBCs from WT mice. We then systematically decrease *P*_M,O_2__ (x-axis), and compute the predicted % decrease in *k*_HbO_2__ (y-axis; blue curve). For each of our 8 experimental conditions (labeled points), we use experimentally determined SC-*k*_HbO_2__ values to determine position on the y-axis, and then place the point on the blue curve. Dropping a perpendicular to the x-axis reveals the % decrease in *P*_M,O_2__. Thus, the observed ∼78% decrease in SC-*k*_HbO_2__ for dKO/pCMBS (middle horizontal arrow) corresponds to a ∼91% decrease in *P*_M,O_2__ (middle downward arrow).

**Additivity of *P*_AQP1,O_2__ and *P*_RhCx,O_2__.** One way of evaluating the internal consistency of our macroscopic mathematical model and resulting simulations is to ask whether the decreases in *P*_M,O_2__ predicted in Figure 11 for the individual channel knockouts sum to the same value as for the double knockout

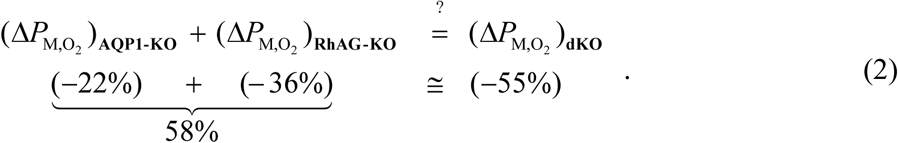

The agreement is good, with the individual KOs summing to a predicted *P*_M,O_2__ decrease of ∼58%, slightly above the observed dKOs value of ∼55%. We conclude that the model, as tested by physiological data, is well behaved.

#### Sensitivity analysis

<u>Paper #3</u>^114^ presents an analysis of the sensitivity of the simulated *k*_HbO_2__ to changes in eight key kinetic and geometric parameters. Our general conclusion is that no reasonable change in any parameter could account for our *k*_HbO_2__ data on control RBCs from KO mice, let alone from inhibitor-treated RBCs from dKO mice. More specifically, *k*_HbO_2__ is: (1) only modestly sensitive to *k*_HbO_2_→Hb+O_2__, which we measure; (2) only mildly sensitive to changes in [Hb_Total_]_i_, which also is a measured parameter; (3) modestly sensitive to changes in O_2_ diffusion constants and virtually insensitive to changes in HbO_2_ and Hb diffusion constants; and (4) moderately sensitive to changes in either the thickness of the unconvected layer or (5) the diameter of the sphere that, in the MMM, emulates Ø_Major_. Because the simulations deal with a single cell with no extracellular hemoglobin, the results are, by definition, both hemolysis- and shape-corrected.

#### Effect of channel knockouts on MCV and MCHC

In our broadest automated-hematology data set (see Table 4), the knockouts involving RhAG (RhAG-KO and dKO) were associated with a statistically significant increases in MCV and reciprocal decreases in MCHC; the AQP1-KO data trended in the same direction. In a narrower data subset restricted to mice used in IFC studies—and then in mathematical simulations—these patterns of MCV and MCHC persisted as trends but did not reach statistical significance (see <u>paper</u> <u>#2</u>^115^).

**MCV.** Our imaging–flow-cytometry studies indicate that the aforementioned increases in MCV are mainly attributable to an increased prevalence of immature cell types (see <u>paper #2</u>^116^), consistent with increased RBC production. Because both AQP1 and Rh_Cx_ are important components of the ankyrin complex (Vallese *et al*., 2022), one might think that the deletion of one or both channels might impact RBC deformability, which in turn could lead to a decrease in average RBC lifespan, consistent with our observation of increased immature cell types. One might also think that the deletions could increase RBC fragility. However, we did not observe genotype-related differences in hemolysis within the SF reaction cell (see Figure 6*F*).

**MCHC.** Whereas the genetic deletion of one or both channels tends to raise MCV, the effect on MCH is virtually nil (see Table 4). As a result, MCHC—that is, the ratio MCH/MCV— tends to fall as MCV rises. The parameter-sensitivity analysis in <u>paper #3</u>^114^ indicates that, all else (including [Hb_Total_]_i_) being equal, increases in the sphere diameter (a surrogate for RBC thickness, which is related to MCV) used in the mathematical model causes MMM-*k*_HbO_2__ to fall rather <u>steeply</u>^117^. On the other hand, the sensitivity analysis also shows that decreases in [Hb_Total_]_i_, all else (including sphere diameter) being equal, causes MMM-*k*_HbO_2__ to rise rather <u>steeply</u>^118^. Intuitively, one might think that the two effects—↑MCV and ↓MCHC—would tend to cancel, a hypothesis that we address in the next paragraph.

**Simulations based on genotype-specific data.** To address the above hypothesis, in paper #3 of this series, we perform a set of mathematical simulations in which we mimic control cells (i.e., not pretreated with inhibitors) of all four genotypes. The input parameters are genotype-specific values of MCV, MCHC, Ø_Major_. This <u>analysis</u> ^119^ predicts that the AQP1-KO would actually increase SC-*k*_HbO_2__ by ∼1% to 3% (vs. the observed 9% decrease in Figure 5*B*). The simulated SC-*k*_HbO_2__ for the RhAG-KO is down by ∼4% from WT vs. the observed ∼17% decrease, and that for the dKO is down by ∼4% to ∼6% from WT vs. the observed ∼30% decrease. We conclude that the effects of anti-parallel (1) increases in MCV and (2) decreases in MCHC do indeed tend to cancel, and that variations in RBC dimensions and [Hb_Total_]_i_ cannot account for our SC-*k*_HbO_2__ data among the four genotypes in the present study.

### Reports on O_2_ channels

**Yeast expression system.** In experiments on protoplasts from the yeast *Saccharomyces cerevisiae*, (Zwiazek *et al*., 2017) used the absorbance spectrum of myoglobin (Mb) to assess O_2_ influx. They found that expression of NtPIP1;3 from *Nicotiana tabacum*—but not that of several other plant AQPs, including AtPIP1;2 from *Arabidopsis thaliana*—substantially and significantly increased the rate of O_2_ uptake (as reported by the Mb absorbance spectrum) vs. mock transfection. Interestingly, the AtPIP1;2 rate trended below “mock”, perhaps an example of a “blocking protein”—a non-conducting membrane protein that reduces available surface area of the lipid bilayer—or the result of reduced expression of other O_2_ channels. The rate of human AQP1 was modestly but significantly greater than that of AtPIP1;2.

The authors did not translate changes in absorbance spectrum to changes in myoglobin O_2_ saturation or O_2_ fluxes, as in Figure 3*D*. Nor did they consider other components of the system that would have impacted rates of O_2_ influx (the opposite direction of those shown in Figure 2): (1) depletion of O_2_ in the EUF, especially in the absence of an extracellular O_2_ donor; (2) diffusion of O_2_, Mb, and MbO_2_ through the intracellular fluid; or (3) the association-reaction rate constant *k*_Mb+O_2_→MbO_2__, analogous to our dissociation-reaction rate constant *k*_HbO_2_→Hb+O_2__, as measured in Figure 10.

**Murine RBCs.** In a preprint, Zhao *et al*. (2020) reported the essential elements and conclusions of the present paper, though without the accommodation for nBCDs that we implement in the present series of papers.

**Human and murine RBCs.** In a recent paper, Al-Samir *et al*. (2025) applied insights from the Zhao preprint to work on RBCs, ±DIDS, from both WT and AQP1-KO mice as well as normal and Colton-null (*Aqp1*–/–) humans. They also extended the temperature from 7 or 10 °C to 25 °C, and 37 °C. We note several differences between their study and ours:

*Mouse strains.* Al-Samir *et al*. used “C57BL/6” mice. Kelmenson (2016) has noted that BL/6 strains can differ substantially. We used “C57BL/6_Case_”, which we characterize (see Table 1).

*Study design.* Our murine study is more expansive in that we include the second inhibitor pCMBS, as well as the strains RhAG-KO and dKO. Whereas Al-Samir *et al*. used WT mice at ages from ∼ 3 to ∼18 months, and AQP1-KO mice at ages ∼4 to ∼16 months, in the present study, we used mice (all genotypes) between ∼2 and ∼4 months of age. Although not a part of the present study, preliminary work shows that *k*_HbO_2__ values of WT/Ctrl RBCs are indistinguishable from ages 2 to 3 months through 12 months of age, but begins to fall at ∼12 months, decreasing by ∼13% at age 29 months (i.e., the lifespan of our C57BL/6_Case_ mice; Zhao & Boron, 2019, Zhao *et al*., 2021). Thus, some of the data of Al-Samir *et al*. could have been confounded by aging.

*Automated hematology*. Al-Samir *et al*. reports MCV on four mice of each genotype/sex. We report a full hematological analysis on each mouse and used the results—along with Ø_Major_ measured by imaging flow cytometry—to compute RBC thickness and [Hb_Total_]_i_, both of which we use in our simulations and estimation of *P*_M,O_2__.

*Hemolysis accommodation*. Al-Samir *et al*. collected effluent from the reaction cell after multiple SF shots, centrifuged, estimated free [Hb], and applied a correction. This approach likely overestimated actual hemolysis because, after the *k*_HbO_2__ assay in the reaction cell, additional RBC lysis is likely to have occurred. Indeed, their reported % hemolysis levels were 3-fold greater than ours. Assuming that their actual % hemolysis values were similar to ours, their overestimates of *Act%H* would have led to an overcorrection and thus lower HC-*k*_HbO_2__ values. If we had applied such underestimated HC-*k*_HbO_2__ values to compute % inhibitions due to DIDS or AQP1-KO (see Figure 8 and Table 3), we would have generated large overestimates of % inhibition.

We use a CA-based assay (Zhao *et al*., 2017) that reports *Act%H* within the SF reaction cell, based on single “shots”. Although this approach is time consuming, it assesses hemolysis under the conditions of the *k*_HbO_2__ determinations. Moreover, our operationally defined correction (see Figure 8) is based on a series of *k*_HbO_2__-like measurements over a range of imposed degrees of hemolysis and thus is free of theoretical assumptions.

*Poikilocytes.* Al-Samir *et al*. noted “highly irregular shaped” RBCs in their confocal observations of “almost all” DIDS-treated murine cells but neither quantitated nBCD prevalence nor accommodated for it.

In the present papers, we report the statistics of studies in which we used IFC to measure nBCD prevalence among hundreds of thousands of cells among various protocols (see <u>paper</u> <u>#2</u>^120^). Moreover, we used these prevalences to correct HC-*k*_HbO_2__ values (see <u>paper #3</u>^121^). If we had ignored the shape correction in the case of WT/DIDS, the result would have been a large underestimate of the true *k*_HbO_2__, and thus a large overestimate DIDS-induced inhibition (see Table 3*A*).

*Proteomics.* The study of Al-Samir *et al*. did not include a proteomic analysis. On a different strain of mice, we saw no major differences in AQP1-KOs except for the loss of AQP1 (see <u>paper #2</u>^122^). In the case of Colton-null human RBCs, no data are available to address the issue of whether the cells compensated for the absence of AQP1.

*Murine data at low temperature.* We chose to work at 10 °C to achieve: (1) maximal signal fidelity, like <u>others</u>^123^ before us, and (2) minimal hemolysis (Gershfeld & Murayama, 1988).

We observed several differences vis-à-vis the murine data of Al-Samir *et al*. and ours.

1. Their absolute t_½_ at 7 °C appears to be ∼2.5-fold faster^124^ than in the present study at 10 °C (e.g., see Figure 3), even though we worked at a somewhat higher temperature. Although a difference in mouse strain could have contributed to this difference, this effect—based on our preliminary work (Zhao & Boron, 2023)—might have increased *k*_HbO_2__ by ∼35%. As noted above, their reported hemolysis was ∼3× ours. If this degree of hemolysis was accurate but unaccommodated, it would have led to an overestimate of *k*_HbO_2__. However, based on our hemolysis accommodations (e.g., see Figure 7), the effect would have been an overestimation of only ∼40%.
2. DIDS seems to have a much larger effect in their study^125^, increasing t_½_ by ∼3-fold (i.e., 1/(t_½_) falls by two-thirds). DIDS caused our *k*_HbO_2__ to fall by only about half as much, ∼31% (see <u>paper #3</u>^126^). The omission of a shape correction would have led to an overestimated % inhibition by DIDS (see Table 3*A*).
3. The data of Al-Samir *et al*. demonstrate a statistically significant effect of the AQP1-KO on t_½_—an increase of ∼26%, corresponding to a decrease of ∼21% in 1/(t_½_)^124^. We observed a decrease of only ∼9% in SC-*k*_HbO_2__ (see Table 3*B*).
4. The computed *P*_M,O_2__ of Al-Samir *et al*. is about the same as our value of ∼0.155 cm s^−1^. This is curious, inasmuch as their absolute rate constants were ∼2.5× higher at a lower temperature. They also found that DIDS reduced *P*_M,O_2__ to nearly zero vs. the ∼56% reduction in our studies (<u>paper #3</u>^126^). Thus the calculations—which for us incorporate MCV, MCHC, Ø_Major_, and *k*_HbO_2_→Hb+O_2__ as well as sequential corrections for hemolysis (in the reaction cell) and shape corrections—must have been fundamentally different.

*Human data at 10°C.* Al-Samir *et al*. report substantially slower t_½_ values for human than for mouse RBCs^124^, consistent with a greater MCV. Compared to normal RBCs, Colton-nulls exhibit a statistically significant difference in t_½_—an increase of ∼13%, corresponding to a decrease of ∼12% in 1/(t_½_). DIDS produces a significant difference in t_½_—an increase of ∼22%, corresponding to a decrease of ∼18% in 1/(t_½_).

*Human and murine data at temperatures out to 37 °C.* For the human data of Al-Samir *et al*., the trend, as temperature increases from 10 °C to 37 °C is for the t_½_ differences among (1) normal, (2) normal/DIDS, (3) Colton-null, and (4) Colton-null/DIDS—already small at 10 °C—to decrease even further^125^. The four mean t_½_ values are indistinguishable at 25 °C. At 37 °C, the authors report a significant difference (i.e., a small increase in t_½_) between normal/Ctrl and normal/DIDS, but no effect of the Colton-null mutation. A contributing factor, despite all the dots in the bar graphs, may be an N of 2 for the Colton-null blood donors.

The murine data, yield “faster” t_½_ values at higher temperature, consistent with the smaller MCV of murine RBCs. As with human RBCs, the absolute differences in t_½_ values tend to collapse with increasing temperature. Even so, at 25 °C, the authors reported a statistically significant difference in t_½_ values (a small increase) between WT/Ctrl and WT/DIDS. At 37 °C, DIDS had no effect with WTs, but did produce a statistically significant effect in AQP1-KOs—a decrease in t_½_ (i.e., “faster”). This last effect is paradoxical—DIDS, if anything, should have caused t_½_ to rise.

The ability of Al-Samir *et al*. to measure t_½_ values as short as ∼0.018 s for murine WT/Ctrl, WT/DIDS, and AQP1-KO/Ctrl cells is impressive. On the other hand, one must ask, especially in light of the aforementioned paradoxical DIDS effect, if the hardware/software are capable of reporting with fidelity events with a t_½_ values as brief as 0.018 s—and, more importantly, events that are substantially faster.

*Q_10_ estimates.* Physiological processes generally have *Q*_10_ values from 2 to 4. From the data of Al-Samir *et al*.^124^, we calculate that 1/(t_½_) increased ∼6.2-fold from 7 °C to 37 °C; this corresponds to a *Q*_10_ of ∼1.8. In a study of glucose transport in human RBCs, Lowe and Walmsley (1986) found that the velocity of ^14^C-glucose uptake increased ∼10 fold between 10 °C and 37 °C^127^ in their slower equilibrium-exchange conditions; this corresponds to a Q_10_ of ∼2.4. In their faster zero-trans experiments, the influx increased >300-fold, with a Q_10_ of nearly 8.3. Based on values of K^+^ influx on guinea-pig RBCs reported by Kimzey and Willis (1971), we calculate a *Q*_10_ of 3.0 for the “total” K^+^ influx, and 5.5 for the “active” (i.e., ouabain-sensitive) component. Based on values for the rate constant of Cl-HCO_3_ exchange in human RBCs by Chow *et al*. (1976), we calculate report a *Q*_10_ of 3.2. Studying the apparent Cl^−^ permeability mediated by AE1, Jensen *et al*. (2001) report a *Q*_10_ for human RBCs, based the data of Brahm (1977), of 7.2 below 15 °C but 3.44 above 15 °C.

O_2_ offloading from RBCs includes at least one enzyme-like step (i.e., dissociation of HbO_2_) and permeation of O_2_ through at least three membrane proteins, which one can consider as transport enzymes. Thus, in the study of Al-Samir *et al*., we would have anticipated a *Q*_10_ in the range of the other processes noted above. Given the exponential nature of *Q*_10_, the Al-Samir velocity at 37 °C—already extremely fast by RBC SF standards—would have had to have been 2.2-fold greater to reach the lower of the two *Q*_10_ values (i.e., 2.4) of Lowe and Walmsley (1986), 4.6-fold greater to reach the lower of the two *Q*_10_ values (i.e., 3.0) of Kimzey and Willis (1971), 5.6-fold greater to reach the *Q*_10_ (i.e., 3.2) of Chow *et al*. (1976), 6.7-fold greater to reach the lower of the two values (i.e., 3.4) from Jensen *et al*. (2001)—and greater yet for the other cited RBC values. If the true *Q*_10_ for O_2_ offloading from murine RBCs is even the lowest cited value of 2.4, the murine *k*_HbO_2__ signal observed by Al-Samir *et al*. at 37 °C would have been clipped at a level far below the actual *k*_HbO_2__, and they would not have been able to detect the effect of DIDS inhibition, and especially not the expected smaller effect of the AQP1-KO.

We propose that the proper control experiment for ruling out clipping of the murine *k*_HbO_2__ signal at 37 °C is to demonstrate that one can detect a substantially faster O_2_-offloading process. Absent that control, the present data do not support the conclusion of Al-Samir *et al*. (2025) that AQP1 and other DIDS-sensitive pathways, although they contribute to O_2_ permeability at 10 °C, lose that property at 37 °C.

Aside from the technical issues raised above, we ask why, during evolution, organisms would have paid the high metabolic price of retaining AQP1 and Rh_Cx_ only so they could promote O_2_ and CO_2_ fluxes at 10 °C but not at physiological temperatures? <u>Paper #2</u>^128^ shows that the combination of AQP1, RhAG, and RhD—which accounts for 55% of *P*_M,O_2__—constitutes >13% of the murine RBC-ghost PMA proteome. We do not know the identity of the pathway responsible for the missing 36% of *P*_M,O_2__ in the dKO ±pCMBS. However, if the missing protein(s) has a unitary conductance and MW similar to those of AQP1 and Rh_Cx_, we conclude that it would account for an additional >8% of the PMA proteome, for a total of ∼22%. Considering that—in the human body—RBCs constitute 70% of total cell number and ∼85% of the new cells created every day, this ∼22% of the PMA proteome represents an enormous whole-body metabolic liability. Of course, AQP1 is a powerful H_2_O channel (Preston & Agre, 1991; Preston *et al*., 1992). However, the physiological role of RBCs is not to transport H_2_O. Both AQP1 and Rh_Cx_ also conduct NH_3_ (Nakhoul *et al*., 2001; Nakhoul & Hamm, 2004). However, we suggest that an easier solution for NH_3_ carriage would have been for RBCs to increase their expression of UT-B, which also conducts NH_3_ (Geyer *et al*., 2013a). Instead, human RBCs have evolved a multicomponent ankyrin-1 complex (Vallese *et al*., 2022) that forms a metabolon that brings AE1, Rh_Cx_, and AQP1 into extremely close proximity, presumably, we suggest, to promote CO_2_ and O_2_ carriage by RBCs— the main physiological role of these cells. And when the mouse attempts to evade the cat, it is to the benefit of the mouse that its RBC gas channels—for which it has paid such a dear metabolic price—make a meaningful contribution at 37 °C.

## Acknowledgements

We thank Jean-Pierre Cartron for the gift of RhAG-KO mice. We thank him and Gerolf Gros and for extremely helpful discussions at the outset of the project. We thank Seong-Ki Lee for developing an improved approach for genotyping the RhAG-KO mice. The authors gratefully acknowledge Daniela Calvetti and Erkki Somersalo for having developed the engine of an earlier version of the CO_2_/pH reaction-diffusion model of an oocyte, which in part served as the starting point for the RBC model. We thank Thomas Radford for organizing the husbandry of the mouse colonies; Gerald Babcock for his role as laboratory manager; James W. Jacobberger and Philip G. Woost of the CWRU Flow Cytometry and Imaging Microscopy Core (FCIMC) for their assistance with flow cytometry; and Daniela Schlatzer of the CWRU Center for Proteomics and Bioinformatics for their assistance with mass spectrometry. This work was supported by Office of Naval Research (ONR) grant N00014-11-1-0889, N00014-14-1-0716, and N00014-15-1-2060; a Multidisciplinary University Research Initiative (MURI) grant N00014-16-1-2535 from the DoD, NIH grant multi-scale modeling grant 5U01GM111251 (to WFB), and NIH grant R01HL160857 (to WFB). R.O. and the modeling were supported in part by NIH grant K01-DK107787. R.R.G was supported by a fellowship grant from the ONR (N00014-12-1-0326). W.F.B. gratefully acknowledges the support of the Myers/Scarpa endowed chair.

## Statistics Tables

We use one-way ANOVA with Holm-Bonferroni post-hoc means comparison for the mean Raw *k*_HbO_2__ values, the mean hemolysis-corrected *k*_HbO_2__ (HC-*k*_HbO_2__) values and the mean shape-corrected *k*_HbO_2__ (SC-*k*_HbO_2__) values of all four mouse strains. Each table is split in two halves, with FWER set at 0.05, the upper half shows the adjusted α-value for each comparison and the lower half the *P*-value. Significant *P*-values are highlighted bold.

**Statistics Table 1:** *P*-values for Table 3*B*.

| Mouse |  | WT | AQP1-KO | RhAG-KO | dKO |
| --- | --- | --- | --- | --- | --- |
| | $P \quad \alpha$ | | $\alpha$ | $\alpha$ | $\alpha$ |
| WT |  |  | 0.0250 | 0.0125 | 0.00833 |
| AQP1-KO | $P$ | **$6.57 \times 10^{-3}$** | | 0.0500 | 0.0100 |
| RhAG-KO | $P$ | **$7.94 \times 10^{-5}$** | 0.0912 | | 0.0167 |
| dKO | $P$ | **$3.32 \times 10^{-10}$** | **$1.02 \times 10^{-6}$** | **$4.52 \times 10^{-4}$** | |

Comparison of HC- $k_{\text{HbO}_2}$
| Mouse |  | WT | AQP1-KO | RhAG-KO | dKO |
| --- | --- | --- | --- | --- | --- |
| | $P \quad \alpha$ | | $\alpha$ | $\alpha$ | $\alpha$ |
| WT |  |  | 0.0250 | 0.0125 | 0.00833 |
| AQP1-KO | $P$ | **$8.36 \times 10^{-3}$** | | 0.0500 | 0.0100 |
| RhAG-KO | $P$ | **$1.83 \times 10^{-5}$** | **$2.58 \times 10^{-2}$** | | 0.0167 |
| dKO | $P$ | **$2.72 \times 10^{-11}$** | **$4.59 \times 10^{-8}$** | **$1.56 \times 10^{-4}$** | |

Comparison of SC- $k_{\text{HbO}_2}$ (same for Fig 5B)
| Mouse |  | WT | AQP1-KO | RhAG-KO | dKO |
| --- | --- | --- | --- | --- | --- |
| | $P \quad \alpha$ | | $\alpha$ | $\alpha$ | $\alpha$ |
| WT |  |  | 0.0250 | 0.0167 | 0.00833 |
| AQP1-KO | $P$ | **$5.22 \times 10^{-5}$** | | 0.0500 | 0.0100 |
| RhAG-KO | $P$ | **$2.92 \times 10^{-7}$** | **0.0481** | | 0.0125 |
| dKO | $P$ | **$1.37 \times 10^{-15}$** | **$1.01 \times 10^{-11}$** | **$1.21 \times 10^{-8}$** | |

**Statistics Table 2:** *P*-values for Figure 6*F*.

Comparison of hemolysis
| Mouse |  | WT | AQP1-KO | RhAG-KO | dKO |
| --- | --- | --- | --- | --- | --- |
| | $P$ $\alpha$ | | $\alpha$ | $\alpha$ | $\alpha$ |
| WT |  |  | 0.0100 | 0.0500 | 0.0250 |
| AQP1-KO | $P$ | 0.199 | | 0.00833 | 0.0167 |
| RhAG-KO | $P$ | 0.761 | 0.136 | | 0.0125 |
| dKO | $P$ | 0.575 | 0.489 | 0.415 | |

**Statistics Table 3:**
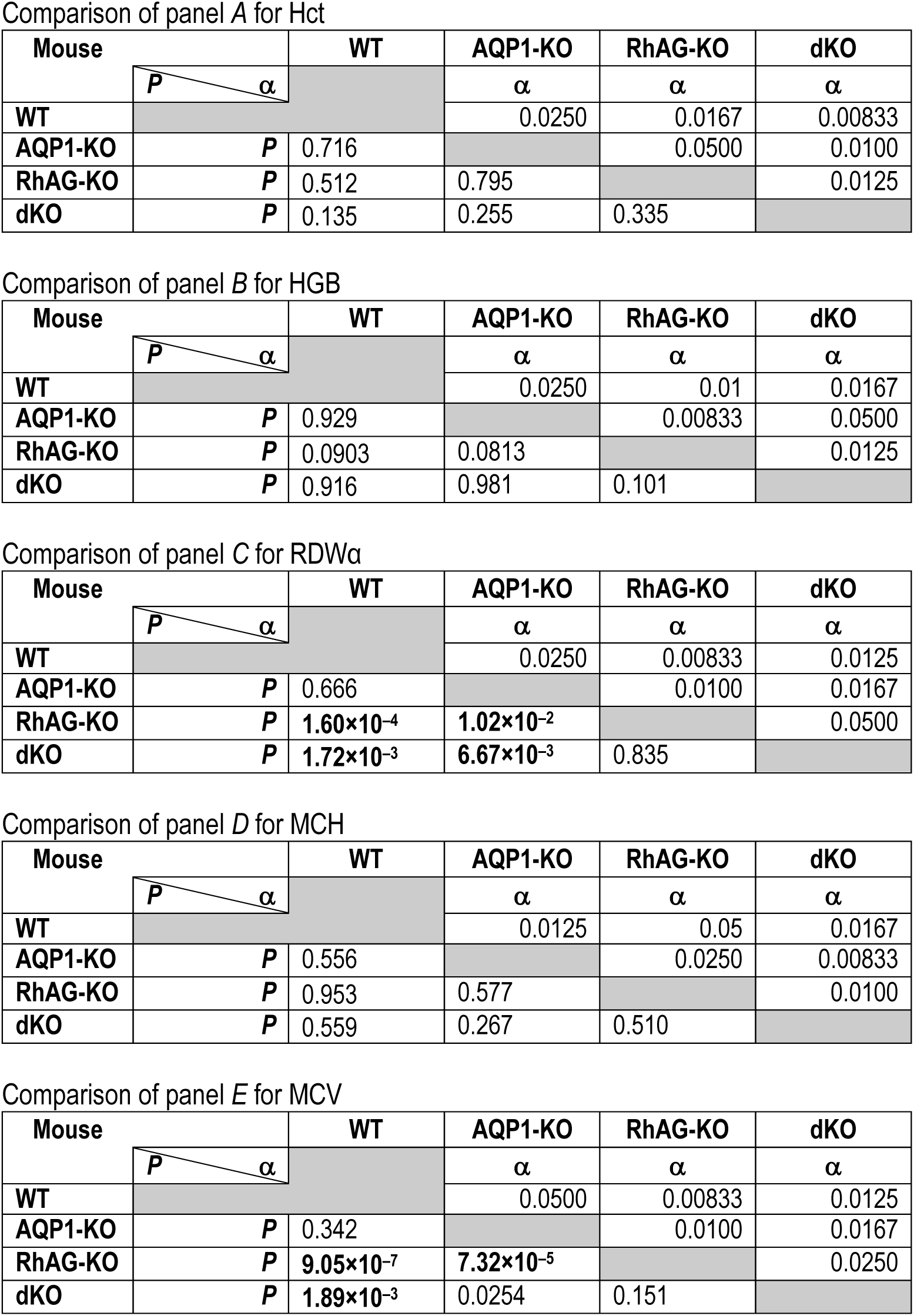
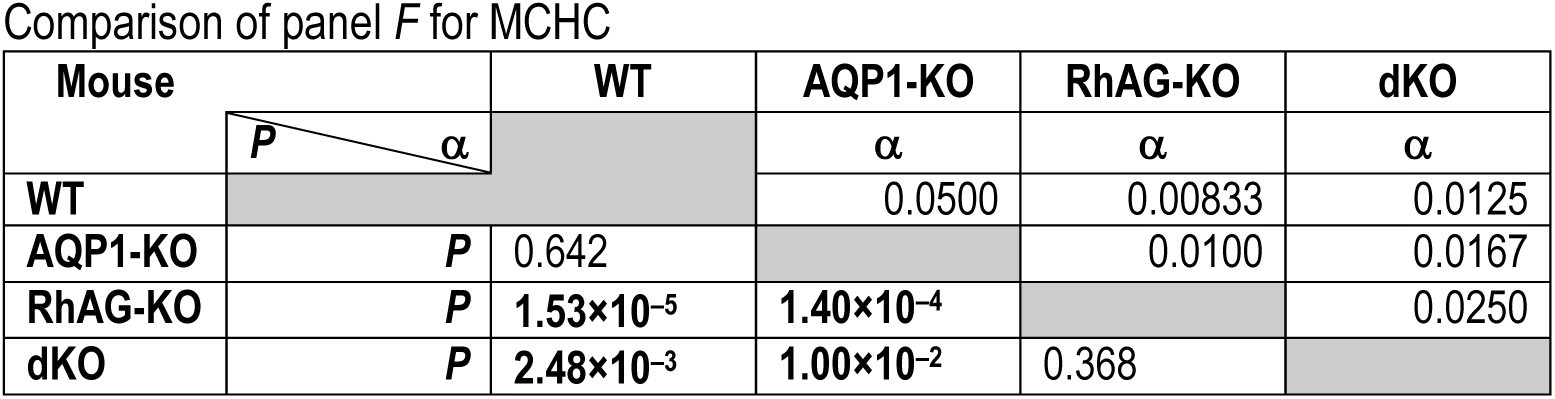
*P*-values for Figure 9.

Comparison of panel A for Hct
| Mouse |  | WT | AQP1-KO | RhAG-KO | dKO |
| --- | --- | --- | --- | --- | --- |
| | $P$ $\alpha$ | | $\alpha$ | $\alpha$ | $\alpha$ |
| WT |  |  | 0.0250 | 0.0167 | 0.00833 |
| AQP1-KO | $P$ | 0.716 | | 0.0500 | 0.0100 |
| RhAG-KO | $P$ | 0.512 | 0.795 | | 0.0125 |
| dKO | $P$ | 0.135 | 0.255 | 0.335 | |

Comparison of panel B for HGB
| Mouse |  | WT | AQP1-KO | RhAG-KO | dKO |
| --- | --- | --- | --- | --- | --- |
| | $P$ $\alpha$ | | $\alpha$ | $\alpha$ | $\alpha$ |
| WT |  |  | 0.0250 | 0.01 | 0.0167 |
| AQP1-KO | $P$ | 0.929 | | 0.00833 | 0.0500 |
| RhAG-KO | $P$ | 0.0903 | 0.0813 | | 0.0125 |
| dKO | $P$ | 0.916 | 0.981 | 0.101 | |

Comparison of panel C for RDW<sub>a</sub>
| Mouse |  | WT | AQP1-KO | RhAG-KO | dKO |
| --- | --- | --- | --- | --- | --- |
| | $P$ $\alpha$ | | $\alpha$ | $\alpha$ | $\alpha$ |
| WT |  |  | 0.0250 | 0.00833 | 0.0125 |
| AQP1-KO | $P$ | 0.666 | | 0.0100 | 0.0167 |
| RhAG-KO | $P$ | $1.60 \times 10^{-4}$ | $1.02 \times 10^{-2}$ | | 0.0500 |
| dKO | $P$ | $1.72 \times 10^{-3}$ | $6.67 \times 10^{-3}$ | 0.835 | |

Comparison of panel D for MCH
| Mouse |  | WT | AQP1-KO | RhAG-KO | dKO |
| --- | --- | --- | --- | --- | --- |
| | $P$ $\alpha$ | | $\alpha$ | $\alpha$ | $\alpha$ |
| WT |  |  | 0.0125 | 0.05 | 0.0167 |
| AQP1-KO | $P$ | 0.556 | | 0.0250 | 0.00833 |
| RhAG-KO | $P$ | 0.953 | 0.577 | | 0.0100 |
| dKO | $P$ | 0.559 | 0.267 | 0.510 | |

Comparison of panel E for MCV
| Mouse |  | WT | AQP1-KO | RhAG-KO | dKO |
| --- | --- | --- | --- | --- | --- |
| | $P$ $\alpha$ | | $\alpha$ | $\alpha$ | $\alpha$ |
| WT |  |  | 0.0500 | 0.00833 | 0.0125 |
| AQP1-KO | $P$ | 0.342 | | 0.0100 | 0.0167 |
| RhAG-KO | $P$ | $9.05 \times 10^{-7}$ | $7.32 \times 10^{-5}$ | | 0.0250 |
| dKO | $P$ | $1.89 \times 10^{-3}$ | 0.0254 | 0.151 | |

Comparison of panel *F* for MCHC
| Mouse |  | WT | AQP1-KO | RhAG-KO | dKO |
| --- | --- | --- | --- | --- | --- |
| | $P$ $\alpha$ | | $\alpha$ | $\alpha$ | $\alpha$ |
| WT |  |  | 0.0500 | 0.00833 | 0.0125 |
| AQP1-KO | $P$ | 0.642 | | 0.0100 | 0.0167 |
| RhAG-KO | $P$ | $1.53 \times 10^{-5}$ | $1.40 \times 10^{-4}$ | | 0.0250 |
| dKO | $P$ | $2.48 \times 10^{-3}$ | $1.00 \times 10^{-2}$ | 0.368 | |

**Statistics Table 4:** *P*-values for Figure 10.

| Mouse |  | WT | AQP1-KO | RhAG-KO | dKO |
| --- | --- | --- | --- | --- | --- |
| | $P$ $\alpha$ | | $\alpha$ | $\alpha$ | $\alpha$ |
| WT |  |  | 0.0167 | 0.0250 | 0.0125 |
| AQP1-KO | $P$ | 0.217 | | 0.0500 | 0.00833 |
| RhAG-KO | $P$ | 0.282 | 0.868 | | 0.0100 |
| dKO | $P$ | $8.08 \times 10^{-3}$ | $4.25 \times 10^{-4}$ | $6.31 \times 10^{-4}$ | |

## Additional Information

### Data availability statement

The data that support the findings of this study are available from the corresponding authors, W.F.B., upon reasonable request.

### Author contributions

P.Z., F.J.M., R.O., R.R.G., & W.F.B. designed the study; P.Z. performed the majority of the stopped-flow experiments, and the automated hematology and analyzed the data; R.R.G. performed preliminary stopped-flow experiments; P.Z. also made the blood smears analyzed by H.J.M.; P.Z. coordinated the microvideography experiments of red blood cells treated with inhibitors; F.J.M. performed mass spectrometry and flow cytometry experiments and analyzed the data. R.O. performed the mathematical modeling and simulations and programming; D.E.H. created a program to do data analysis on raw HbSat data to produce raw-*k*_HbO_2__ values; P.Z. and F.J.M. performed statistical analysis; P.Z., F.J.M., R.O., & W.F.B. worked on flow-chart and artist optimized the finalized version. P.Z. prepared figures; P.Z., F.J.M., R.O., & W.F.B. wrote the manuscript.

## Supporting Information

### Supporting Files

**Supporting file 1**: 1--Suppl_Tables_SF_Statistics_For_Table_4_&_Table_5.xlsx

## Footnotes

1 We use HbO_2_ to represent the generic oxygenated hemoglobin (i.e., oxyhemoglobin), measured in terms of mass concentration (e.g., g/dl). Of course, the Hb molecule is a tetramer. Under physiological conditions at sea level with the subject at rest, this tetramer enters the mammalian lung with 3 O_2_ bound on average, Hb(O_2_)_3_, and exits the lung with 4 bound on average, Hb(O_2_)_4_.

2 This manuscript was first published as a preprint: Zhao P, Moss FJ, Occhipinti R, Geyer RR, Huffman DE, Meyerson HJ & Boron W F (2025). Role of channels in the O₂ permeability of murine red blood cells. I. Stopped-flow and hematological studies. bioRxiv. https://doi.org/10.1101/2025.03.05.639948

3 See Methods>Preparation of RBCs>Processing of RBCs for stopped-flow studies. The “>” symbol separates heading levels. It is understood that the reference is to a location in the present paper unless “See” is followed by “Paper #2” or “Paper #3”.

4 See Paper #2>Methods>Preparation of RBCs>First four assays

5 See Methods>Automated hematological analyses (workflow #7-9)

6 See Paper #2>Methods>Blood smears

7 See Paper #2>Methods>Proteomic analysis

8 See Methods>Preparation of RBCs>Processing of RBCs for assays 1–4

9 See Methods>Preparation of RBCs>Collection of blood from mice

10 See Methods>Inhibitors

11 The “Pro-KIV” software supplied by Applied Photophysics assumes that the absorbances at all wavelengths decay as a perfect exponential, and fits them all simultaneously, producing a single time constant/rate. Because this approach could introduce a bias into the analysis, we developed the method described here. Nevertheless, the two approaches produce numerical results that differ only slightly, and do not affect the conclusions that we draw.

12 See Methods>Calculation of time course of HbO_2_ desaturation>Apply weighting factors

13 In some cases, HbSat fell so rapidly that we were forced to use < 5 points to the left.

14 See Methods>Calculation of raw-*k*_HbO_2__ (workflow #3)

15 See Methods>Accommodation for hemolysis (workflow #5)>Control and pCMBS-treated RBCs

16 See Paper #2>Methods>Flow cytometry>Imaging flow cytometry

17 See Methods>Preparation of RBCs>Processing of RBCs for automated hematology (workflow #7 – #9)

18 See Methods>Preparation of RBCs>Processing of RBCs for assays 1–4

19 See Paper #2>Methods>Flow cytometry>Use of inhibitors in imaging flow cytometry

20 Paper #3>Methods>Mathematical modeling and simulations>Calculation of RBC thickness based on the geometry of a torus

21 See Methods>Measurement of major cell diameter (workflow #6 & 6′)

22 See Methods>Automated hematological analyses (workflow #7 – #9)

23 See Methods>Calculation of raw-*k*_HbO_2__ (workflow #3)

24 See Methods>Preparation of RBCs>Processing of RBCs for stopped-flow studies>“osmotic lysis” in text

25 See Methods> Macroscopic Mathematical Modeling (MMM) and simulation of HbO_2_-deoxygenation time courses, yielding the “sigmoidal curves” (workflow #12 & #12′)

26 See Methods>Accommodation for nBCDs>Assignment of Provisional *P*_M,O_2__ (workflow #13)

27 See Methods>Accommodation for nBCDs>Simulations (workflow #14 & #14′)

28 See Paper #2>Methods>Flow cytometry>Imaging flow cytometry (workflow #15 & #15′)

29 We term this “macroscopic” to distinguish the present simulations at the cellular scale from <u>meso</u>scopic modeling at the supramolecular scale and molecular-dynamics (MD) simulations at the atomic scale.

30 See Paper #3>Methods>Mathematical modeling and simulations

31 See Methods>Estimation of thickness of biconcave disks (workflow #10)

32 See Paper #3>Methods>Mathematical modeling and simulations>Kinetics of the oxygen-hemoglobin reaction

33 Methods>Automated hematological analyses (workflow #7 – #9)

34 See Paper #2>Methods>Flow cytometry>Imaging flow cytometry

35 Paper #3>Methods>Mathematical modeling and simulations>Computational model

36 Paper #3>Methods>Mathematical modeling and simulations>Parameter values

37 Paper #3>Methods>Mathematical modeling and simulations>Simulation of time course of HbO_2_ deoxygenation, and estimation of *k*_HbO_2__

38 Paper #3>table 1

39 Paper #3>table 2

40 See Paper #2>Methods>Flow cytometry>Still microphotography and microvideography of living RBCs

41 See Paper #2>Methods>Flow cytometry>Imaging flow cytometry (IFC)

42 See Paper #2>Methods>Flow cytometry>Imaging flow cytometry

43 Paper #3>Results>Accommodation for non-BCDs

44 Because the intracellular diffusion distances are greater for nBCDs than for BCDs, the O_2_-offloading rates are all lower for nBCDs, so that the sigmoid curve is flatter.

45 See Methods>Stopped-flow absorbance spectroscopy (workflow #1)

46 See Methods>Calculation of time course of HbO_2_ desaturation (workflow #2)

47 See Methods>Calculation of raw -*k*_HbO_2__ (workflow #3)

48 See Results>Accommodation for Hemolysis

49 See Results>Accommodation for nBCDs

50 See Paper #3>Results>Accommodation for nBCDs

51 See Methods>Quantitation of hemolysis in SF reaction cell (workflow #4)

52 See Results>Accommodation of raw-*k*_HbO_2__ for hemolysis>DIDS-treated RBCs

53 See Results> Mathematical simulations

54 See Paper #2>Results>Proteomics

55 See Paper #2>table 10

56 See Paper #2>table 9

57 See Results> Mathematical simulations

58 See Methods>Quantitation of hemolysis in SF reaction cell (workflow #4)

59 See Results>Accommodation for hemolysis>Accommodation of raw-*k*_HbO_2__ …

60 See Methods>Quantitation of hemolysis in SF reaction cell (workflow #4)

61 See Methods>Accommodation for hemolysis (workflow #5)>DIDS-treated RBCs

62 Evaluating the above equation, as written, yields 6.11538, as opposed to 6.114. The discrepancy is due to rounding errors in the written equation.

63 Evaluating the above equation, as written, yields 1.56, as opposed to 1.57. The discrepancy is due to rounding errors in the written equation.

64 See Results>Mathematical simulations

65 See Methods>Automated hematological analyses (workflow #7 – #9)

66 See (a) Paper #2>table 2 … & … (b) ..>figure 6e

67 Paper #3>Results>Mathematical simulations…>Ø_Sphere_>Tendency of MCHC and MCV effects to cancel

68 See Methods>Automated hematological analyses (workflow #7 – #9)>Use of inhibitors in automated hematological studies

69 See Paper #2>table 6

70 See Paper #2>Results>Morphometry>Blood smears

71 See Paper #2>figure 2

72 See Paper #2>figure 3

73 See (1) Results>Hematological and related parameters>Morphometry>Living RBCs tumbling through the focal plane of an inverted microscope … & … (2) ..> Living RBCs visualized using imaging flow cytometry (IFC)

74 See Paper #2>Results>Morphometry>Imaging of living, tumbling RBCs

75 See Paper #2>figure 4

76 See Paper #2>Supporting Information

77 See Paper #2>figure 5

78 See Paper #2>Results>Morphometry>Light-scattering flow cytometry

79 See Paper #2>Results>Morphometry>Imaging flow cytometry

80 See (a) Paper #2>figure 8b*–c* … & … (b) ..>table 5

81 See Results>Accommodation for nBCDs

82 See Paper #2>table 5

83 See (a) Results>Mathematical simulations … & … (b) ..>Accommodation for nBCDs

84 See Paper #3>figure 8

85 See Paper #3>Introduction

86 See Paper #3>Methods>Mathematical modeling and simulations>Parameter values

87 See Paper #3>table 1

88 See Paper #3>table 2

89 See Paper #3>table 5

90 See Paper #3>table 4

91 See Paper #3>Results>Simulations of O_2_ efflux from a control RBC of WT mouse

92 See Methods>Accommodation for nBCDs (workflow #13, #14 & #14′, #16 – #22)

93 See Paper #3>Results>Accommodation for non-BCDs

94 See Paper #3>Discussion>Krogh-Erlang Equation

95 See Discussion>Implications of the macroscopic mathematical model> Predictions for effects on *P*_M,O_2__

96 See Discussion>Implications of the macroscopic mathematical model>Effect of channel knockouts on MCV and MCHC

97 See Paper #2>figure 5

98 See Paper #2>figure 8b*–c*

99 See Paper #2>table 5

100 See Paper #3>Results>Accommodation for non-BCDs>Linear combination of hemoglobin-deoxygenation time courses

101 See Paper #2>table 5

102 See Paper #3>table 3

103 See Paper #3>table 3, data row #3

104 See Paper #2>figure 8b*–c*

105 See (a) Paper #2>figures 2–5 and 8 … & … (b) Also see Suppl_Video_1--WT

106 See Paper #3>Results>Accommodation for non-BCDs

107 See Paper #2>tables 3 and 5

108 See Paper #2>Results>Proteomics

109 See Paper #3>figure 11

110 For our current SF experiments, we purchase several sealed glass bottles of NDT, and open them one at a time until we find one that has a powdery NDT-gas interface. The other bottles, which have a film or crust at the surface (due to NDT that has reacted with O_2_ and H_2_O), we discard. We store both new and open bottles in an evacuated desiccator.

111 See Paper #3>table 4

112 See Paper #3>figure 6

113 See Paper #3>Discussion>Calculation of the contribution of the plasma membrane to total diffusive resistance to O_2_>Potential contribution of channels to *R*_M_ and *R*_Total_.

114 See Paper #3>Results>Mathematical simulations exploring the predicted sensitivity of *k*_HbO_2__ to eight key kinetic and geometric parameter

115 See paper #2>table 4, which is a subset of the larger data set in Table 4 (in paper #1) and is restricted to the mice used in the experiments on imaging flow cytometry and the mathematical simulations. Although the numbers of observations are smaller, the trends are similar.

116 See paper #2>table 2

117 See paper #3>figure 12

118 See paper #3>figure 9

119 See paper #3> figure 5

120 See Paper #2>table 5

121 See Paper #3>Results>Accommodation for non-BCDs

122 See Paper #2>table 7

123 See Discussion>Reports on O_2_ channels>*Q*_10_ estimates

124 See Al-Samir *et al*., figure 5

125 See Al-Samir *et al*., figure 7

126 Paper #3>table 3

127 Interpolated from their figure 2

128 See Paper #2>Results>Proteomics

129 See Methods>Statistical analysis

130 See Methods>Automated hematological analyses (workflow #7 – #9)>Use of inhibitors in automated hematological studies

131 See Methods>Statistical analysis

132 See Methods>Calculation of time course of HbO_2_ desaturation (workflow #2)>Apply weighting factors

133 See Methods>Accommodation for hemolysis (workflow #5)

134 See Methods>Accommodation for nBCDs (workflow #13 – #22)

135 See Methods>Statistical analysis

136 See Methods>Preparation of RBCs>Processing of RBCs for stopped-flow studies>”osmotic lysis” in text

137 See Methods>Preparation of RBCs>Use of inhibitors in stopped-flow studies

138 See Methods>Calculation of raw-*k*_HbO_2__ (workflow #3)

139 See Methods> Quantitation of hemolysis in SF reaction cell (workflow #4)

140 See Methods>Calculation of raw-*k*_HbO_2__ (workflow #3)

141 See Methods>Preparation of RBCs>Use of inhibitors in stopped-flow studies>“millisecond-to-second DIDS exposures” in text

142 See Methods>Accommodation for hemolysis (workflow #5)>DIDS-treated RBCs

143 See (a) Methods>Calculation of time course of HbO_2_ desaturation (workflow #2) & (b) ..>Calculation of raw-*k*_HbO_2__ (workflow #3)

144 Paper #3>Methods>Mathematical modeling and simulations

